# metascreen: A modular tool for the design and analysis of drug combination screens

**DOI:** 10.1101/2022.07.10.499451

**Authors:** Robert Hanes, Pilar Ayuda-Durán, Leiv Rønneberg, Manuela Zucknick, Jorrit Enserink

## Abstract

There is a rapidly growing interest in high-throughput drug combination screening to identify synergizing drug interactions for treatment of various maladies, such as cancer and infectious disease. This creates the need for pipelines that can be used to design such screens, perform quality control on the data, and generate data files that can be analyzed by synergy-finding bioinformatics applications. metascreen is an open source, end-to-end modular tool available as an R-package for the design and analysis of drug combination screens. The tool allows for a customized build of pipelines through its modularity and provides a flexible approach to quality control and data analysis. metascreen is adaptable to various experimental requirements with an emphasis on precision medicine. It can be coupled to other R packages, such as bayesynergy, to identify synergistic and antagonistic drug interactions in cell lines or patient samples. metascreen is scalable and provides a complete solution for setting up drug sensitivity screens, read raw measurements and consolidate different datasets, perform various types of quality control, and analyze, report and visualize the results of drug sensitivity screens.

**Availability and implementation:** The R-package and technical documentation is available at https://github.com/Enserink-lab; the R source code is publicly available at https://github.com/Enserink-lab/metascreen under GNU General Public License v3.0; bayesynergy is accessible at https://github.com/ocbe-uio/bayesynergy/

Selected modules will be available through Galaxy, an open-source platform for FAIR data analysis, Norway: https://usegalaxy.no

## 1. Introduction

The advent of targeted therapy has revolutionized the treatment of many types of cancer. However, despite often eliciting a strong initial response, most targeted therapies ultimately fail due to a variety of reasons, including mutations in the molecular target, overexpression of the drug target, or activation of compensatory mechanisms (Bell and Gilan, 2020; Bergholz and Zhao, 2021; Logue and Morrison, 2012). One solution to this problem is the use of combinations of drugs (Bayat Mokhtari *et al*., 2017; Saputra *et al*., 2018; Plana *et al*., 2022). This is exemplified by clinical trials with melanoma, which is a form of cancer frequently driven by mutations that activate the Ras-Raf-MEK-ERK pathway, such as BRAF V600 mutations (Hodis *et al*., 2012). Ras pathway-driven forms of melanoma can be treated with various kinase inhibitors, including the Raf inhibitors vemurafinib and dabrafenib and the MEK inhibitors trametinib and cobimetinib (Robert *et al*., 2015; Luke and Hodi, 2013). Randomized phase III clinicals trials have demonstrated that Raf inhibitors are associated with increased progression-free survival (Chapman *et al*., 2011; Hauschild *et al*., 2012). However, acquired resistance to these single drug treatments is a major problem, and only a minority of patients showed durable responses (Hauschild *et al*., 2012; Chapman *et al*., 2011). Resistance to BRAF inhibitors can occur via multiple mechanisms, although reactivation of the MAPK pathway is a common theme (Solit and Rosen, 2011), which can be partially overcome by combining BRAF inhibitors with MEK inhibitors (Flaherty *et al*., 2012). Additional combinations of targeted therapy and, more recently, with immunotherapy have been identified that can overcome resistance (Luke *et al*., 2017). Similar effects have been observed for a wide variety of cancers, including AML, lung cancer and breast cancer (Fisusi and Akala, 2019; Latif *et al*., 2021; Yuan *et al*., 2019), and combination treatment clearly has the potential to significantly improve survival rates for cancer patients. However, given the sheer number of targeted therapies, identifying synergistic drug combinations is a major challenge, and several obstacles still need to be overcome.

One such obstacle is the lack of an integrated bioinformatics workflow that integrates the design and execution of drug combination screens. Such a workflow should also include quality control measures to identify and reduce technical variability, which is a persistent problem that limits the reproducibility of high-throughput drug screens (Niepel *et al*., 2019; Larsson *et al*., 2020; Hatzis *et al*., 2014). Lack of reproducibility is a well-documented problem in the screening of new drugs for the treatment of cancer. However, including strict quality control measures during the early stages of preclinical development can contribute to reducing the high attrition rates associated with cancer drug sensitivity screens (Larsson *et al*., 2020). The bioinformatics workflow should also include a data output step that produces data files that can be visualized and analyzed across multiple platforms.

Here we present metascreen, which provides a comprehensive end-to-end solution that integrates design, execution, quality control and data analysis of large-scale drug combination screens.

## 2. Materials and methods

The metascreen main modules and functions.

**Figure 1.**
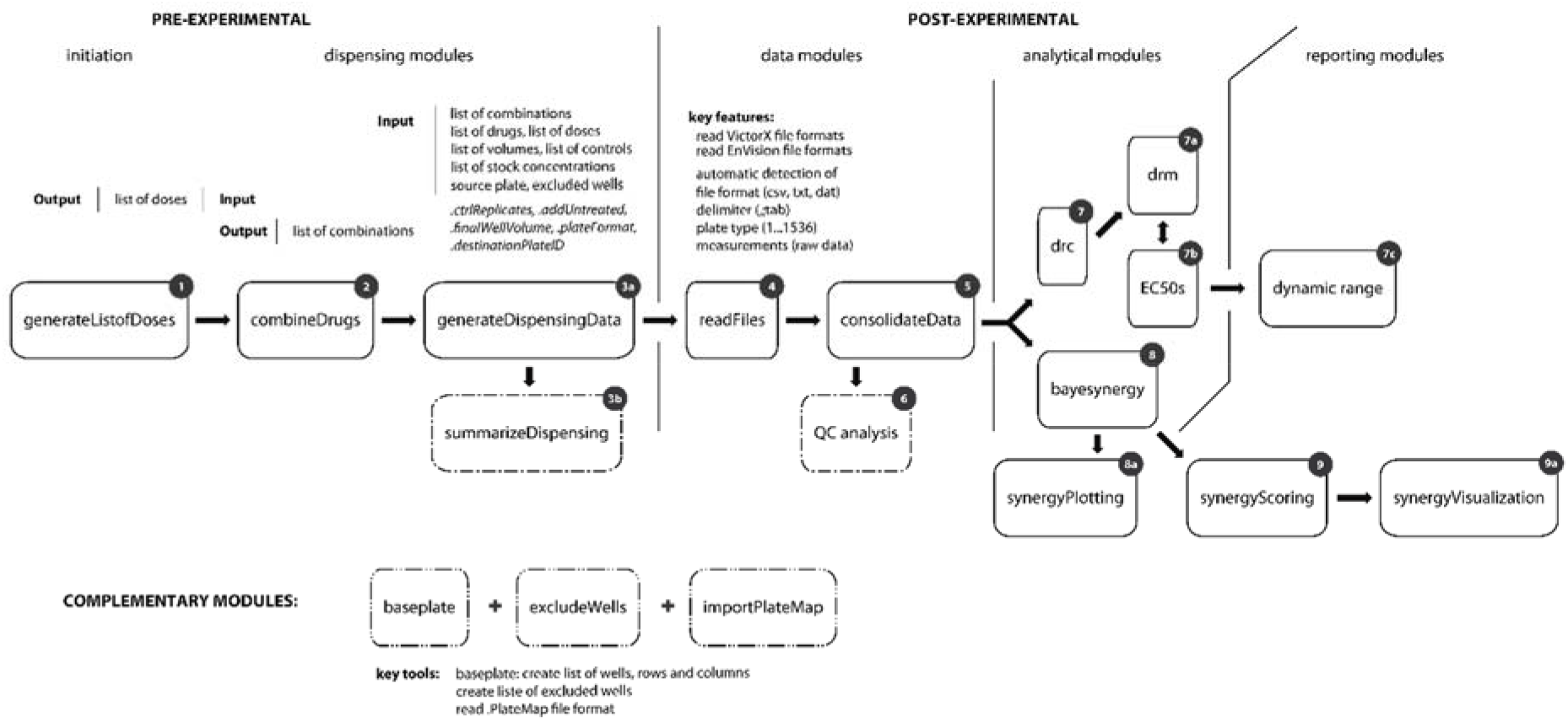
A pipeline with the main modules and functions of metascreen

### 2.1. pre-experimental: setting-up the experiment and generating dispensing data

#### 2.1.1. generating a list of drug concentrations (*doses*)

To be able to designate combinations of drugs in a drug combination screen, an inventory list of drug names and concentrations needs to be provided. This list can be provided in a wide data format, which is readable for humans and intuitively used by many users, in form of a comma-separated values (CSV) file.

The function of this module is to import the list from a CSV file and convert it from a wide data format into a machine-readable long format by grouping multiple numbers of concentrations into single columns. It also allows the user to exclude any non-essential columns. The essential columns are labeled as *Drug, Dose* and *Unit*, respectively. The provided list of drug concentrations needs to contain columns with these three terms. The input data can be provided in a wide format with multiple columns of concentrations. Alternatively, the columns containing the drug concentrations can be selected through the argument `.doseIdentifier`, allowing the function to identify all columns with the stated term as essential columns.

The function can be called

~~~
generateListofDoses(listofDoses, .doseIdentifier=“dose”, .dropCol = TRUE)
~~~

With the argument `.dropCol`, non-essential columns can be excluded.

The output from the function above is a list of drug concentrations in a long data-format, which can then be used to combine drugs at individual drug concentrations with each other.

#### 2.1.2. generating a list of combinations

This module will generate a list of combinations from the list of drugs and drug concentrations provided. This list will then be used to generate the dispensing layout and files.

~~~
combineDrugs(listofDoses, .combineDoses, .noReplicates = 1,
.drugRepAttrib = c(“all”, “single”))
~~~

In addition, it can be specified, which of the drug concentrations should be combined and how many replicates of either single or all drug treatments should be generated.

With the argument `.combineDoses`, drugs can be combined at individual drug concentrations, or at a certain drug concentration range, in which a low index corresponds with a lower drug concentration. i.e. a value of 1 selects for the first, and consequently, for the lowest drug concentration among all the drug concentrations of a drug. A range of drug concentrations can be selected as well as a number of individual drug concentrations. With the argument `.noReplicates` the number of replicates can be specified for either single drug treatments or for all treatments, and both single drugs and drug combinations using the argument `.drugRepAttrib`.

The output from the function is essential for the generation of the dispensing files.

#### 2.1.3. The exclusion of individual wells from an experimental set-up

A complementary module of this package allows for generation of a list of individual wells or series of wells that should be excluded from the experimental set-up, for instance to exclude certain wells on multi-well microplates that are sensitive to evaporation. This function supports the most common microplate formats, ranging from 6 to 1536 wells. It allows for selection of a combination of designated rows, columns and individual wells for exclusion. Rows and columns can be specified by their names, which are designated letters for rows (A, B, C, ..) and designated numbers for columns (1, 2, 3, ..). Alternatively, in addition to entire rows and columns, individual wells (A1, B2, C3, ..) can be marked for exclusion. The nomenclature of wells follows the ANSI guidelines set by the Society for Laboratory Automation and Screening (Society for Laboratory Automation and Screening). In addition, the function offers the convenient possibility to exclude the outer wells of any given plate format.

~~~
excludeWells(plateType, wells = NULL, outer.wells = FALSE)
~~~

The function will generate a list of well coordinates that can be used to specifically exclude wells from the dispensing procedure. The function can also be used in a more general manner to generate any sequence of wells for various analytical or experimental tasks, such as generating a list of wells of interest that can be used to specifically select or filter for individual wells, rather than for exclusion.

#### 2.1.4. The import of plate maps

This pipeline offers a module that allows the user to import plate maps from various sources. These plate maps can be directly imported from a comma separated values (.csv) file, or by utilizing a third-party software, such as the IncuCyte® Plate Map Editor. Third-party applications and tools usually work with proprietary file formats, which are not directly usable with R.

The function importPlateMap allows the user to import IncuCyte® PlateMap files, which are in a proprietary file format used by the IncuCyte® Plate Map Editor. This allows users to design source plates using the IncuCyte® Plate Map Editor and import the plate map into their pipeline. This module can also be used outside the scope of drug sensitivity screens, e.g. for projects that utilize the IncuCyte® live cell imaging system.

~~~
importPlateMap(importFile, .fileFormat, .plateMapGrp = list(type, name,
group), .sourcePlateConv = FALSE)
~~~

The plate maps can be imported either from a single file or from multiple files. It is recommended to specify the file format of the file to import with the argument`.fileFormat`, especially, if the folder from which the files are imported contain files with different file formats. Alternatively, the function is able to detect any of the supported file formats automatically and import them. With the argument `.plateMapGrp` individual groups of cells, drugs or conditions can be defined. The argument `.sourcePlateConv` allows for converting the file format of the plate map to only the essential columns required for the generation of the dispensing files by removing all non-essential data, such as ambiguous columns or empty rows.

#### 2.1.5. generating dispensing files

It is crucial to the overall design and execution of the assay to generate a dispensing file that contains instructions regarding which drugs should be dispensed at which concentration from source plates holding drug stocks onto destination plates that will be used for the actual assay. With this module it is possible to generate a plate layout in which the individual single drug treatments and drug combinations are distributed across a number of plates. It takes into account the controls and excluded wells that have been previously defined. It is also possible to generate dispensing files in a predefined format specific for (various) dispensing robots. At the moment the LABCYTE Echo 550 acoustic dispenser is supported (more dispensers can be integrated at a later time point). Due to the technical properties of the Echo acoustic dispenser, usually only volumes of 2.5 nl increments can be dispensed, which needs to be taken into account when designing the source plate with the appropriate stock concentrations of drugs. This technical limitation can be accommodated by providing a list of volumes to be dispensed from the source plate to obtain the desired final drug concentration of a given treatment, when generating the dispensing files. The list of the volumes serves consequently as a reference between the stock concentration on the source plate and the final concentrations on the destination plate. This allows the transfer of a drug from a single well on the source plate to the destination plate at any given volume, instructing the robot which well on the source plate and which well on the destination plate should be used. Future work will allow a more customizable approach, in which individual dilutions can be dispensed from individual or multiple wells.

Furthermore, a list of combinations, a list of controls and a list of excluded wells is required to generate a dispensing file. Depending on the experimental set-up, it is possible to exclude certain wells from being targeted to avoid technical and experimental variation, such as the edge effect on most microtiter plates due to a higher evaporation on the outer wells. In addition, a list of wells or a plate map of the source plate is required. The source plate contains the drugs in their stock concentrations to be dispensed in dilutions onto the destination plate. This module is able to read the proprietary plate map file format from the Incucyte® ZOOM Live-Cell Analysis System from Essen BioScience. Alternatively, a reference list of wells on the source plate can be provided separately.

The final dispensing layout on the destination plate can arrange sample and drug treatments in sequential order, or as recommended, randomized across a set of plates. In addition, the number of sets can be specified in case several drug sensitivity screens will be carried out with multiple patient samples or cell lines. For each dispensing and drug combination a unique identifier is given, which is linked to a unique coordinate on the destination pate. Additional features of this module include the visualization of the final plate layout. In cases where the dispensing volume needs to be monitored in order to avoid critical levels and the depletion of drugs, the module is able to provide feedback on the dispensed volume per set as well as the total volume for a given source plate.

~~~
generateDispensingData(listofCombinations, listofDrugs, listofDoses,
listofVolumes, listofCtrls, listofStockConcentrations, sourcePlate,
listofExWells, .ctrlReplicates, .addUntreated, .finalWellVolume,
.plateFormat, .destinationPlateID, .randomizeDispensing = TRUE,
.probeDispensing = FALSE)
~~~

The function requires a list of combinations, a list of drugs, drug concentrations, volumes, as well as a list of stock concentrations and controls. Furthermore, it requires a source plate and optionally a list of excluded wells. It allows for specifying the number of replicates for control treatments, the designation of an individual plate ID or barcode, as well as the possibility for full randomization of the drug treatments across all plates. An additional feature of this module is to probe the final dispensing based on the provided input data. This will simulate the dispensing procedure by performing a dry-run and return statistics regarding the number of unique dispensions for single and combination drug treatments, controls and the number of plates that the final experimental set-up will require. In order to *save* the dispensing data, *print* a summary, *plot* or *export* the dispensing files, these functions can be used, which extend the generic functions *print, summary, plot* and *save* for objects of class ‘dispensingData’:

**print**, to print and extract the dispensing data set

**summary**, to show a dispensing summary, with the number of plates, drug treatments and much more.

**plot**, to plot the dispensing layout for each plate.

**save**, to save and export the dispensing data to individual files for each set.

### 2.2. post-experimental: reading raw data and consolidating datasets

metascreen offers modules to read the raw data and consolidate datasets prior to any analysis. These modules are utilized post experimentally and include various modules for data handling, analysis and data reporting.

#### 2.2.1. reading raw data files

With this module the raw measurements are read from multiple files and formatted for data processing, which is necessary for downstream analysis. At the moment different output formats and files from the *PerkinElmer EnVision* and *Victor X Multimode Microplate Reader* are supported. Due to its modular build, this module can be extended with file formats from additional plate reading devices at any given time.

Reading raw data:

~~~
readFiles(.readfrom = “data/raw”, .fileformat = c(“.csv”, “.txt”), .format
= “EnVision”)
~~~

The output only contains the raw measurements in reference to the plate and well number, without being associated to any treatments or samples. This step is being done by another module. This function only reads data from microplate readers and can be used for a number of different studies.

#### 2.2.2. consolidate Data

Before the raw measurements can be used for downstream analysis of drug sensitivities, a final reference data set needs to be built. This can be achieved by consolidating the raw measurements with the generated dispensing data that was used to run the experiments. This module associates the raw measurements with the dispensing data. Its main purpose is to build a complete data set with all the raw measurements in reference to the individual drug treatments on all the plates, which is assigned a unique combination identifier and contains the complete set of meta data, such as plate number and plate barcode, drug identifiers (CAS no.), sample type, source and destination wells, transfer volumes etc. The output from this module ensembles the first consolidated full data set formatted for downstream processing and analysis. The integrity of the data set is maintained and is not subject to changes during any of the downstream processes, but is rather used as a master dataset for different derivatives.

Before this can be done, it is necessary to import a barcode reference list with the names of the samples used in the drug screen and by associating them to the corresponding plate id and set. Once this information is provided, the raw measurements can be consolidated with the dispensing data, using the function below:

~~~
consolidateData(dispensingData, rawMeasurements, .barcodeReference)
~~~

Since this function is dependent on datasets previously generated, the data needs to be of a specific class to ensure data integrity. The dispensingData needs to an object of class *‘dispensingData’*, while the rawMeasurements an object of class *‘rawMeasurements’*. The `.barcodeReference` is a list of samples for each set of plates.

#### 2.2.3. QC (quality control)

The package comes with a set of quality control (qc) tools that are imperative for the quality assurance of a drug sensitivity screen. This module provides a function that offers a set of quality assessments by looking at the variance of individual controls, reporting the Z’-factor (Zhang *et al*., 1999) between the positive and negative controls, as well as looking at the signal of empty and untreated wells.

~~~
qc(consolidatedData, .ctrls, .qcMethod)
~~~

One of the quality methods of the function *qc* is used to assess the quality of a drug screen by looking at the variance and signal distribution between individual controls. The argument

`.qcMethod` is used to select between individual quality assessments, or alternatively run multiple or all quality methods at once. At the moment the following qc methods are available:

*variance* : assessing the variance between individual controls both, across all plates, as well as by individual plate *emptywells* : assessing the signal of empty and untreated wells, this will also include any excluded wells *firstcolumn* : assessing the signal of wells in the first column of each plate *zprime* : assessing the distribution between the positive and negative controls.

#### 2.2.4. data processing

With this module the data is prepared for various tasks in the downstream analysis of the drug sensitivity screen. It carries out three essential steps in data processing: normalizing, splitting and assembling (re-formatting) the data.

~~~
processData(consolidatedData, .ctrls = list(positive, negative))
~~~

The data processing unit takes advantage of the consolidated raw data and creates the first processed and modified data set for downstream analysis. This function processes the dataset by normalizing the raw measurements to the positive and negative control. It subsequently splits the data into individual datasets, one for the controls, single drug treatments and combination treatments. These datasets are formatted in two ways, as a list of dataframes using a tabular structure or as a list of dose-response matrices. This is done to ensure adaptability of the dataset to different analysis tools with unique data format requirements.

#### 2.2.5. dose-response analysis

The dose-response module allows the assessment of single drug dose-responses. It gives an overall overview of the performance of individual drugs. runDRM is a function that runs the dose-response model on the single drug treatments. It performs curve fitting using a four-parameter log-logistic function (LL.4), and estimates the EC10, EC50 and EC90. The function also plots the single drug curves by viability and inhibition for each single drug.

~~~
runDRM(processedData, .saveto, .plot)
~~~

The dose-response modeling is performed using the function **drm** from the R-package *drc* (Ritz *et al*., 2015).

#### 2.2.6. custom plotting

This module generates a set of custom plots for the visualization of the drug-dose responses. customPlotting is a function that generates a set of plots that provide an overview of the dose-response not only between drugs, but also between individual samples. Furthermore, it plots the dose-response matrix for drug combinations along with the dose-response curves for each individual drug pair.

Input: The function requires the processed data as an object of class *‘processedData’*, along with the dose response data as an object of class *‘doseRespModel’*.

~~~
customPlotting(processedData, doseRespModel, .saveto)
~~~

Output: A set of image/png files saved to a specified location on disk.

#### 2.2.7. dynamic range

This module offers the assessment of the dynamic drug-activity range for individual drug-dose responses.

~~~
dynamicRange(data, .saveto)
~~~

The dynamic range is used to provide an assessment of the drug activity range across the chosen range of drug concentrations. This is an important indicator, particularly for drug combination screens, because it determines whether the selected drug concentration ranges used for the combination treatments do indeed fall within the activity ranges of the drugs. The dynamic range is considered the dose-response range between the ED10 and ED90. This function will indicate the expected and the observed drug activity range for each drug and sample. It will generate plots based on the fitted dose-response models as well as unfitted curves.

#### 2.2.8. Synergy Analysis with bayesynergy

This particular module is primarily used for the estimation of synergies from the interaction between two drugs across their drug concentration range based on a bayesian semi-parametric model. It calculates synergies using the R-package bayesynergy (Rønneberg *et al*., 2021). The output of this module is directly streamlined from the output of bayesynergy as a list containing the posterior distribution summaries of the volume under the surface (VUS), which summarize efficacy and interaction effects of the drug combinations, the EC50s based on the monotherapy curves, additional statistics of the fitted bayesyenrgy model and additional quality parameters, including a measure of synergy (bayesfactor). Optionally, the output and bayesynergy plots can be saved to a user specified location.

~~~
bayesynergy(processedData, .saveoutput, .plot, .saveto)
~~~

The function bayesynergy uses the processed and normalized data set ‘*processedData*’ and returns a class S3 object ‘*bayesdata*’ of type list.

#### 2.2.9. Synergy scoring

With this module the synergies are ranked based on the median synergistic VUS divided by the mean absolute deviation. The antagonistic score is derived from the antagonistic VUS. The synergy score is not the product of the mean synergy and antagonism, but rather decoupled from the antagonism. Even though the primary focus of this screen is to find synergies, the importance of the antagonistic response of a drug combination should not be overlooked, since it might have clinical relevance. In cases where the estimation is accompanied with a high uncertainty in its prediction the corresponding drug combination and synergy score is flagged yet still retained. The ranked synergy scores can be saved together with the antagonistic scores and several other statistical information along with a number of quality control parameters as a csv file.

The synergy scores can furthermore be visualized in a number of different plots.

~~~
synergyScoring(bayesdata, .saveoutput, .plot, .saveto)
~~~

### 2.3. Documentation and extended development

Both user documentation and technical documentation are available for the use and implementation of the R-package. The R-package is available as an open source project under GPL (GNU General Public License) v3.0 and open for further developmental contribution. metascreen can be extended to integrate additional experimental methods and will be subject to further development as a base for feature engineering.

### 2.4. Installation

metascreen can be installed from a github repository using the R-package *devtools*.

~~~
install.packages(‘devtools’)
library(devtools)
# Install R-package from github
devtools::install_github(‘Enserink-lab/metascreen’, build = TRUE,
build_opts = c(“--no-resave-data”, “--no-build-vignettes”))
# Load ‘metascreen’ package
library(metascreen)
~~~

### 2.5. Prerequisite

In order to install and run *metascreen*, the following dependencies need to be installed prior the installation of metascreen:

bayesynergy: https://github.com/ocbe-uio/bayesynergy/

### 2.6. Real case application

The R-package was implemented and validated in a real case scenario in a large-scale drug combination screen with more than 23 melanoma cell lines, 61 drugs in pairs on a 1536-well plate-format. The implementation of modules into a fully functional pipeline and its results are shown in the Supplement. The data used in the case study is available at https://github.com/Enserink-lab/metascreen/tree/main/data

### 2.7. Discussion

metascreen makes it possible to adapt the analysis with a customizable workflow on the basis of its modularity. While coupling multiple individual modules, metascreen has a strong emphasis on retaining data integrity throughout the pipeline to ensure data quality and consistency. In order to maintain data integrity, metascreen carries through the consolidation of data from different sources, and performs cleaning and formatting of data into various pre-set formats for different downstream process that are dependent on different data formats. metascreen necessitates the predefinition of standards and requirements for experimental quality, which significantly reduces variability and increases reproducibility. The quality of the data has to be ensured prior to any analytical proceedings. metascreen offers an individual module for examining data quality, providing an analytical and visual feedback prior to the analysis.

The metascreen package was also designed to adapt to technological advances with the ability to up-scale an experimental design whenever necessary. However, up-scaling leads to substantial practical and technical challenges, one of them being the processing of big data and the associated computational challenges. The increase in throughput leading to lower volumes leading to an increase in susceptibility to experimental and technical variation, making it a challenge to separate the signal from the noise of technical and biological variation. With this in mind, metascreen was developed with a strong focus on quality assurance and data integrity.

metascreen was also designed to accommodate data integration from various sources, considering that at various stages of the experiment, data from different machines and devices will be generated, all having different formats. Another important objective in mind was the visualization of data throughout the analytical pipeline. At any essential stage of the analysis, data can be reported back in a visual context for easier interpretation of the results and better decision-making for consecutive experiments. metascreen has profound data analytics and allows the exploration of data by means of visual representation. metascreen’s potential can be integrated throughout different stages of an experiment, from the initiation and design of an experiment, to fundamental stages of data acquisition and processing, to experimental analysis, as well as post-experimental evaluation.

metascreen guides the user through various stages by offering a number of tools to get started. It offers tools to convert different data formats, import data from different sources, generate machine readable data files, such as dispensing files, that are crucial for the execution of the experiment, design the layout of the experiment, test and validate the experimental design prior the run of an experiment. Another important feature of metascreen is that it offers features that allow to customize the design of an experiment by choosing a number of different plate formats, allowing to up-or downscale an experiment, work with a wide range of drugs, volumes and drug concentrations, how many replicates to use, the use of different experimental controls, the possibility of randomization and exclusion of wells and much more.

Finally, the output of metascreen can be used to connect to a variety of modules for the post-experimental phase. It is able to read raw data from selective sources and consolidate different datasets prior any normalization and analytical processing. The analytical modules are separated in modeling the dose response of single drug treatments and the combinatory drug effects, respectively. Both modules provide feedback on the overall quality of the modeling and offer the visualization of a vast amount of data in a representative and meaningful way.

## 3. Conclusion

We have designed an integrated solution for high-throughput drug combination screening, which should be of use to a wide variety of researchers. The R-package proved to be of substantial significance for the successful completion of our drug combination screen made of a multitude of samples. A major strength of this tool was the ability to create a customized pipeline due to its modularity and the ability to scale the experimental design to the best cost-to-benefit ratio. Individual modules, such as the quality control assessment, showed to be of tremendous value for the overall success of the drug screen. The visualization and reporting of individual modules allowed the interpretation of results in real-time at any given step of the analytical process. We believe that metascreen is an important resource for the field.

## Acknowledgements

We like to thank members of the Enserink lab for helpful discussions.

## Funding disclosure

This research was funded by grants 2018012 and 2019096 from Helse Sør-Øst. This research was funded by grant 182524 from The Norwegian Cancer Society. This research was funded by grants 294916 and 262652 from The Research Council of Norway. The project has also received funding from the European Union’s Horizon 2020 research and innovation programme under grant agreement No. 847912 (RESCUER).

## Notes

https://github.com/Enserink-lab/metascreen

### Competing Interest Statement

The authors have declared no competing interest.

