## Supplementary for "metascreen: A modular tool for the design and analysis of drug combination screens"

**Case study**

In a case study, the pipeline was successfully employed in a large-scale drug combination screen with more than 23 melanoma cell lines, 61 drugs in pairs on a 1536-well plate-format (will be reported elsewhere).

**(a) pre-experimental:**

For the initial set-up of the drug screen, a number of files had to be provided.

Before a list of combinations could be generated, it was necessary to provide a *list of drugs* with a unique drug number, drug name, the doses in sequential order and the unit of the dose.

| Number | Drug | 6th Dose | 5th Dose | 4th Dose | 3rd Dose | 2nd Dose | 1st Dose | Unit |
|---|---|---|---|---|---|---|---|---|
| 2 | Venetoclax | 60 | 30 | 7,5 | 1,875 | 0,46875 | 0,1171875 | µM |
| 13 | Methotrexate | 0,3 | 0,15 | 0,0375 | 0,009375 | 0,00234375 | 0,0005859375 | µM |
| 23 | Cladribine | 10 | 5 | 1 | 0,2 | 0,04 | 0,008 | µM |
| 24 | Everolimus | 20 | 10 | 1 | 0,1 | 0,01 | 0,001 | µM |
| 28 | Cytarabine | 10 | 5 | 1 | 0,2 | 0,04 | 0,008 | µM |

Each dose for a given drug is provided in a wide format. There are not restrictions to the number of doses that can be used. The labels for each dose are free to choose, but must contain the word »*dose*« or have the identifier for the doses declared through the respective argument. Each drug is provided as a single line.

In this very first step, the *list of doses* was converted from a wide- to a long-format using the function `generateListofDoses`. The function checks for conformity of the list and allows any non-essential columns to be dropped.

```
generateListofDoses(listofDoses, .doseIdentifier = "dose", .dropCol = TRUE)
```

Next, the *list of doses* was used to generate a *list of combinations* through the function `combineDrugs`.

```
combineDrugs(listofDoses, .combineDoses=c(2:5), .noReplicates = 3,
.drugRepAttrib = "single")
```

| Drug.1 | Dose.1 | Unit.1 | Drug.2 | Dose.2 | Unit.2 |
|---|---|---|---|---|---|
| Venetoclax | 30 | µM | Cytarabine | 5 | µM |
| Methotrexate | 0.15 | µM | Venetoclax | 30 | µM |
| Cytarabine | 10 | µM | Cytarabine | 10 | µM |
| AG-221 (Enasidenib) | 50 | µM | Ponatinib | 2 | µM |
| Methotrexate | 0.15 | µM | AG-221 (Enasidenib) | 12.5 | µM |

This function uses the *list of doses* generated in the previous step and allows to select individual doses to be combined. In this study we used a dose range of total 6 concentrations for each drug, but combined only the four innermost doses, meaning the second lowest up to the fifth dose. This was achieved by providing the dose range through the argument `.combineDoses = c(2:5)`.

In addition we chose to use technical triplicates for the single drug treatments through the arguments `.noReplicates = 3` and `.drugRepAttrib = “single”`.

After we obtained a list of combinations, we had to import a set of additional files, which included a *list of drugs*, *volumes*, *controls* and *stock concentrations*, respectively.

The *list of drugs* contained a unique id or number, the drug name and its CAS number.

| NUMBER | NAME | CAS_NUMBER |
|---|---|---|
| 2 | Venetoclax | 1257044-40-8 |
| 13 | Methotrexate | 1959-05-02 |
| 23 | Cladribine | 4291-63-8 |
| 24 | Everolimus | 159351-69-6 |
| 28 | Cytarabine | 147-94-4 |

The *list of volumes* contains the volumes at which each dose is being dispensed from the source plate to the destination plate. The list resembles and follows the same formatting of the *list of doses* that was provided. Due to the limitations of the acoustic liquid dispenser all volumes were dispensed at increments of 2.5 nl.

| Number | Drug | Vol 6th Dose | Vol 5th Dose | Vol 4th Dose | Vol 3rd Dose | Vol 2nd Dose | Vol 1st Dose | Unit |
|---|---|---|---|---|---|---|---|---|
| 2 | Venetoclax | 10 | 5 | 5 | 5 | 5 | 5 | nl |
| 13 | Methotrexate | 10 | 5 | 5 | 5 | 5 | 5 | nl |
| 23 | Cladribine | 10 | 5 | 5 | 5 | 5 | 5 | nl |
| 24 | Everolimus | 10 | 5 | 5 | 5 | 5 | 5 | nl |
| 28 | Cytarabine | 10 | 5 | 5 | 5 | 5 | 5 | nl |

The *list of stock concentrations* lists the stock concentrations of the drugs used.

| NUMBER | DRUG | CONCENTRATION | UNIT |
|---|---|---|---|
| 2 | Venetoclax | 100 | mM |
| 13 | Methotrexate | 200 | mM |
| 23 | Cladribine | 100 | mM |
| 24 | Everolimus | 30 | mM |
| 28 | Cytarabine | 200 | mM |

In addition, our experimental design required the exclusion of outer wells. A list of excluded wells was generated with the corresponding function.

```
excludeWells(1536, outer.wells=TRUE)
```

To be able to generate instructions for dispensing by the means of a dispensing file, we had to import a plate map of the source plate, which contained the drug names and drug concentrations along with the location on the source plate from which each drug and dose was dispensed. Dispensing files are instructions for pipetting and dispensing robots on how to dispense individual drug treatments. Specifically, they provide the volume to pipette, from what well on the source plate, to the well location on the destination plate. Source plates are usually used as a library of drugs at given concentrations from which drug treatments, such as drug combinations, are dispensed. Those drug treatments are usually carried out on one or more destination plates on which experiments are being conducted.

A schematic representation of the source plate is shown in Supplementary Figure 1 and 2 with each dose used in the drug sensitivity screen located in an individual well. The controls used in the screen were on a separate source plate, containing the positive (BzCl) and negative (DMSO) control in multiple wells with enough volume for a given number of experimental set of plates.

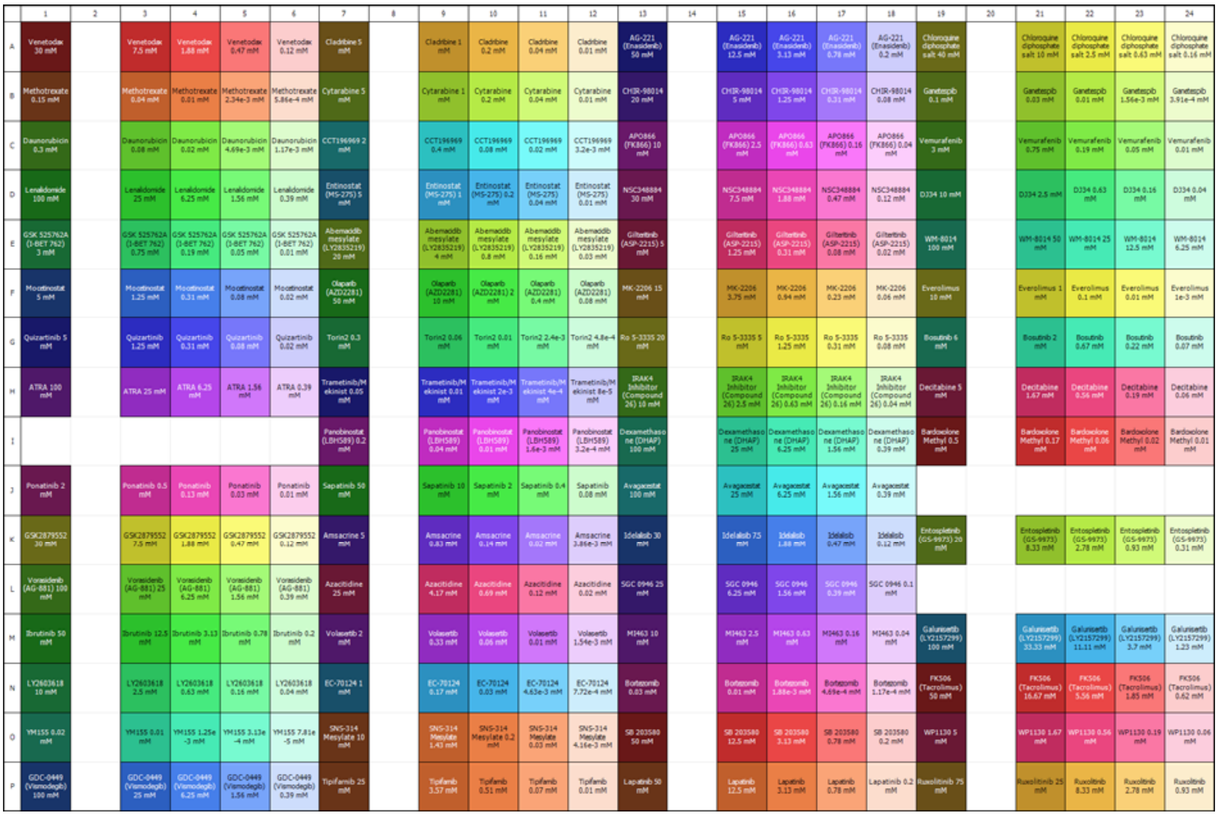

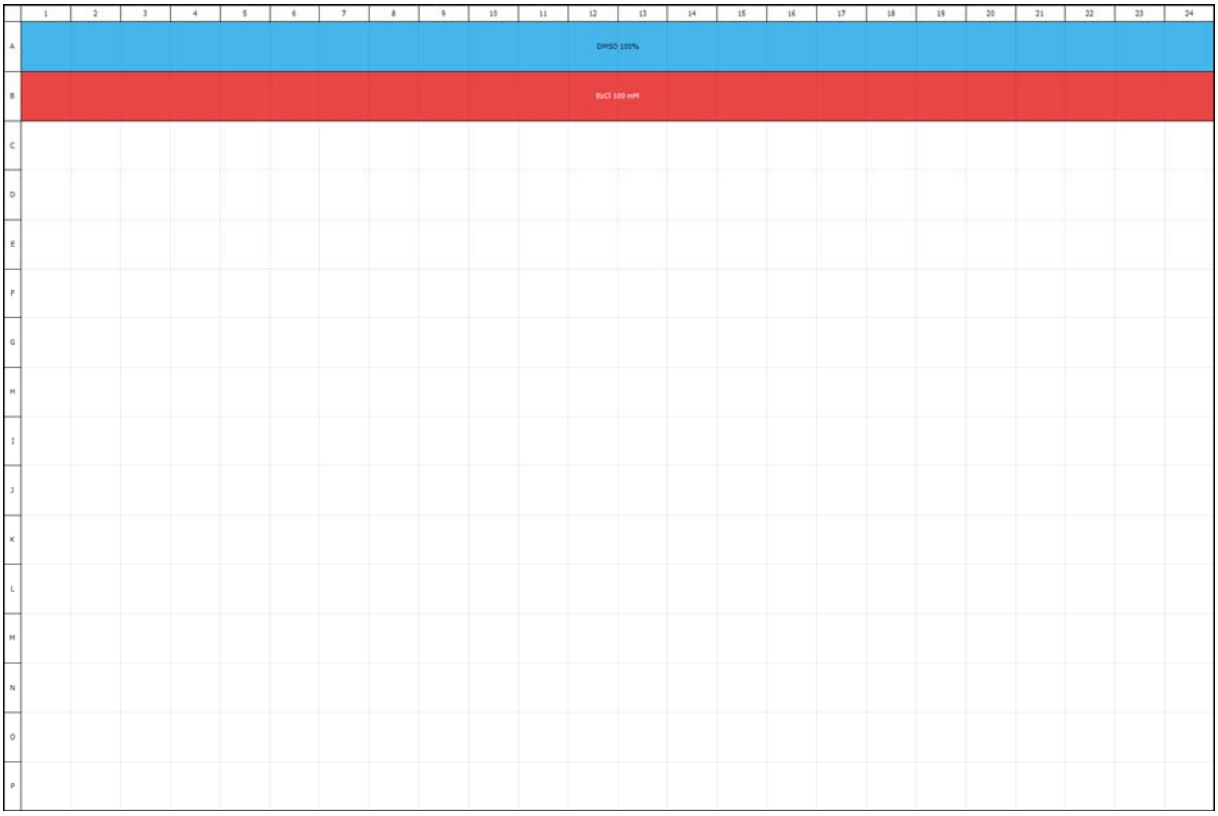

Supplementary Figure 1 and 2. Source plate with the stock concentrations of drugs

The source plates were designed using the IncuCyte® Plate Map Editor and exported in the proprietary IncuCyte® .PlateMap file format. They were imported into the pipeline with the help of a supplementary module, using the corresponding function, specifically designed for that purpose.

```
importPlateMap(libDirectory, .fileFormat = ".PlateMap", .sourcePlateConv =
TRUE)
```

The option `.sourcePlateConv` will automatically convert the plate map to the requirements for generating dispensing files, which removes empty wells, or missing components from the plate map.

With all files imported and all the necessary list ready for use, we were finally able to generate the required dispensing files with the following function.

```
generateDispensingData(listofCombinations, listofDrugs, listofDoses,
listofVolumes, listofCtrls, listofStockConcentrations,
sourcePlate, listofExWells, .ctrlReplicates = 8,
.addUntreated = list(name = "Untreated", replicates = 8),
.finalWellVolume = 5, .plateFormat = 1536,
.destinationPlateID = "0920",
.randomizeDispensing = TRUE, .probeDispensing = FALSE)
```

The requirements for this function are relatively extensive, with a number of lists, as previously mentioned. We also provided the source plate as well as the list of excluded wells. In addition, we chose to use eight technical replicates for each control and added an additional untreated control to the experiment. Each control will be dispensed at the given number of replicates on each individual plate. We also had to specify the final volume per well and the plate format with the number of wells. Each experiment had a unique id, which was used to label each set of destination plates. We also opted-in to randomize all drug treatments across a set of plates.

The function will generate an R-object of class S3:dispensingData, with all the input data retained.

A helpful feature was the possibility of probing a potential dispensing based on the data provided, without actually generating any dispensing data. The probing provided a short summary and allowed us to estimate the number of drugs, controls, and plates among other things that were required for our drug screen.

The same can also be achieved once the dispensing data was generated through the use of the *summary* function. This can especially be beneficial, if an experimental design needs to be repeated after a period of time without remembering the exact details of the design, or if an experimental design was provided by another lab.

```
summary(dispensingData)
```

The function produced the following summary:

Summary for dispensing ID: 0920
Number of drug treatments: 30 378
Number of unique drug combinations: 29 280
Number of unique single drug treatments: 366
Number of excluded wells per plate: 156
Number of controls per plate: 24
Number of total plates: 23
Number of drugs: 61, Number of doses: 6/4

We can see from above that our experiment required a total number of 23 1536-well plates for a single experimental sample with 30 378 drug treatments. We had a total of 6 doses per drug of which 4 doses were used for the combination treatment.

In cases where only the dispensing data needs to be used, such as for any other data processing, the function *print* will extract only the dispensing data, without any metadata other any of the initial data sets.

```
print(dispensingData)
```

The function prints the dispensing file as shown below:

| [1] | [2] | [3] | [4] | [5] | [6] | [7] | [8] | [9] | [10] | [11] |
|---|---|---|---|---|---|---|---|---|---|---|
| 1 | NSC348884 | 81624-55-7 | 0,46875 | µM | 5 | C010 | D17 | E5 | 0920 | Plate22 |
| 2 | Avagacestat | 1146699-66-2 | 25 | µM | 5 | C010 | J15 | H25 | 0920 | Plate11 |
| 3 | GSK2879552 | 1401966-69-5 | 7,5 | µM | 5 | C010 | K3 | S36 | 0920 | Plate15 |
| 4 | CHIR-98014 | 252935-94-7 | 5 | µM | 5 | C010 | B15 | U41 | 0920 | Plate22 |
| 5 | CHIR-98014 | 252935-94-7 | 20 | µM | 5 | C010 | B13 | I16 | 0920 | Plate21 |

The headers are labeled as followed: [1] Combination.ID, [2] Sample Name, [3] CAS number, [4] Drug Concentration, [5] Unit, [6] Transfer Volume, [7] Source Plate Barcode, [8] Source Well, [9] Destination Well, [10] Destination Plate Barcode, [11] Plate Number

In addition, the dispensing data can be visualized on the basis of a plate map using the function *plot*.

```
plot(dispensingData, .saveto)
```

This will plot each plate of a given set and save it as an image/png file, if requested.

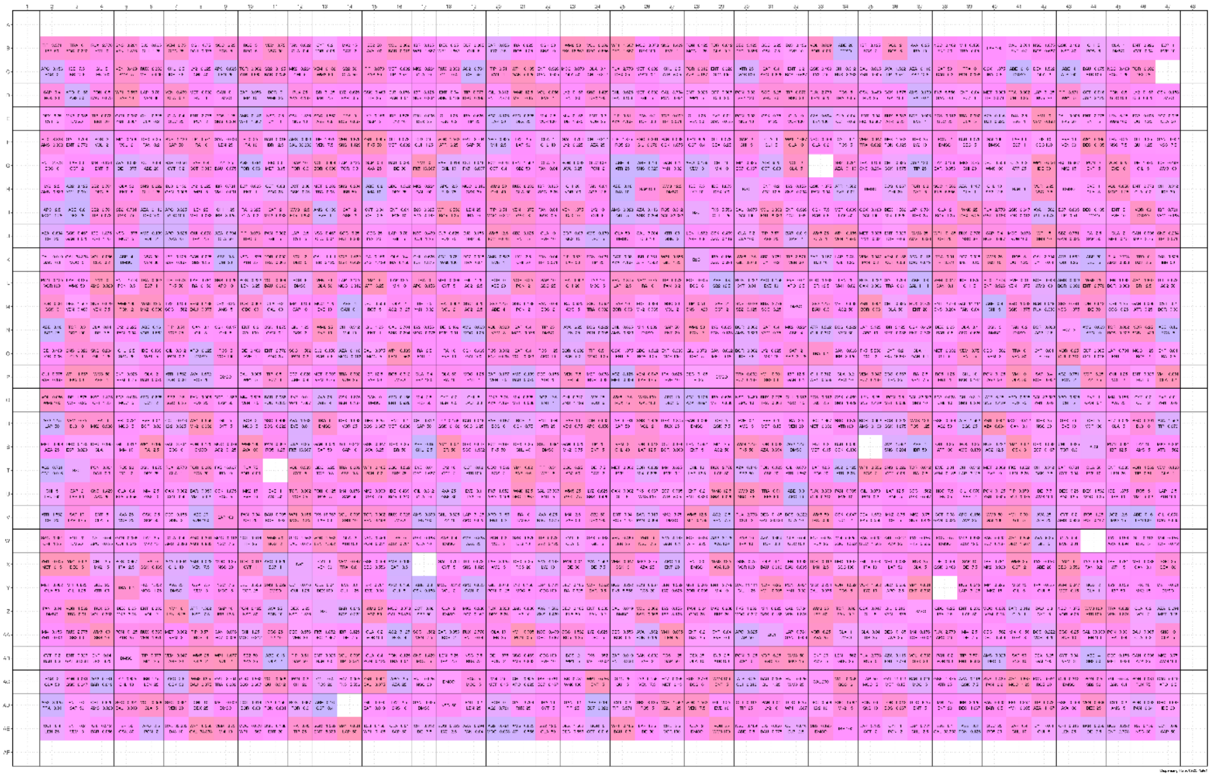

Supplementary Figure 3. Plate map showing one of 23 destination plates with randomized drug combination treatments, single drug treatments and eight positive, negative and untreated controls

Finally, the dispensing data was saved to a set of files, which were used to instruct the dispensing robot. This was achieved using the function *save*.

```
save(dispensingData, .saveto, .sets = 26, .labels = "alphabetic", .split = FALSE, .format = "Echo")
```

In the example above we generated 26 sets of plates, one for each experiment with a different cell line. We chose to label each set in alphabetic sequential order (A, B, C, … AA, AB, AC) and generate a single dispensing file. Alternatively, it would have been possible to split the dispensing data to individual files for each set. Since this study used an Echo Acoustic Liquid Handler (E5XX-1366, Echo 550) from Labcyte Inc., Beckman Coulter Life Sciences, the data format has been selected accordingly.

| Sample Name | Source Plate Barcode | Source Well | Transfer Volume | Destination Well |
|---|---|---|---|---|
| NSC348884 | C010 | D17 | 5 | E5 |
| Avagacestat | C010 | J15 | 5 | H25 |
| GSK2879552 | C010 | K3 | 5 | S36 |
| CHIR-98014 | C010 | B15 | 5 | U41 |
| CHIR-98014 | C010 | B13 | 5 | I16 |

**(b) post-experimental:**

Once the experiment has been run and the plates have been read, we were able to import the raw measurements with a dedicated function that reads raw data files from a specific plate reader.

To be able to correctly associate each plate with the exported raw data, we had to name each file with the specific plate barcode, or a unique plate identifier.

```
readFiles(.readfrom, .fileformat = c(".csv", ".txt"), .format = "EnVision")
```

Even though the raw measurement files were exported as csv files, we did specify both files in the function, as a premeditative measure. Since the experimental plates were read with the EnVision multimode plate reader, we specified the format accordingly, since most file formats are proprietary and specific to a certain manufacturer. This function was capable of detecting the export-format and identifying the used text delimiter, such as comma-, semicolon- or tab-separated. It also automatically detected the plate format and identified the raw data among all the metadata.

After all the raw data was imported, we could start consolidating the raw measurements with the dispensing data. In order to be able to map each plate and each set of plates to a given experiment, we had to provide a barcode reference, which included the unique plate id, the set, optionally the number of plates, but more importantly the experimental sample, which in our case was the MeWo human malignant melanoma cell line (RRID:CVCL_0445)

The barcode reference was imported from a csv file prior to consolidation of both data sets.

| PlateID | Set | Number | Sample |
|---|---|---|---|
| 0920 | A | 23 | MeWo |

```
barcodeReference <- read.csv2(file=file.path(“data/lib/platebarcode.csv"),
check.names=FALSE, header=TRUE, stringsAsFactors=FALSE,
colClasses=c("PlateID"="character"), comment.char = "#",
blank.lines.skip = TRUE, na.strings = "", sep = ";",
dec = ",", nrows = 1, skip=0)
```

```
consolidateData(dispensingData, rawMeasurements, barcodeReference)
```

Once both data set were consolidated, we were able to perform the first analysis by running the QC.

```
qc(consolidatedData, .ctrls = c("BzCl", "DMSO", "Untreated"), .qcMethod =
“all”)
```

We conducted the QC on both, the positive and negative control, as well as the untreated controls. We performed all available QC methods, which included the assessment of the *variance* between individual controls both, across all plates, as well as by individual plate. Another important QC method was *zprime*, assessing the distribution between the positive and negative control, in that case BzCl and DMSO, respectively. Additional methods included *emptywells*, providing an assessment of the signal in empty and untreated wells, which also included all the excluded wells, while *firstcolumn* did provide and assessment of the signal in wells of the first column for each plate.

The function returned an object of class S3:controlData, with the data and plots for each method. The data structure is shown below.

| | | |
|---|---|---|
| qcdata | list [1] (S3: controlData) | List of length 1 |
| MeWo | list [5] | List of length 5 |
| data | list [329 x 14] (S3: data.fram | A data.frame with 329 rows and 14 columns |
| variance | list [3] | List of length 3 |
| data | list [3] | List of length 3 |
| boxplot | list [3] | List of length 3 |
| boxplot-byplate | list [3] | List of length 3 |
| empty-wells | list [23] | List of length 23 |
| first-column | list [9] (S3: gg, ggplot) | List of length 9 |
| z-factor | list [2] | List of length 2 |
| data | list [23 x 4] (S3: data.frame) | A data.frame with 23 rows and 4 columns |
| plot | list [9] (S3: gg, ggplot) | List of length 9 |

Supplementary Figure 4. Data structure of the quality control assessment output

The function also generated a number of plots, providing an insight into the overall ***variance*** and noise based on the used controls. One of the plots compares the raw signal between the controls across all plates and provides a first assessment of variance within an experiment. In Supplementary Figure 5, we can see that the untreated control and the negative control (DMSO) have the same signal strength and level of variance as expected, compared to the positive control (BzCl) with a signal strength close to zero.

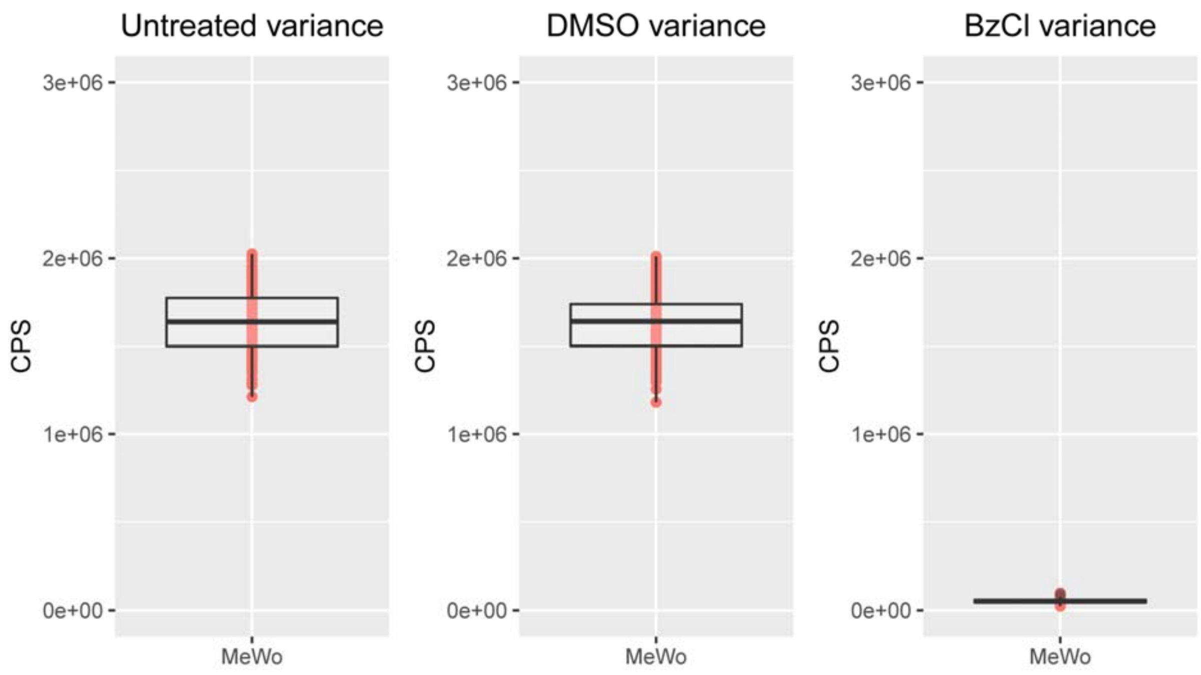

Supplementary Figure 5. Raw signal (CPS) distribution for each control

Another plot shows the variance of controls across all individual plates. Here as well, the pattern of signal strength and variance is quite similar between the untreated and negative control. This plot allows to identify potential irregularities on individual plates. We could observe individual outliers in the negative controls (DMSO), such as on plate 3, shown in Supplementary Figure 6. This issue was circumvented through the use of multiple replicates for each control and choosing the median signal for any downstream analysis.

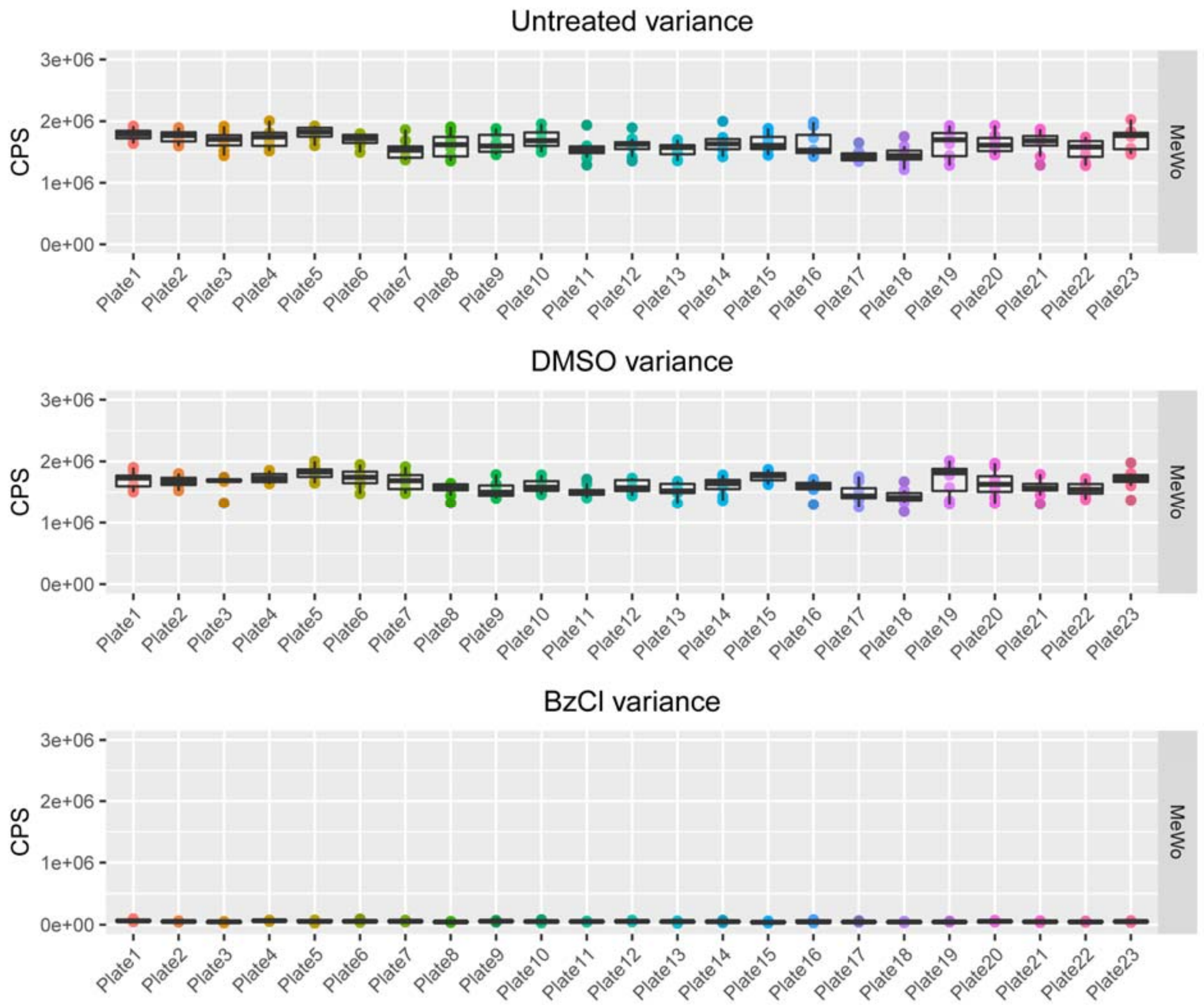


Supplementary Figure 6. Raw signal (CPS) distribution for each of the controls across all plates

With another plot, it was possible to identify those outliers on a well to well basis. All wells outside a certain tolerance level are labeled. This allows to identify potential technical issues in dispensing, if certain wells are repetitively affected.

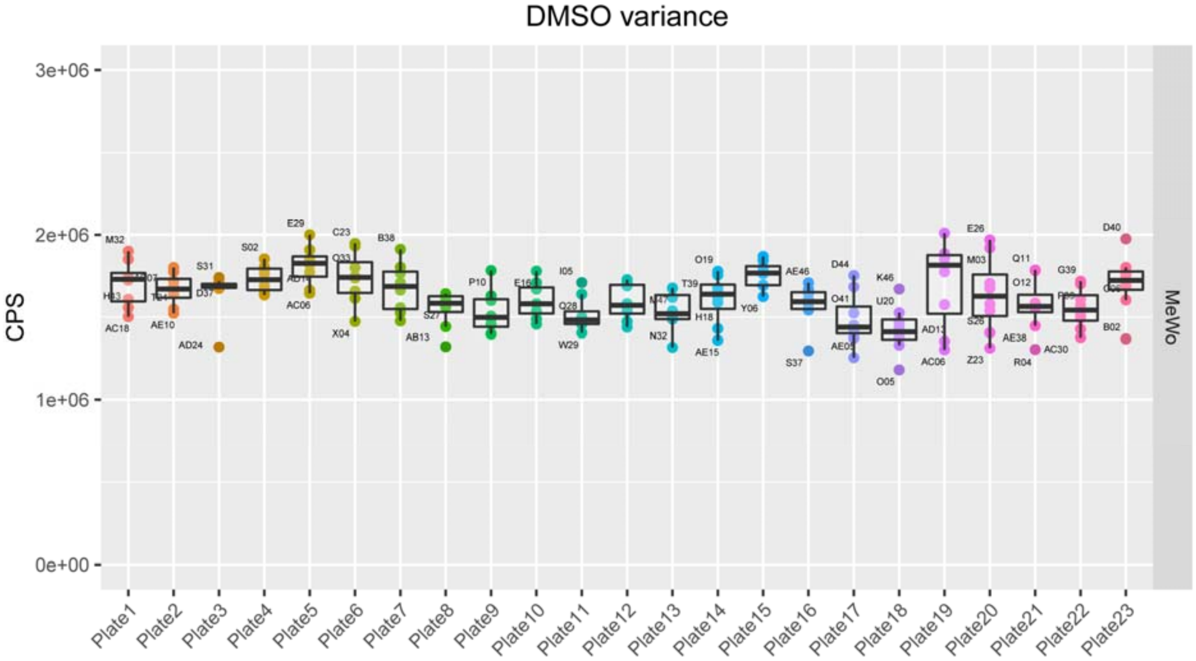


Supplementary Figure 7. Variance distribution indicating wells of individual outliers

Another set of plots looks at the signal of all the empty and excluded wells on each individual plate. It might give insight into certain pasterns of variance across an individual plate and reveal certain dispensing issues.

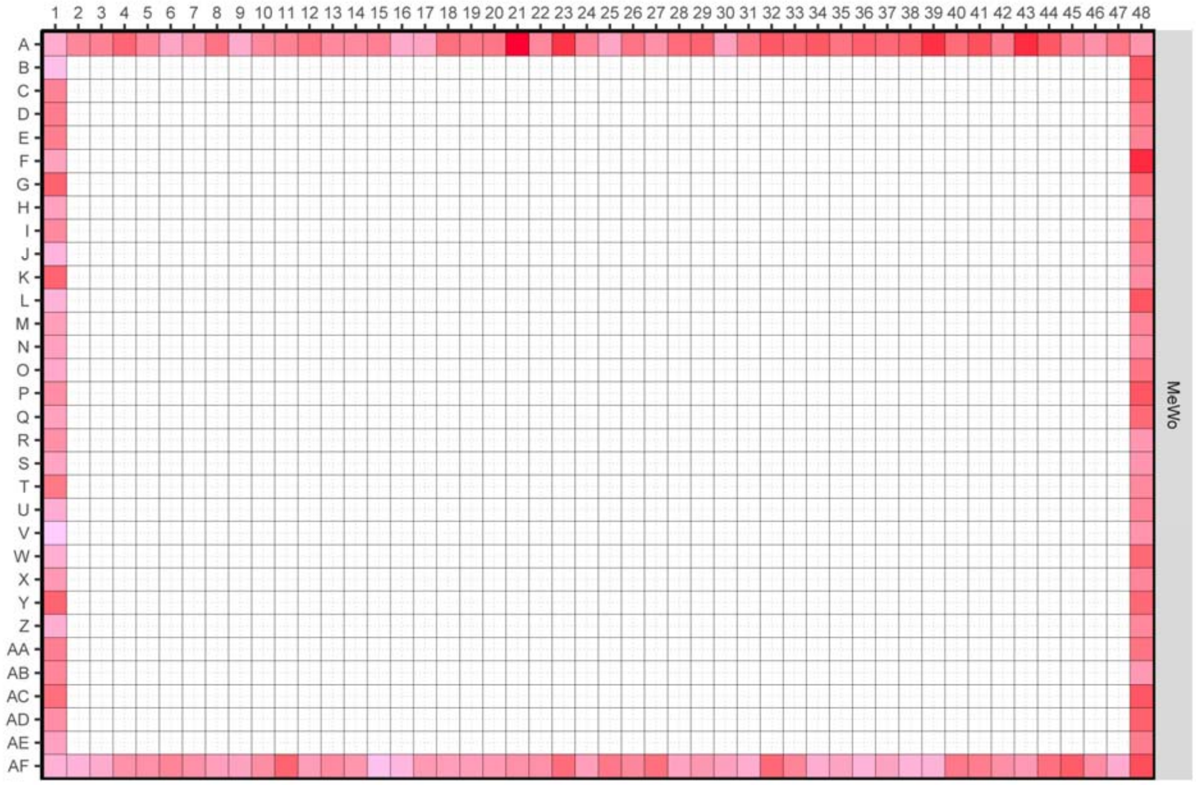


Supplementary Figure 8. Signal measurement of the outer (excluded) wells

A variation of this plot extracts the first column of each plate and plots them side by side in order to identify potential patterns of variation across multiple plates.

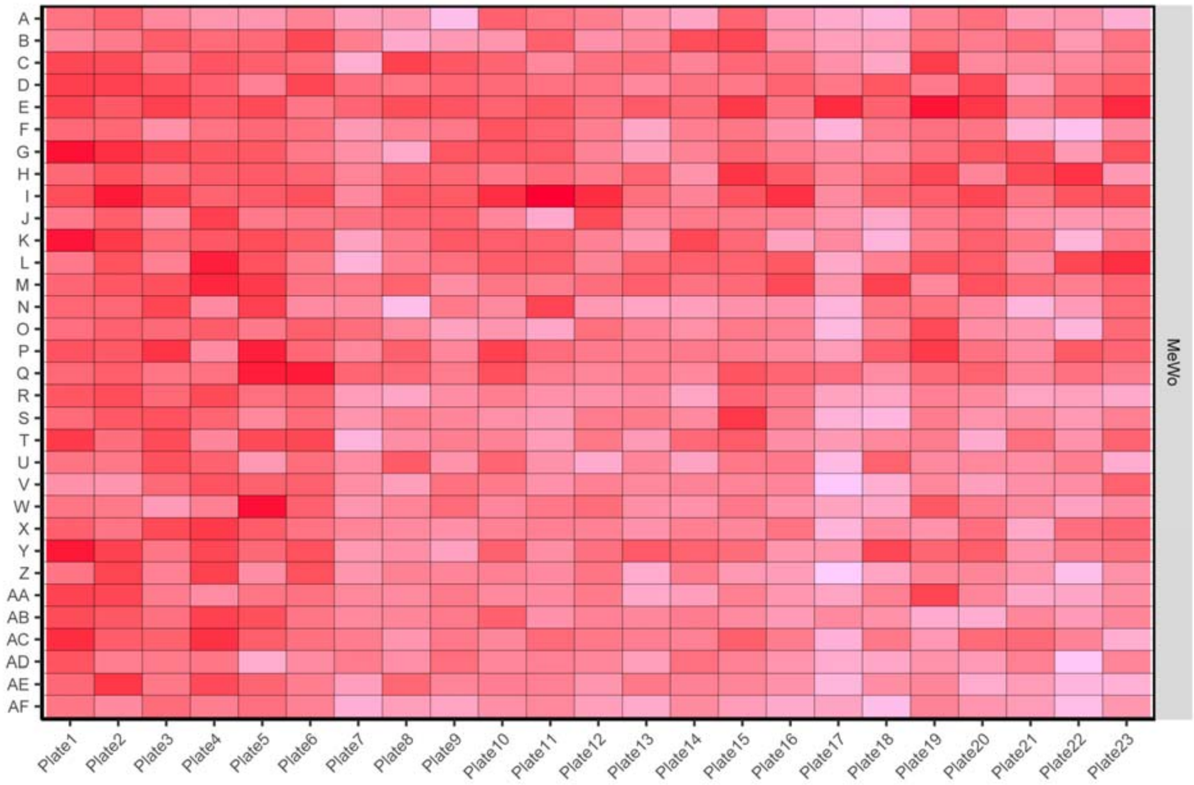


Supplementary Figure 9. Signal distribution of (excluded) wells in the first column of each plate

One of the more important qc assessments is provided by *Z'-factor*, which is assessing the distribution between the positive and negative control and an indicator how reliable and trustworthy the screen can be considered. The associated plot shows the *Z'-factor* for a set of controls on each individual plate, as shown in Supplementary Figure 10. Any score above the threshold of 0.5 can be considered as reliable, with a wide enough separation between the positive and negative control to be able to distinguish any measurements that fall between those points.

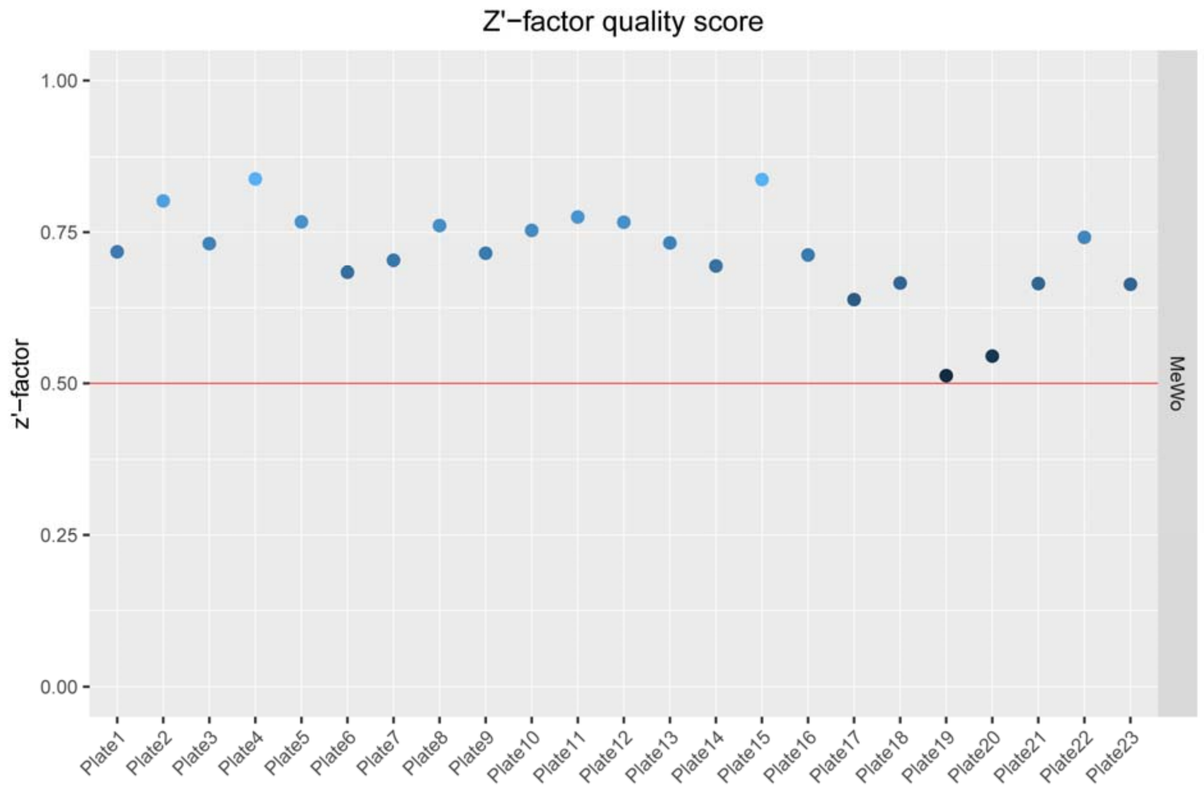


Supplementary Figure 10. Z'-factor for each set of controls across all plates

Once the data passed the QC, we continued with processing the data for downstream analysis using the function

```
processData(consolidatedData, .ctrls=list(positive="BzCl",
negative="DMSO"))
```

The first step that this module performs is the normalization of the raw measurements to the positive and negative control, followed by splitting the data into individual data sets, one for the controls, single drug treatments and combination treatments. Finally, the data is re-formatted into a table and a matrix for each drug pair, representing the dose response between two drugs at each dose.

The function returned an object of class S3:processedData, which was then used to run the single drug assessment. There are two main module linages post data assembly. The first one is concerned with single drug treatments and their responses, while second with the drug combination treatments and their drug interaction effects.

Once the data has been normalized and individual data sets assembled, we were able to perform the first analysis, assessing the single drug responses by running a dose response model for each drug.

```
runDRM(processedData, .saveto, .plot = TRUE)
```

The function will analyze single drug treatments. It performs curve fitting using a four-parameter log-logistic function (LL.4), and estimates the EC10, EC50 and EC90. The function also plots the single drug curves by viability and inhibition for each single drug.

An excerpt of the output data is shown below along with a few plots.

| Sample | Drug | ED | Estimate | Std. Error |
|---|---|---|---|---|
| MeWo | Abemaciclib mesylate (LY2835219) | 10 | 18,5166585030146 | 139,447775779473 |
| MeWo | Abemaciclib mesylate (LY2835219) | 50 | 18,7410035107027 | 120,298549656558 |
| MeWo | Abemaciclib mesylate (LY2835219) | 90 | 18,9680666482556 | 101,30964028357 |
| MeWo | AG-221 (Enasidenib) | 10 | 11,5666411485189 | 4,96699580857896 |
| MeWo | AG-221 (Enasidenib) | 50 | 36,2896471575161 | 12,3114505023091 |

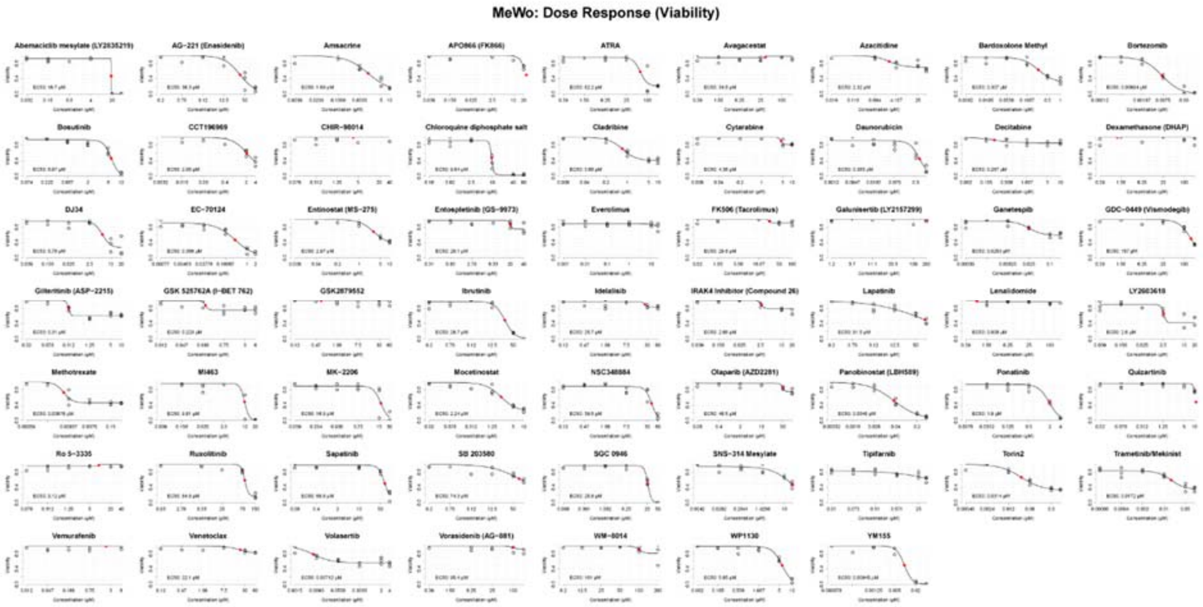


Supplementary Figure 11. Single drug response curves by viability for all drugs

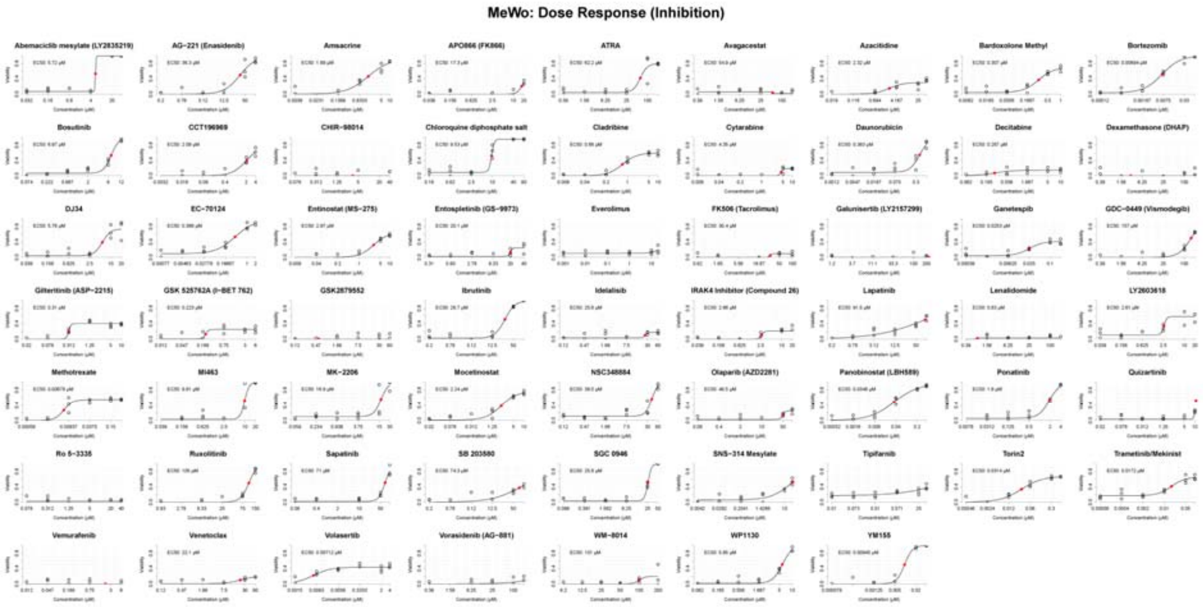


Supplementary Figure 12. Single drug response curves by inhibition for all drugs

In addition to the basic plots with only the dose response curves illustrated, another set of plots can be generated containing the dose response matrix along with the individual dose response curves for each single drug of a given pair in a composite plot.

```
customPlotting(processedData, doseRespModel, .saveto)
```

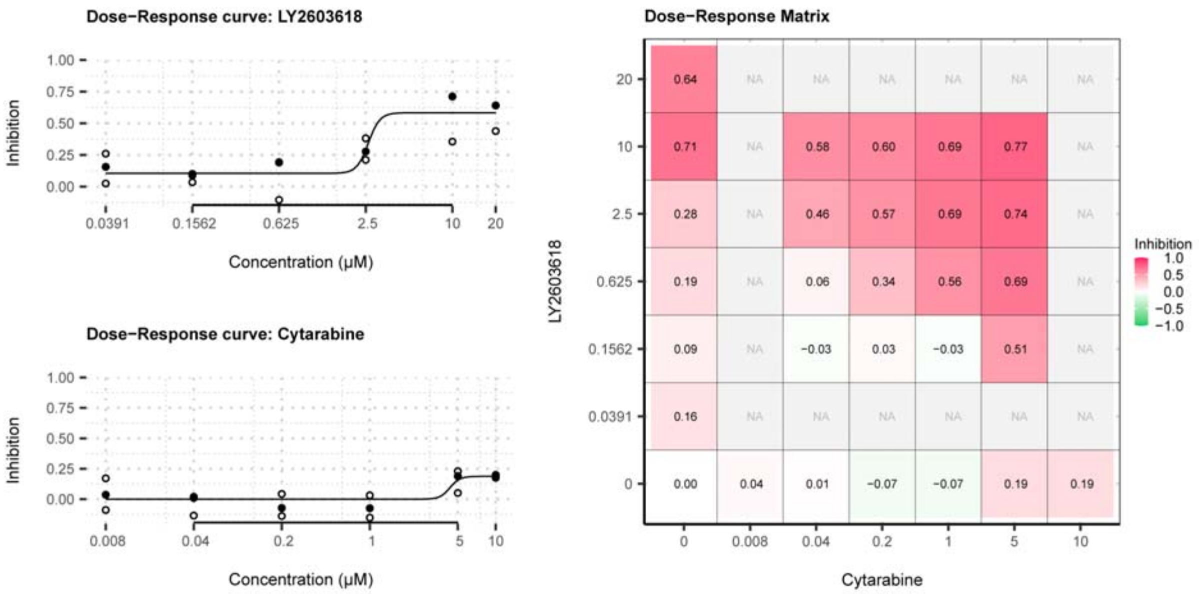


Supplementary Figure 13. Dose response curves and matrix for a drug (combination) pair. Single dose-response curves (left) with measurement points in triplicates for each concentration and the median indicated by a black point. Median drug responses also shown in the dose-response matrix (right) for single drug treatments.

Furthermore, the function will also plot an overview of single drug responses for each sample, as well as for each drug.

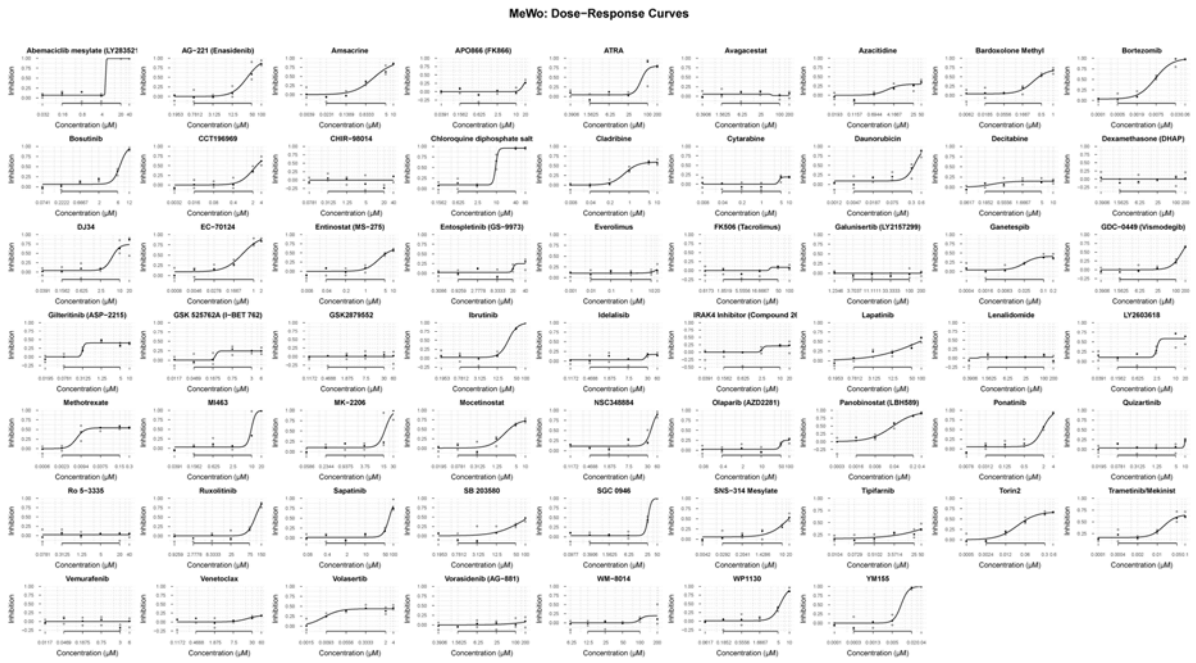


Supplementary Figure 14. Single drug response curves of all drugs for a given cell line

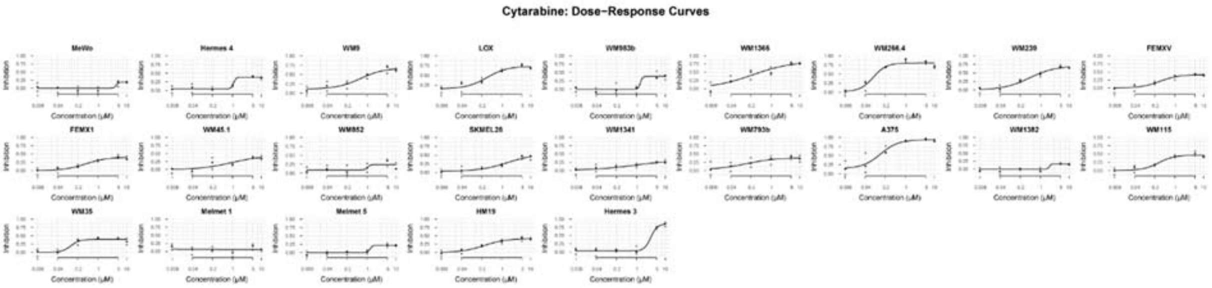


Supplementary Figure 15. Single drug response curves of all cell lines for a given drug

With the single drug response data, it is possible to assess the dynamic drug-activity range for individual drugs. The function estimates the dynamic drug-activity range (DDAR) across a number of doses for each drug response and generates a set of plots. The DDAR is predefined as the range between the ED10 and ED90.

```
dynamicRange(doseRespModel)
```

One set of plots looks at the dynamic range for each individual drug and highlights potential drugs that do not show any effect across all doses.

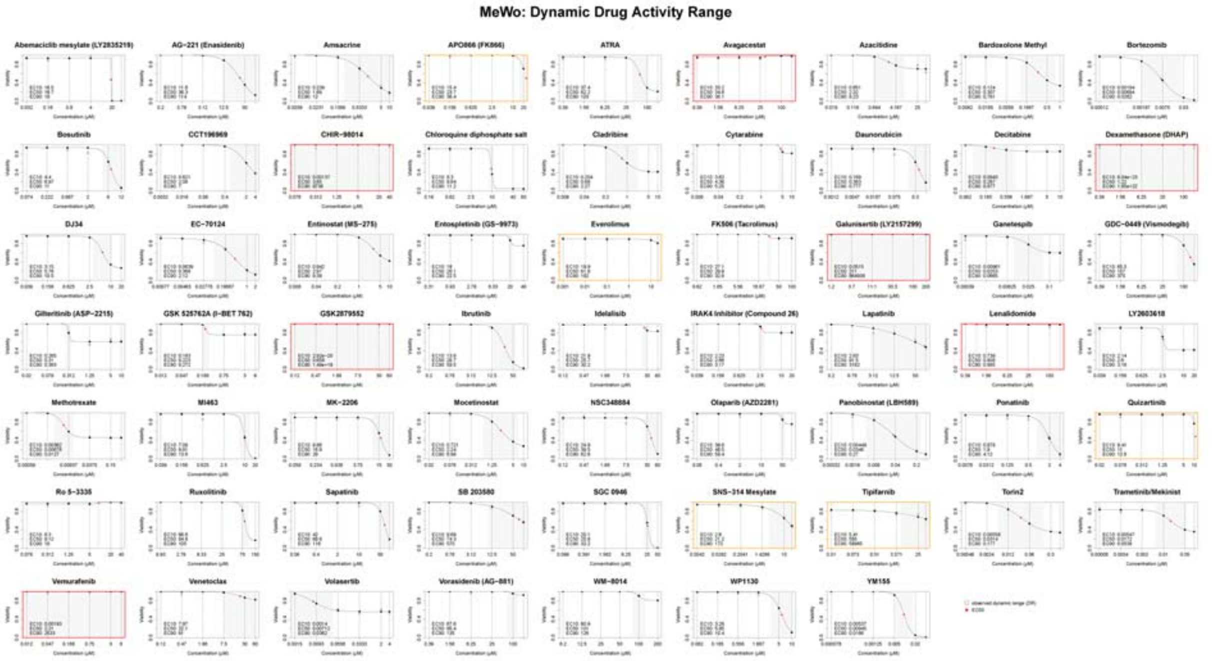


Supplementary Figure 16. Dynamic drug-activity range for all drugs

Another set of plots looks at the average DDAR across all samples for a single drug.

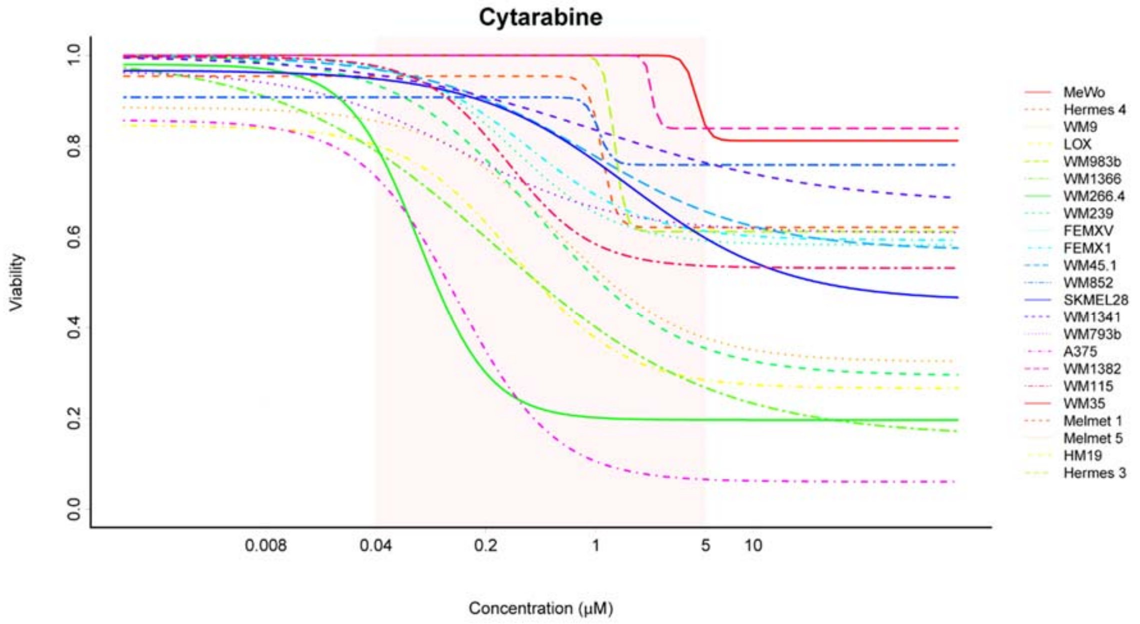


Supplementary Figure 17. Dynamic drug-activity range for all cell lines

This function will also plot the DDAR across all samples for all the individual drugs indicating the expected drug activity range compared to the observed DDAR.

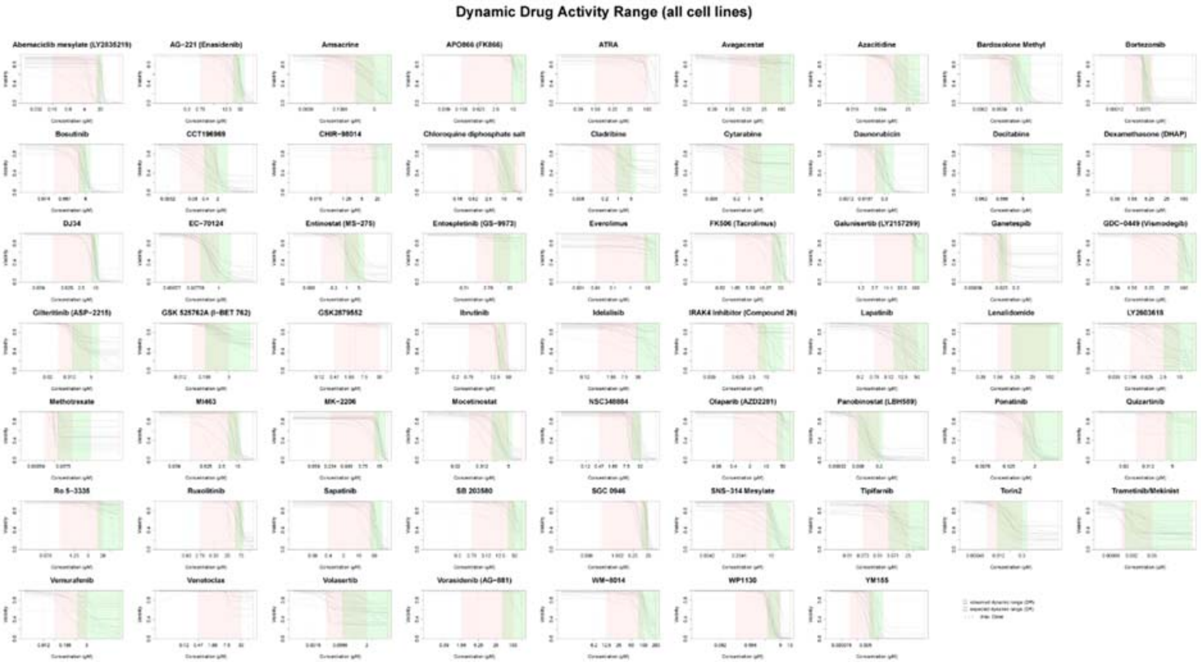


Supplementary Figure 18. Dynamic drug-activity range for all drugs and cell lines

The same graph is also shown without the indicated DDAR and without the curves fitted to the points, but rather showing the raw fit of the dose responses.

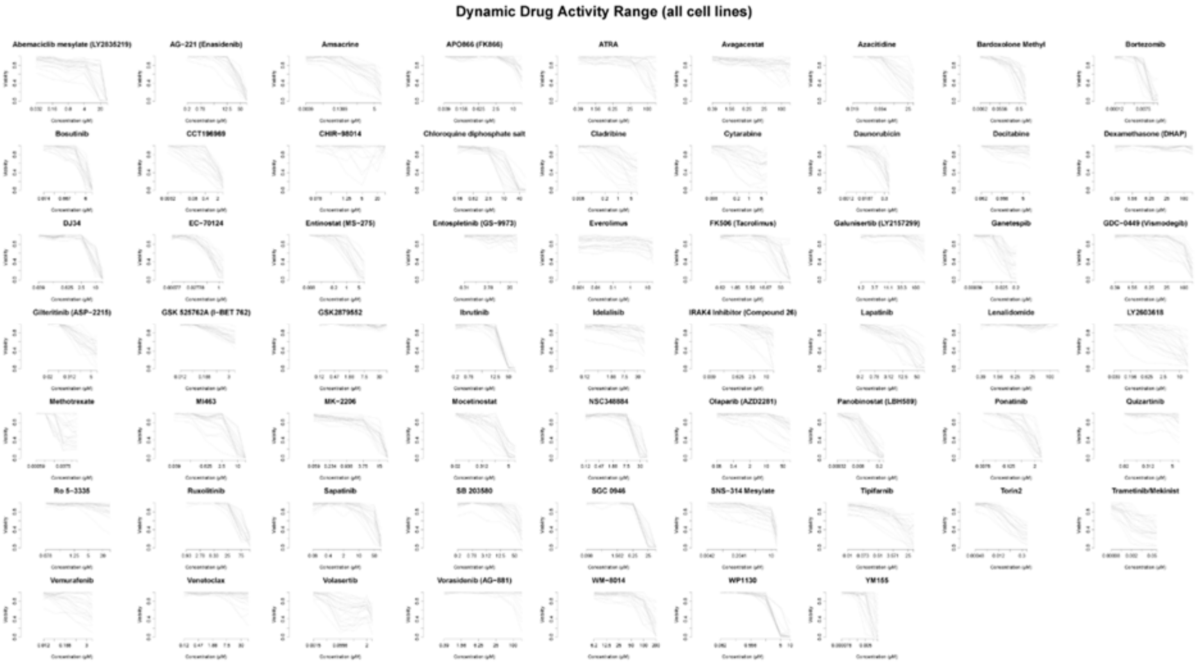

Supplementary Figure 19. Unfitted dose response curves for all drugs

In the second module linage, the drug combinations are assessed by estimating synergy and antagonism for each individual drug pair. This is being done primarily with the R-package bayesynergy (Rønneberg et al., 2021), which was specifically designed for this pipeline and with large drug-sensitivity screens in mind.

```
bayesynergy(processedData, .saveoutput = TRUE, .plot = TRUE, .saveto)
```

The function uses data of class S3:processedData and runs bayesynergy with a predefined set of parameters for ease of use. The actual output and plots of bayesynergy can be saved to file.

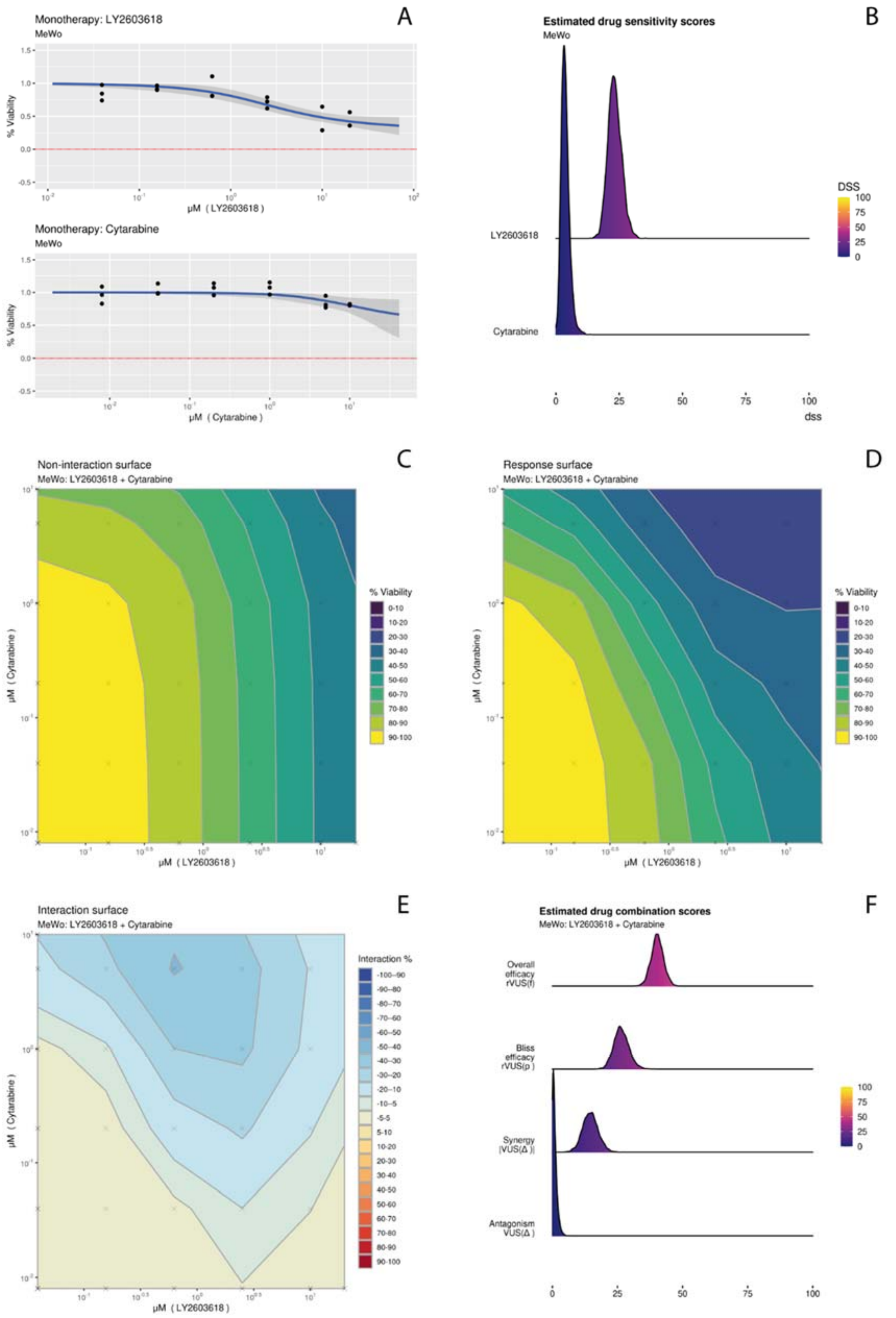


Supplementary Figure 20. Graphs from bayesynergy showing A. monotherapies, B. drug sensitivity scores (DSS), C. non-interaction surface, D. response surface, E. interaction surface and F. drug combination scores based on the volume under the surface (VUS) for both the synergy and antagonistic effect as described by Rønneberg, 2021, Brief Bioinform.

The plots include the drug response as monotherapies for the corresponding drug pair and a histogram with the drug sensitivity scores (DSS). Furthermore, the non-interaction, interaction and response surface are plotted. Another plot shows the estimated drug combination scores.

The function returned an object of class S3:bayesdata, which then was used to score the drug interactions.

```
synergyScoring(bayDFS, .saveoutput = TRUE, .plot = TRUE, .saveto)
```

The estimated synergies were then subject to scoring, in which the drug pairs are ranked based on the highest synergy. In addition to the synergy scores, additional statistical data is being provided along with a number of QC parameters. The data was saved as a csv file and a number of plots were generated. The first plot shows both the synergy and antagonistic scores for all drug pairs, while the second plot shows only the synergy scores for each drug and its corresponding drug pair. Another plot shows a more detailed distribution of synergy scores for each individual drug.

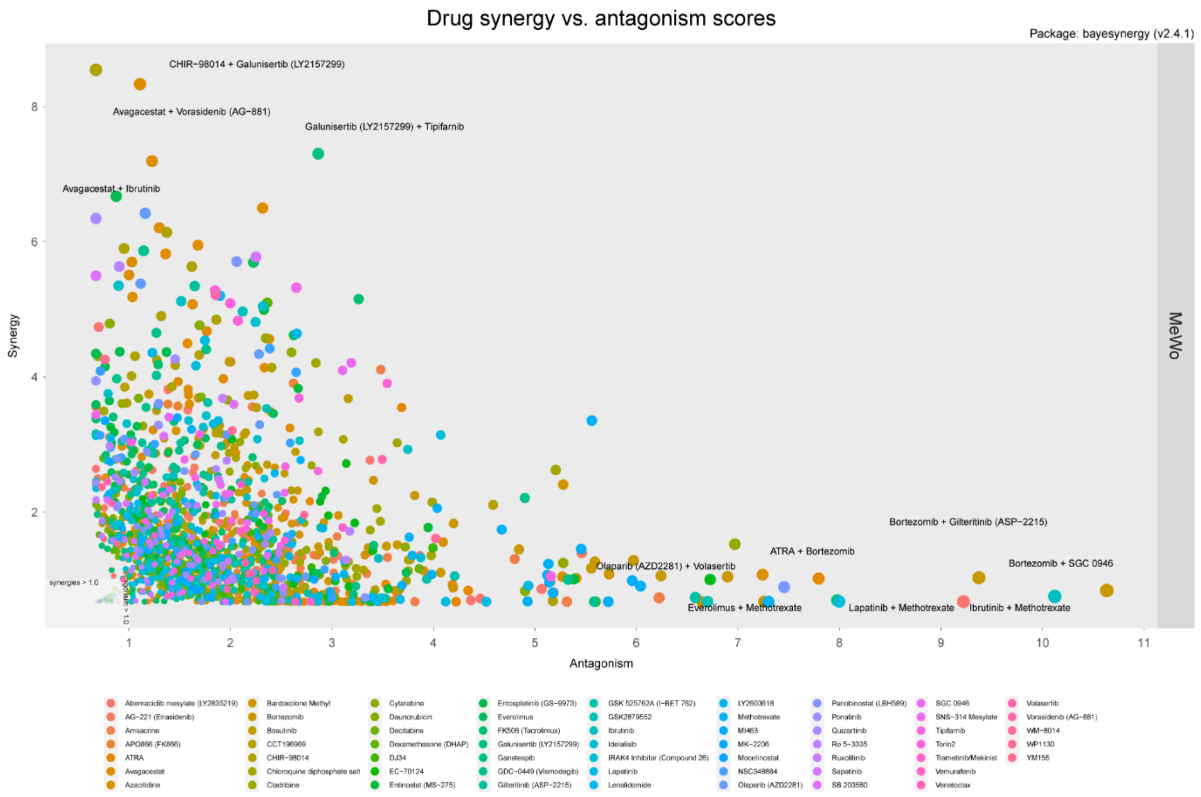


Supplementary Figure 21. Drug synergy vs. antagonism scores

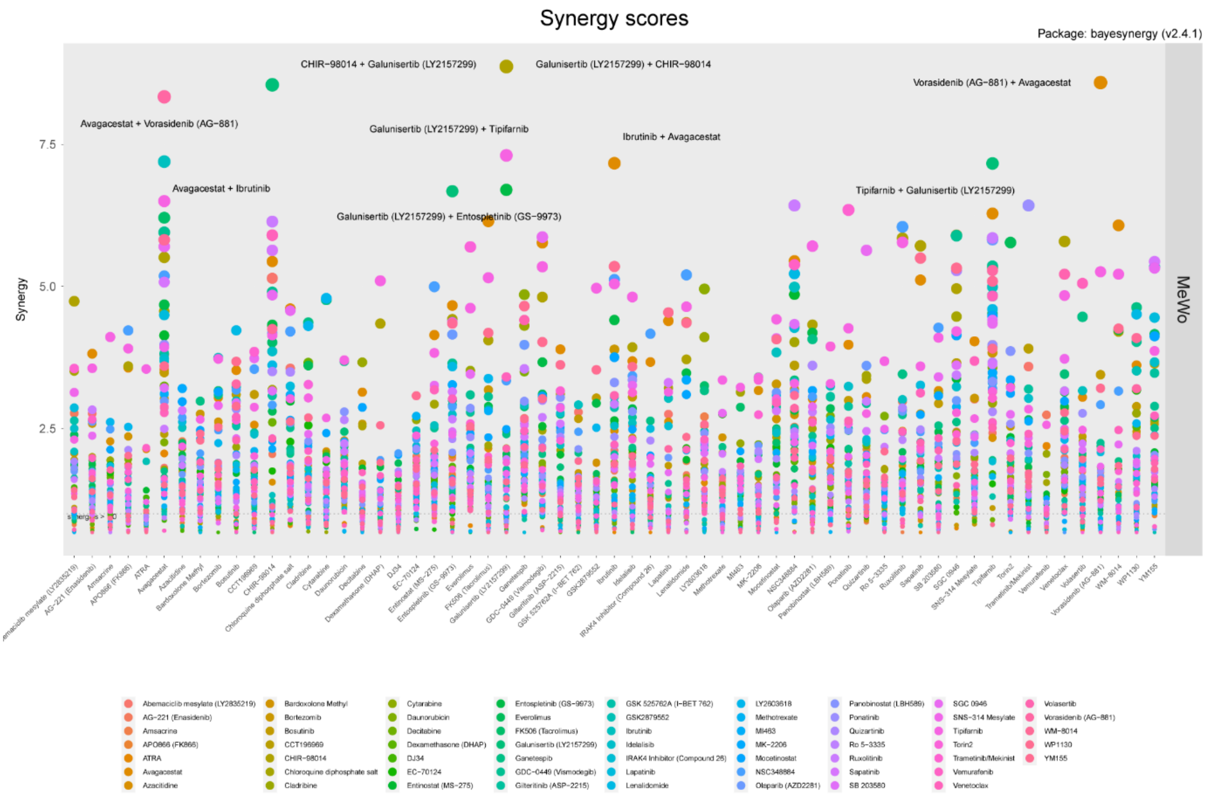


Supplementary Figure 22. Drug synergy scores for each drug

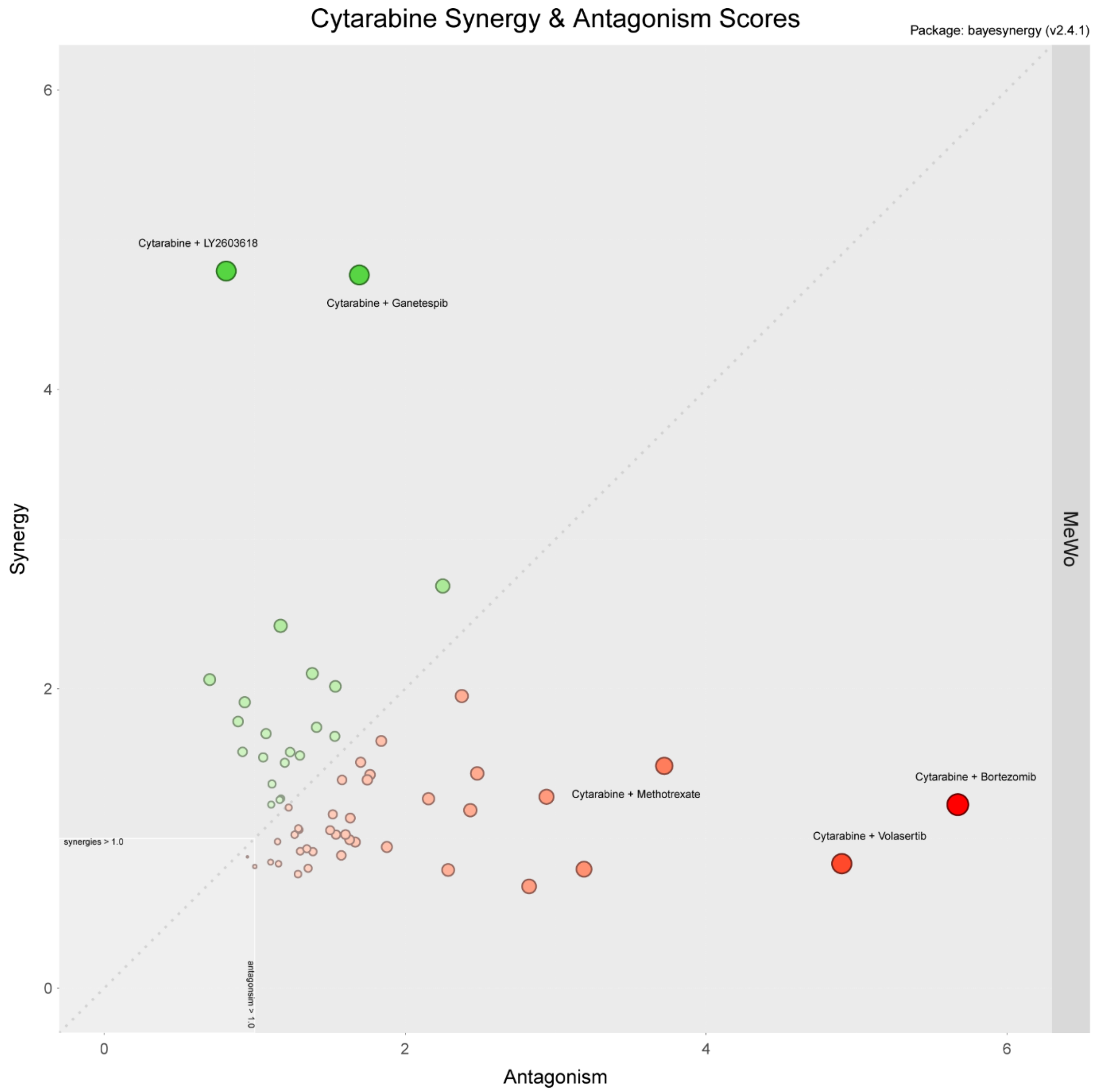


Supplementary Figure 23. Drug synergy and antagonism scores for each individual drug
