## Supplementary Figures (1-4) for "metascreen: A modular tool for the design and analysis of drug combination screens"

| | 1 | 2 | 3 | 4 | 5 | 6 | 7 | 8 | 9 | 10 | 11 | 12 | 13 | 14 | 15 | 16 | 17 | 18 | 19 | 20 | 21 | 22 | 23 | 24 |
|---|---|---|---|---|---|---|---|---|---|---|---|---|---|---|---|---|---|---|---|---|---|---|---|---|
| A | Venetoclax 30 mM | | Venetoclax 7.5 mM | Venetoclax 1.88 mM | Venetoclax 0.47 mM | Venetoclax 0.12 mM | Cladribine 5 mM | | Cladribine 1 mM | Cladribine 0.2 mM | Cladribine 0.04 mM | Cladribine 0.01 mM | AG-221 (Enasidenib) 50 mM | | AG-221 (Enasidenib) 12.5 mM | AG-221 (Enasidenib) 3.13 mM | AG-221 (Enasidenib) 0.78 mM | AG-221 (Enasidenib) 0.2 mM | Chloroquine diphosphate salt 40 mM | | Chloroquine diphosphate salt 10 mM | Chloroquine diphosphate salt 2.5 mM | Chloroquine diphosphate salt 0.63 mM | Chloroquine diphosphate salt 0.16 mM |
| B | Methotrexate 0.15 mM | | Methotrexate 0.04 mM | Methotrexate 0.01 mM | Methotrexate 2.34e-3 mM | Methotrexate 5.86e-4 mM | Cytarabine 5 mM | | Cytarabine 1 mM | Cytarabine 0.2 mM | Cytarabine 0.04 mM | Cytarabine 0.01 mM | CHIR-98014 20 mM | | CHIR-98014 5 mM | CHIR-98014 1.25 mM | CHIR-98014 0.31 mM | CHIR-98014 0.08 mM | Ganetespib 0.1 mM | | Ganetespib 0.03 mM | Ganetespib 0.01 mM | Ganetespib 1.56e-3 mM | Ganetespib 3.91e-4 mM |
| C | Daunorubicin 0.3 mM | | Daunorubicin 0.08 mM | Daunorubicin 0.02 mM | Daunorubicin 4.69e-3 mM | Daunorubicin 1.17e-3 mM | CCT196969 2 mM | | CCT196969 0.4 mM | CCT196969 0.08 mM | CCT196969 0.02 mM | CCT196969 3.2e-3 mM | APO866 (FK866) 10 mM | | APO866 (FK866) 2.5 mM | APO866 (FK866) 0.63 mM | APO866 (FK866) 0.16 mM | APO866 (FK866) 0.04 mM | Vemurafenib 3 mM | | Vemurafenib 0.75 mM | Vemurafenib 0.19 mM | Vemurafenib 0.05 mM | Vemurafenib 0.01 mM |
| D | Lenalidomide 100 mM | | Lenalidomide 25 mM | Lenalidomide 6.25 mM | Lenalidomide 1.56 mM | Lenalidomide 0.39 mM | Entinostat (MS-275) 5 mM | | Entinostat (MS-275) 1 mM | Entinostat (MS-275) 0.2 mM | Entinostat (MS-275) 0.04 mM | Entinostat (MS-275) 0.01 mM | NSC348884 30 mM | | NSC348884 7.5 mM | NSC348884 1.88 mM | NSC348884 0.47 mM | NSC348884 0.12 mM | DJ34 10 mM | | DJ34 2.5 mM | DJ34 0.63 mM | DJ34 0.16 mM | DJ34 0.04 mM |
| E | GSK 525762A (I-BET 762) 3 mM | | GSK 525762A (I-BET 762) 0.75 mM | GSK 525762A (I-BET 762) 0.19 mM | GSK 525762A (I-BET 762) 0.05 mM | GSK 525762A (I-BET 762) 0.01 mM | Abemaciclib mesylate (LY2835219) 20 mM | | Abemaciclib mesylate (LY2835219) 4 mM | Abemaciclib mesylate (LY2835219) 0.8 mM | Abemaciclib mesylate (LY2835219) 0.16 mM | Abemaciclib mesylate (LY2835219) 0.03 mM | Gilteritinib (ASP-2215) 5 mM | | Gilteritinib (ASP-2215) 1.25 mM | Gilteritinib (ASP-2215) 0.31 mM | Gilteritinib (ASP-2215) 0.08 mM | Gilteritinib (ASP-2215) 0.02 mM | WM-8014 100 mM | | WM-8014 50 mM | WM-8014 25 mM | WM-8014 12.5 mM | WM-8014 6.25 mM |
| F | Mocetinostat 5 mM | | Mocetinostat 1.25 mM | Mocetinostat 0.31 mM | Mocetinostat 0.08 mM | Mocetinostat 0.02 mM | Olaparib (AZD2281) 50 mM | | Olaparib (AZD2281) 10 mM | Olaparib (AZD2281) 2 mM | Olaparib (AZD2281) 0.4 mM | Olaparib (AZD2281) 0.08 mM | MK-2206 15 mM | | MK-2206 3.75 mM | MK-2206 0.94 mM | MK-2206 0.23 mM | MK-2206 0.06 mM | Everolimus 10 mM | | Everolimus 1 mM | Everolimus 0.1 mM | Everolimus 0.01 mM | Everolimus 1e-3 mM |
| G | Quizartinib 5 mM | | Quizartinib 1.25 mM | Quizartinib 0.31 mM | Quizartinib 0.08 mM | Quizartinib 0.02 mM | Torin2 0.3 mM | | Torin2 0.06 mM | Torin2 0.01 mM | Torin2 2.4e-3 mM | Torin2 4.8e-4 mM | Ro 5-3335 20 mM | | Ro 5-3335 5 mM | Ro 5-3335 1.25 mM | Ro 5-3335 0.31 mM | Ro 5-3335 0.08 mM | Bosutinib 6 mM | | Bosutinib 2 mM | Bosutinib 0.67 mM | Bosutinib 0.22 mM | Bosutinib 0.07 mM |
| H | ATRA 100 mM | | ATRA 25 mM | ATRA 6.25 mM | ATRA 1.56 mM | ATRA 0.39 mM | Trametinib/Mekinist 0.05 mM | | Trametinib/Mekinist 0.01 mM | Trametinib/Mekinist 2e-3 mM | Trametinib/Mekinist 4e-4 mM | Trametinib/Mekinist 8e-5 mM | IRAK4 Inhibitor (Compound 26) 10 mM | | IRAK4 Inhibitor (Compound 26) 2.5 mM | IRAK4 Inhibitor (Compound 26) 0.63 mM | IRAK4 Inhibitor (Compound 26) 0.16 mM | IRAK4 Inhibitor (Compound 26) 0.04 mM | Decitabine 5 mM | | Decitabine 1.67 mM | Decitabine 0.56 mM | Decitabine 0.19 mM | Decitabine 0.06 mM |
| I | | | | | | | Panobinostat (LBH589) 0.2 mM | | Panobinostat (LBH589) 0.04 mM | Panobinostat (LBH589) 0.01 mM | Panobinostat (LBH589) 1.6e-3 mM | Panobinostat (LBH589) 3.2e-4 mM | Dexamethasone (DHAP) 100 mM | | Dexamethasone (DHAP) 25 mM | Dexamethasone (DHAP) 6.25 mM | Dexamethasone (DHAP) 1.56 mM | Dexamethasone (DHAP) 0.39 mM | Bardoxolone Methyl 0.5 mM | | Bardoxolone Methyl 0.17 mM | Bardoxolone Methyl 0.06 mM | Bardoxolone Methyl 0.02 mM | Bardoxolone Methyl 0.01 mM |
| J | Ponatinib 2 mM | | Ponatinib 0.5 mM | Ponatinib 0.13 mM | Ponatinib 0.03 mM | Ponatinib 0.01 mM | Sapatinib 50 mM | | Sapatinib 10 mM | Sapatinib 2 mM | Sapatinib 0.4 mM | Sapatinib 0.08 mM | Avagacestat 100 mM | | Avagacestat 25 mM | Avagacestat 6.25 mM | Avagacestat 1.56 mM | Avagacestat 0.39 mM | | | | | | |
| K | GSK2879552 30 mM | | GSK2879552 7.5 mM | GSK2879552 1.88 mM | GSK2879552 0.47 mM | GSK2879552 0.12 mM | Amsacrine 5 mM | | Amsacrine 0.83 mM | Amsacrine 0.14 mM | Amsacrine 0.02 mM | Amsacrine 3.86e-3 mM | Idelalisib 30 mM | | Idelalisib 7.5 mM | Idelalisib 1.88 mM | Idelalisib 0.47 mM | Idelalisib 0.12 mM | Entospletinib (GS-9973) 20 mM | | Entospletinib (GS-9973) 8.33 mM | Entospletinib (GS-9973) 2.78 mM | Entospletinib (GS-9973) 0.93 mM | Entospletinib (GS-9973) 0.31 mM |
| L | Vorasidenib (AG-881) 100 mM | | Vorasidenib (AG-881) 25 mM | Vorasidenib (AG-881) 6.25 mM | Vorasidenib (AG-881) 1.56 mM | Vorasidenib (AG-881) 0.39 mM | Azacitidine 25 mM | | Azacitidine 4.17 mM | Azacitidine 0.69 mM | Azacitidine 0.12 mM | Azacitidine 0.02 mM | SGC 0946 25 mM | | SGC 0946 6.25 mM | SGC 0946 1.56 mM | SGC 0946 0.39 mM | SGC 0946 0.1 mM | | | | | | |
| M | Ibrutinib 50 mM | | Ibrutinib 12.5 mM | Ibrutinib 3.13 mM | Ibrutinib 0.78 mM | Ibrutinib 0.2 mM | Volasertib 2 mM | | Volasertib 0.33 mM | Volasertib 0.06 mM | Volasertib 0.01 mM | Volasertib 1.54e-3 mM | MI463 10 mM | | MI463 2.5 mM | MI463 0.63 mM | MI463 0.16 mM | MI463 0.04 mM | Galunisertib (LY2157299) 100 mM | | Galunisertib (LY2157299) 33.33 mM | Galunisertib (LY2157299) 11.11 mM | Galunisertib (LY2157299) 3.7 mM | Galunisertib (LY2157299) 1.23 mM |
| N | LY2603618 10 mM | | LY2603618 2.5 mM | LY2603618 0.63 mM | LY2603618 0.16 mM | LY2603618 0.04 mM | EC-70124 1 mM | | EC-70124 0.17 mM | EC-70124 0.03 mM | EC-70124 4.63e-3 mM | EC-70124 7.72e-4 mM | Bortezomib 0.03 mM | | Bortezomib 0.01 mM | Bortezomib 1.88e-3 mM | Bortezomib 4.69e-4 mM | Bortezomib 1.17e-4 mM | FK506 (Tacrolimus) 50 mM | | FK506 (Tacrolimus) 16.67 mM | FK506 (Tacrolimus) 5.56 mM | FK506 (Tacrolimus) 1.85 mM | FK506 (Tacrolimus) 0.62 mM |
| O | YM155 0.02 mM | | YM155 0.01 mM | YM155 1.25e-3 mM | YM155 3.13e-4 mM | YM155 7.81e-5 mM | SNS-314 Mesylate 10 mM | | SNS-314 Mesylate 1.43 mM | SNS-314 Mesylate 0.2 mM | SNS-314 Mesylate 0.03 mM | SNS-314 Mesylate 4.16e-3 mM | SB 203580 50 mM | | SB 203580 12.5 mM | SB 203580 3.13 mM | SB 203580 0.78 mM | SB 203580 0.2 mM | WP1130 5 mM | | WP1130 1.67 mM | WP1130 0.56 mM | WP1130 0.19 mM | WP1130 0.06 mM |
| P | GDC-0449 (Vismodegib) 100 mM | | GDC-0449 (Vismodegib) 25 mM | GDC-0449 (Vismodegib) 6.25 mM | GDC-0449 (Vismodegib) 1.56 mM | GDC-0449 (Vismodegib) 0.39 mM | Tipifarnib 25 mM | | Tipifarnib 3.57 mM | Tipifarnib 0.51 mM | Tipifarnib 0.07 mM | Tipifarnib 0.01 mM | Lapatinib 50 mM | | Lapatinib 12.5 mM | Lapatinib 3.13 mM | Lapatinib 0.78 mM | Lapatinib 0.2 mM | Ruxolitinib 75 mM | | Ruxolitinib 25 mM | Ruxolitinib 8.33 mM | Ruxolitinib 2.78 mM | Ruxolitinib 0.93 mM |

| | 1 | 2 | 3 | 4 | 5 | 6 | 7 | 8 | 9 | 10 | 11 | 12 | 13 | 14 | 15 | 16 | 17 | 18 | 19 | 20 | 21 | 22 | 23 | 24 |
|---|---|---|---|---|---|---|---|---|---|---|---|---|---|---|---|---|---|---|---|---|---|---|---|---|
| A | DMSO 100% | | | | | | | | | | | | | | | | | | | | | | | |
| B | BzCl 100 mM | | | | | | | | | | | | | | | | | | | | | | | |
| C | | | | | | | | | | | | | | | | | | | | | | | | |
| D | | | | | | | | | | | | | | | | | | | | | | | | |
| E | | | | | | | | | | | | | | | | | | | | | | | | |
| F | | | | | | | | | | | | | | | | | | | | | | | | |
| G | | | | | | | | | | | | | | | | | | | | | | | | |
| H | | | | | | | | | | | | | | | | | | | | | | | | |
| I | | | | | | | | | | | | | | | | | | | | | | | | |
| J | | | | | | | | | | | | | | | | | | | | | | | | |
| K | | | | | | | | | | | | | | | | | | | | | | | | |
| L | | | | | | | | | | | | | | | | | | | | | | | | |
| M | | | | | | | | | | | | | | | | | | | | | | | | |
| N | | | | | | | | | | | | | | | | | | | | | | | | |
| O | | | | | | | | | | | | | | | | | | | | | | | | |
| P | | | | | | | | | | | | | | | | | | | | | | | | |

| | 1 | 2 | 3 | 4 | 5 | 6 | 7 | 8 | 9 | 10 | 11 | 12 |
|---|---|---|---|---|---|---|---|---|---|---|---|---|
| A | | | | | | | | | | | | |
| B | | TIP 3.571<br>IBR 50 | TRA 0<br>BOS 0.222 | RUX 2.778<br>YM1 0 | SNS 0.204<br>VEN 1.875 | DJ3 0.625<br>QUI 1.25 | VEM 0.75<br>GDC 25 | GSK 0.75<br>DAU 0.075 | SGC 6.25<br>RO5 5 | RO5 5<br>DEC 5 | VEM 0.75<br>NSC 30 | PAN 0.002<br>OLA 0.4 |
| C | | APO 0.156<br>BOS 2 | IDE 7.5<br>IBR 12.5 | GIL 5<br>LEN 100 | VEN 0.469<br>AMS 5 | RUX 8.333<br>MET 0.009 | CHL 2.5<br>IDE 7.5 | LY2 0.625<br>CHL 40 | APO 0.625<br>LEN 25 | TOR 0.002<br>ATR 1.562 | SB2 3.125<br>BAR 0.056 | MK2 0.234<br>CHI 5 |
| D | | SAP 0.4<br>RO5 20 | APO 0.156<br>AG2 50 | PON 0.5<br>DAU 0.075 | WP1 0.556<br>SAP 10 | LAP 0.781<br>VEN 30 | VEN 0.469<br>CLA 0.2 | MET 0.002<br>ENT 1 | GAN 0.1<br>DMSO | BAR 0.056<br>MI4 10 | BOR 0<br>WM8 25 | RUX 25<br>AZA 4.167 |
| E | | DEX 6.25<br>CYT 5 | MK2 0.938<br>DMSO | TIP 0.073<br>ENT 20 | GAN 0.002<br>YM1 0.001 | CHL 0.625<br>CLA 0.2 | OLA 0.4<br>GDC 25 | ENT 2.778<br>PON 2 | RO5 1.25<br>SNS 0.204 | AMS 0.139<br>WP1 1.667 | NSC 7.5<br>CYT 1 | GSK 1.875<br>SNS 10 |
| F | | RUX 8.333<br>AMS 0.833 | GAL 3.704<br>ENT 2.778 | BOS 2<br>VOL 2 | MI4 0.156<br>VOL 2 | DEX 6.25<br>PAN 0.2 | SNS 1.429<br>SAP 50 | TIP 3.571<br>IRA 10 | GAN 0.006<br>LEN 25 | RUX 25<br>IRA 10 | DAU 0.019<br>IBR 12.5 | AMS 0.023<br>GAL 33.333 |
| G | | ENT 2.778<br>BOS 6 | LAP 50<br>CCT 2 | MI4 0.156<br>ENT 2 | AVA 1.562<br>IDE 1.875 | CCT 0.08<br>ABE 20 | BAR 0.056<br>CYT 0.2 | SAP 0.4<br>CCT 0.016 | LY2 2.5<br>DAU 0.075 | AG2 0.781<br>TOR 0.06 | FK5 50<br>MET 0.15 | SAP 10<br>TOR 10 |
| H | | LY2 2.5<br>AMS 0.833 | AG2 3.125<br>YM1 0.001 | SB2 0.781<br>DEC 5 | OLA 50<br>ENT 5 | SNS 0.029<br>QUI 1.25 | DJ3 0.156<br>GAN 0.006 | LY2 0.625<br>WP1 5 | IBR 0.781<br>YM1 0.001 | EC7 0.005<br>ENT 1 | CCT 0.08<br>SGC 25 | CCT 0.08<br>GSK 0.75 |
| I | | APO 2.5<br>MOC 1.25 | ABE 0.8<br>DEX 25 | SB2 0.781<br>TIP 0.073 | SB2 0.781<br>GSK 7.5 | AZA 0.116<br>DEX 100 | APO 0.625<br>GDC 100 | LEN 25<br>MET 0.038 | CHI 1.25<br>IBR 3.125 | IRA 0.625<br>CLA 0.2 | SAP 2<br>MK2 0.938 | WM8 12.5<br>FK5 1.852 |
| J | | AZA 0.694<br>CHI 20 | GSK 0.469<br>GAN 0.025 | IDE 1.875<br>CHI 1.25 | NSC 1.875<br>MOC 5 | AMS 0.139<br>VOL 2 | APO 0.625<br>AZA 25 | QUI 0.078<br>IRA 2.5 | AZA 0.694<br>OLA 50 | TIP 0.073<br>DEC 5 | PON 0.002<br>GIL 5 | LAP 12.5<br>CYT 5 |
| K | | DAU 0.019<br>GAN 0.1 | GAL 33.333<br>MOC 5 | VOL 0.056<br>CLA 0.2 | ABE 4<br>DMSO | NSC 30<br>VEN 30 | IBR 3.125<br>BAR 0.167 | GAN 0.025<br>SB2 12.5 | ABE 0.8<br>CHI 20 | NSC 1.875<br>ATR 25 | TOR 0.002<br>TRA 0.002 | VOL 2<br>DEC 5 |
| L | | PON 0.125<br>VOR 100 | LAP 3.125<br>WM8 50 | YM1 0<br>AMS 0.023 | ENT 2.778<br>PON 0.5 | TRA 0.002<br>EC7 1 | TIP 0.073<br>FK5 50 | LAP 0.781<br>IRA 0.156 | SNS 0.029<br>APO 10 | BOS 0.222<br>LEN 6.25 | VOL 0.009<br>DAU 0.019 | BOS 6<br>DMSO |
| M | | TOR 0.06<br>GSK 3 | DEX 1.562<br>VEN 0.469 | SNS 0.029<br>VEN 7.5 | WM8 100<br>PON 2 | WM8 12.5<br>MK2 0.938 | LY2 0.156<br>SGC 1.562 | VEM 0.188<br>DAU 0.075 | DEX 6.25<br>AMS 5 | TOR 0.002<br>GDC 100 | CLA 0.2<br>LY2 0.625 | MI4 0.156<br>SAP 10 |
| N | | ABE 0.16<br>SGC 25 | TOR 0.3<br>GDC 100 | CLA 0.04<br>IDE 7.5 | LY2 0.625<br>PAN 0.04 | ABE 0.16<br>VOR 1.562 | TIP 0.51<br>VOR 25 | GAN 0.1<br>GIL 5 | GSK 0.047<br>CHI 5 | ENT 0.2<br>GAL 100 | GSK 1.875<br>GAN 0.1 | GIL 1.25<br>GSK 3 |
| O | | IDE 0.469<br>TOR 0.06 | SNS 0.029<br>CLA 1 | SNS 0.204<br>GIL 1.25 | CHL 2.5<br>SGC 25 | IDE 0.469<br>WP1 5 | GIL 0.312<br>PAN 0.2 | APO 0.625<br>DMSO | RO5 5<br>VOR 100 | IDE 0.469<br>EVE 1 | ENT 2.778<br>SGC 25 | SGC 1.562<br>PAN 0.2 |
| P | | QUI 0.078<br>BAR 0.167 | WP1 1.667<br>FK5 50 | WM8 50<br>ABE 4 | ENT 0.926<br>VEM 0.75 | OLA 2<br>DAU 0.075 | ATR 1.562<br>TOR 0.06 | AVA 1.562<br>RO5 20 | DMSO | DAU 0.005<br>WP1 5 | TIP 0.51<br>EC7 1 | EC7 0.028<br>ABE 0.8 |
| Q | | VOL 0.056<br>WM8 100 | IDE 1.875<br>VOR 6.25 | NSC 1.875<br>CHI 1.25 | EC7 0.028<br>MOC 5 | APO 0.625<br>CCT 2 | SB2 12.5<br>SGC 6.25 | PAN 0.008<br>RUX 25 | GDC 1.562<br>LAP 50 | MI4 0.625<br>VEN 7.5 | BAR 0.056<br>PON 0.125 | EVE 0.01<br>WP1 1.667 |
| R | | AG2 12.5<br>LAP 50 | VEM 0.75<br>DJ3 10 | TRA 0<br>MK2 0.938 | IBR 0.781<br>MOC 5 | EVE 0.01<br>BOR 0.03 | CCT 0.016<br>LY2 0.625 | WP1 0.556<br>MK2 0.938 | GIL 5<br>CYT 5 | BOS 2<br>MOC 5 | NSC 0.469<br>BOS 0.222 | AVA 1.562<br>EVE 0.01 |
| S | | MET 0.009<br>AZA 25 | DEC 0.185<br>EC7 0.028 | DEC 0.185<br>CLA 1 | GIL 0.312<br>MI4 10 | WP1 0.185<br>IRA 2.5 | GAL 3.704<br>BOS 6 | PON 0.125<br>DMSO | MOC 0.078<br>AG2 3.125 | WM8 100<br>AVA 100 | PAN 0.002<br>RO5 1.25 | AZA 4.167<br>FK5 16.667 |
| T | | AG2 0.781<br>CCT 0.016 | BzCl | PON 0.031<br>DAU 0.3 | RO5 20<br>VOL 2 | GSK 1.875<br>DEX 25 | RUX 2.778<br>CLA 1 | TOR 0.002<br>RO5 20 | FK5 16.667<br>IRA 10 | RUX 75<br>BOR 0.03 | | VOL 0.333<br>CYT 5 |
| U | | CHI 5<br>LEN 100 | SAP 2<br>LAP 50 | SNS 1.429<br>AZA 25 | OLA 0.4<br>FK5 5.556 | MI4 2.5<br>GSK 3 | GAN 0.002<br>CCT 0.4 | BAR 0.056<br>YM1 0.02 | GSK 1.875<br>RO5 5 | MK2 15<br>EVE 10 | EVE 1<br>AG2 50 | TOR 0.002<br>VOR 25 |
| V | | ATR 1.562<br>CHI 20 | SAP 10<br>LAP 12.5 | ENT 5<br>CLA 5 | AVA 25<br>VOR 25 | GSK 7.5<br>SAP 50 | DEC 0.185<br>GSK 3 | ABE 20<br>VOR 100 | SAP 100 | PAN 0.04<br>CHI 20 | DAU 0.005<br>BOR 0.03 | WP1 0.185<br>AMS 0.833 |
| W | | SGC 0.391<br>CHI 1.25 | ENT 20<br>DMSO | CYT 0.04<br>APO 0.625 | GAN 0.006<br>QUI 0.312 | LY2 2.5<br>MK2 15 | OLA 0.4<br>AMS 0.023 | SGC 0.391<br>AMS 0.833 | MOC 0.312<br>NSC 7.5 | CCT 0.016<br>TIP 25 | WM8 50<br>BOS 2 | GDC 1.562<br>DAU 0.075 |
| X | | YM1 0.005<br>MET 0.15 | YM1 0.02<br>DEC 5 | CHL 2.5<br>MK2 15 | BOS 0.667<br>IRA 2.5 | TRA 0<br>GSK 0.469 | AZA 0.694<br>CHL 40 | AMS 0.023<br>VEN 7.5 | MOC 1.25<br>NSC 30 | TOR 0.012<br>CYT 1 | APO 0.625<br>EC7 1 | BzCl |
| Y | | MET 0.002<br>OLA 50 | MI4 0.625<br>LY2 2.5 | GDC 25<br>ATR 100 | TRA 0.1 | FK5 1.852<br>MOC 1.25 | AVA 25<br>DMSO | GSK 7.5<br>VEM 3 | NSC 7.5<br>MOC 5 | DEC 0.556<br>VEN 30 | SNS 0.029<br>DMSO | CCT 0.016<br>LAP 50 |
| Z | | PAN 0.04<br>DMSO | VOR 1.562<br>TRA 0.01 | RUX 25<br>VOL 0.333 | DEX 6.25<br>MK2 3.75 | ENT 8.333<br>VOL 2 | YM1 0<br>LEN 6.25 | ATR 1.562<br>NSC 7.5 | SAP 10<br>IBR 12.5 | PON 0.125<br>NSC 30 | AVA 25<br>SNS 10 | DEX 6.25<br>NSC 1.875 |
| AA | | MI4 0.156<br>YM1 0.001 | RUX 2.778<br>SB2 50 | WM8 100<br>SNS 10 | PON 0.125<br>ATR 25 | RUX 8.333<br>VEN 1.875 | MET 0.002<br>VEM 3 | TIP 3.571<br>BOS 6 | GAN 0.006<br>EC7 0.028 | CHI 1.25<br>MOC 0.312 | SGC 25<br>LY2 10 | DEC 0.556<br>VEN 30 |
| AB | | CYT 0.2<br>DAU 0.3 | ENT 0.926<br>GAL 33.333 | ENT 0.04<br>DEX 6.25 | DMSO | TIP 0.073<br>MI4 2.5 | VEM 0.047<br>ABE 20 | WM8 25<br>OLA 2 | WP1 1.667<br>VOL 2 | SB2 50<br>AZA 25 | ABE 0.16<br>OLA 2 | TIP 0.51<br>SAP 50 |
| AC | | BOS 2<br>OLA 50 | PON 0.031<br>GSK 0.047 | AG2 0.781<br>BAR 0.019 | LY2 0.625<br>CHL 10 | IBR 12.5<br>LEN 25 | APO 2.5<br>GSK 30 | WM8 12.5<br>DAU 0.075 | GAL 3.704<br>TRA 0.002 | GDC 1.562<br>BOS 0.667 | VOL 0.056<br>QUI 0.312 | PAN 0.2<br>CHI 20 |
| AD | | AG2 3.125<br>TRA 0.01 | ENT 0.2<br>SAP 50 | CHL 0.625<br>AMS 0.023 | SGC 0.391<br>DAU 0.019 | VOL 0.056<br>CLA 5 | SGC 0.391<br>VEN 30 | CHL 10<br>DEX 25 | MET 0.009<br>DMSO | CCT 0.016<br>BOR 0.03 | LY2 0.156<br>PON 0.031 | BOS 0.667<br>TOR 0.3 |
| AE | | CCT 0.4<br>LEN 25 | ENT 0.2<br>VEM 3 | GIL 0.312<br>BAR 0.056 | IBR 50<br>GSK 30 | APO 2.5<br>PON 2 | GAL 33.333<br>DJ3 10 | WP1 0.556<br>GAL 33.333 | VEM 0.75<br>MI4 10 | MOC 0.078<br>WP1 1.667 | GSK 0.188<br>ENT 20 | TRA 0.05<br>TIP 25 |
| AF | | | | | | | | | | | | |

| | 13 | 14 | 15 | 16 | 17 | 18 | 19 | 20 | 21 | 22 | 23 | 24 |
|---|---|---|---|---|---|---|---|---|---|---|---|---|
| A | | | | | | | | | | | | |
| B | CYT 0.2<br>TOR 0.06 | MK2 15<br>CCT 2 | SB2 50<br>AVA 100 | VOL 0.009<br>BAR 0.019 | IBR 3.125<br>WP1 1.667 | DEX 6.25<br>WP1 1.667 | EC7 0.005<br>QUI 5 | BAR 0.056<br>LY2 2.5 | IRA 0.625<br>RO5 1.25 | GSK 30<br>SNS 10 | WM8 50<br>FK5 16.667 | VOL 0.009<br>DEC 0.556 |
| C | VEM 0.188<br>WM8 50 | SB2 50<br>OLA 50 | TIP 0.073<br>SAP 50 | MET 0.002<br>TIP 3.571 | MK2 0.234<br>OLA 10 | RUX 8.333<br>CCT 0.4 | AG2 0.781<br>CHL 40 | TIP 0.51<br>CYT 1 | WP1 0.185<br>GAN 0.025 | ENT 0.926<br>GAN 0.025 | MOC 0.078<br>OLA 50 | OLA 0.4<br>CHI 0.312 |
| D | IBR 3.125<br>MI4 2.5 | LY2 0.625<br>BAR 0.5 | GSK 0.469<br>TIP 25 | DJ3 0.156<br>PAN 0.04 | IBR 3.125<br>SNS 0.204 | ENT 0.04<br>GSK 0.188 | TIP 0.073<br>DEC 0.185 | GIL 0.312<br>ENT 2.778 | WM8 12.5<br>MET 0.15 | VOL 0.056<br>EVE 1 | LY2 0.156<br>ENT 20 | SNS 1.429<br>DEX 100 |
| E | SGC 1.562<br>ABE 0.8 | TRA 0<br>BOS 0.667 | LY2 0.156<br>SAP 2 | IDE 1.875<br>LY2 10 | DAU 0.005<br>DEX 25 | GIL 1.25<br>PAN 0.2 | IRA 0.625<br>TIP 25 | AG2 3.125<br>LAP 50 | APO 0.625<br>DEX 25 | MI4 2.5<br>VOR 100 | MI4 0.156<br>IDE 1.875 | ENT 0.2<br>VOR 25 |
| F | IDE 1.875<br>VEN 7.5 | VEN 1.875<br>SNS 1.429 | YM1 0.005<br>FK5 50 | GIL 0.078<br>MET 0.002 | DJ3 2.5<br>QUI 1.25 | VOR 1.562<br>ATR 6.25 | GAL 3.704<br>SAP 50 | APO 0.625<br>MI4 2.5 | LY2 2.5<br>LAP 50 | CLA 1<br>CHL 40 | GAL 3.704<br>LY2 0.625 | CHI 0.312<br>AZA 25 |
| G | VOL 0.009<br>BOR 0.008 | LAP 3.125<br>TOR 0.3 | SAP 0.4<br>AVA 25 | DAU 0.005<br>IDE 30 | YM1 0<br>FK5 16.667 | BAR 0.019<br>CHL 10 | CCT 0.016<br>FK5 16.667 | EC7 0.028<br>CCT 0.4 | LEN 1.562<br>SB2 50 | OLA 2<br>PAN 0.04 | BOR 0.002<br>AVA 6.25 | GDC 100<br>PON 2 |
| H | MET 0.002<br>TRA 0.05 | SNS 0.204<br>IBR 50 | ABE 0.8<br>MI4 10 | GDC 1.562<br>DEX 25 | MK2 0.234<br>GAL 100 | APO 2.5<br>RUX 75 | MOC 0.312<br>GSK 30 | WM8 50<br>CHL 40 | RUX 8.333<br>OLA 50 | IBR 3.125<br>SGC 6.25 | CHI 20<br>VOR 100 | PAN 0.04<br>AG2 12.5 |
| I | AMS 0.139<br>EVE 1 | SAP 2<br>GSK 3 | CYT 0.04<br>CHL 2.5 | ENT 0.04<br>EVE 10 | DJ3 0.625<br>AZA 4.167 | WP1 0.556<br>RO5 1.25 | LEN 25<br>IRA 10 | TIP 0.51<br>MOC 0.312 | VEN 1.875<br>GSK 3 | PAN 0.04<br>BAR 0.5 | VEN 1.875<br>DJ3 10 | LY2 10<br>QUI 5 |
| J | NSC 0.469<br>CLA 0.04 | SGC 6.25<br>ENT 8.333 | GDC 25<br>LY2 2.5 | LAP 0.781<br>GIL 5 | NSC 0.469<br>QUI 1.25 | DJ3 0.625<br>RUX 75 | DJ3 0.156<br>SB2 50 | WM8 12.5<br>EC7 0.028 | SB2 3.125<br>ENT 5 | OLA 10<br>EVE 10 | BOR 0.03<br>MK2 15 | AMS 0.139<br>DEC 5 |
| K | GAL 11.111<br>IBR 3.125 | VOR 1.562<br>GSK 0.469 | IRA 0.156<br>SNS 1.429 | OLA 0.4<br>IRA 0.156 | CHL 0.625<br>IDE 1.875 | AG2 0.781<br>WM8 100 | EC7 0.028<br>TRA 0.01 | AMS 0.833<br>EVE 1 | LY2 2.5<br>CHI 20 | PAN 0.04<br>TIP 3.571 | TIP 3.571<br>LAP 50 | PON 0.031<br>TIP 25 |
| L | VEN 7.5<br>OLA 50 | LAP 3.125<br>MOC 0.312 | YM1 0<br>ATR 6.25 | LAP 12.5<br>MI4 10 | TRA 0<br>APO 0.156 | MET 0.009<br>CYT 5 | GIL 1.25<br>AG2 12.5 | BOR 0<br>ABE 20 | VOR 6.25<br>PON 2 | SAP 10<br>GDC 25 | TRA 0.002<br>CHI 20 | LY2 10<br>MOC 5 |
| M | MOC 1.25<br>EVE 10 | ABE 4<br>GAN 0.1 | DEC 0.556<br>RO5 5 | OLA 2<br>AG2 3.125 | IDE 7.5<br>YM1 0.02 | TRA 0.002<br>VOL 2 | SGC 6.25<br>AG2 12.5 | SB2 12.5<br>ABE 4 | GSK 0.188<br>PON 2 | PAN 0.04<br>BOS 2 | IRA 0.625<br>AMS 5 | SGC 1.562<br>TRA 0.002 |
| N | WM8 25<br>EVE 10 | CHI 0.312<br>MI4 10 | GIL 1.25<br>ENT 1 | GAL 3.704<br>SNS 0.204 | LY2 0.156<br>MOC 0.078 | IDE 0.469<br>EVE 0.01 | AMS 0.023<br>MOC 1.25 | VOL 0.333<br>VEM 3 | SAP 0.4<br>MOC 0.078 | TIP 25<br>DMSO | AVA 6.25<br>LEN 25 | BOS 0.222<br>PAN 0.008 |
| O | DJ3 0.156<br>BAR 0.5 | AZA 0.116<br>MOC 0.312 | DAU 0.075<br>MOC 5 | WP1 0.185<br>LEN 1.562 | IRA 2.5<br>ENT 1 | IRA 10<br>EVE 10 | GSK 0.469<br>GAL 100 | RO5 0.312<br>EC7 0.028 | SAP 0.4<br>VOL 0.333 | RO5 1.25<br>APO 10 | BOR 0.002<br>PAN 0.2 | TIP 0.51<br>MOC 1.25 |
| P | MET 0.009<br>VEM 0.75 | TRA 0.002<br>VOR 25 | IBR 12.5<br>SAP 50 | RO5 0.312<br>DMSO | OLA 0.4<br>SB2 12.5 | OLA 50<br>IRA 10 | MOC 1.25<br>ENT 1 | BAR 0.167<br>VEN 7.5 | AMS 0.139<br>DAU 0.075 | DEC 0.185<br>ATR 25 | VEN 7.5<br>IBR 50 | MET 0.038<br>BOR 0.03 |
| Q | AVA 25<br>ABE 4 | GSK 1.875<br>BAR 0.167 | GAN 0<br>DMSO | APO 0.156<br>ENT 0.926 | MI4 2.5<br>FK5 50 | CYT 0.2<br>BAR 0.5 | CHI 5<br>DAU 0.3 | MK2 3.75<br>AZA 4.167 | GSK 1.875<br>MET 0.038 | AG2 12.5<br>ENT 1 | CHI 0.312<br>SNS 1.429 | AVA 25<br>BOS 6 |
| R | TRA 0<br>DMSO | DEX 1.562<br>VOR 25 | DEX 6.25<br>BOS 0.667 | DEX 6.25<br>MET 0.038 | CHI 0.312<br>SAP 50 | LY2 0.156<br>GSK 0.188 | DAU 0.019<br>SGC 6.25 | BAR 0.167<br>BOS 6 | BOS 2<br>GSK 0.75 | ENT 0.04<br>ATR 25 | SAP 10<br>VEM 0.75 | TOR 0.002<br>APO 0.156 |
| S | GAN 0.025<br>SAP 25 | GIL 0.312<br>SAP 10 | GSK 0.047<br>AVA 6.25 | DEX 6.25<br>IBR 50 | ABE 0.16<br>CHL 2.5 | YM1 0.02<br>IBR 50 | GSK 0.047<br>SGC 1.562 | EC7 0.167<br>FK5 50 | DEX 6.25<br>CHL 40 | SGC 0.391<br>DMSO | GAN 0.025<br>MK2 3.75 | TIP 25<br>ENT 5 |
| T | GDC 6.25<br>TOR 0.3 | TRA 0.002<br>ABE 0.8 | WP1 0.185<br>PAN 0.2 | SGC 1.562<br>RUX 75 | EVE 0.01<br>DEC 1.667 | CLA 10<br>CCT 2 | CCT 0.08<br>ATR 100 | GDC 1.562<br>BOS 2 | YM1 0.002<br>BAR 0.5 | TIP 0.51<br>WP1 5 | SGC 6.25<br>AG2 50 | IDE 7.5<br>CCT 2 |
| U | PON 0.125<br>RO5 5 | MI4 0.156<br>SAP 50 | MK2 0.938<br>GSK 7.5 | IDE 0.469<br>RO5 0.312 | GIL 0.312<br>OLA 10 | AVA 25<br>GSK 30 | EVE 0.1<br>LY2 10 | FK5 1.852<br>APO 0.156 | WM8 12.5<br>DEC 1.667 | GAL 33.333<br>ENT 5 | WM8 25<br>GSK 7.5 | LY2 0.625<br>SNS 1.429 |
| V | FK5 16.667<br>MET 0.038 | VOL 0.009<br>SNS 10 | TOR 0.002<br>VEN 0.469 | RUX 0.926<br>DMSO | AMS 0.833<br>LEN 100 | CHL 0.625<br>LY2 10 | LAP 3.125<br>APO 0.625 | APO 0.156<br>ENT 0.2 | IRA 10<br>DMSO | AVA 6.25<br>NSC 1.875 | MI4 2.5<br>GSK 5 | SB2 50<br>DEX 100 |
| W | VOR 1.562<br>LEN 1.562 | OLA 2<br>ATR 100 | NSC 0.469<br>PON 0.031 | TRA 0<br>AG2 0.781 | GSK 0.188<br>OLA 10 | SAP 0.4<br>DMSO | AMS 0.833<br>AZA 25 | PAN 0.2<br>VOL 2 | GSK 1.875<br>DJ3 10 | TIP 0.073<br>SB2 3.125 | CYT 5<br>CLA 5 | GSK 0.188<br>GIL 5 |
| X | CYT 5<br>VEN 30 | VEM 0.047<br>TRA 0.01 | GSK 0.469<br>DEC 0.556 | AMS 0.139<br>BAR 0.5 | | GAL 3.704<br>CYT 5 | GIL 0.078<br>SNS 1.429 | LAP 3.125<br>AMS 5 | DEX 6.25<br>RO5 20 | AZA 4.167<br>LY2 10 | PAN 0.04<br>IDE 30 | AVA 1.562<br>IDE 7.5 |
| Y | CLA 0.04<br>DEX 100 | EVE 0.1<br>AG2 50 | IBR 0.781<br>EVE 0.01 | APO 0.156<br>QUI 5 | ABE 0.8<br>GSK 0.188 | MOC 0.312<br>VOL 2 | APO 0.625<br>GAN 0.1 | GIL 0.078<br>PON 0.125 | LY2 0.156<br>MOC 2 | LAP 0.781<br>GDC 100 | DEC 0.185<br>IRA 0.625 | VOR 1.562<br>BAR 0.5 |
| Z | BzCl | BAR 0.019<br>BOS 0.222 | WM8 50<br>AG2 50 | MOC 0.078<br>GAL 33.333 | CCT 0.08<br>SB2 50 | OLA 2<br>DMSO | MK2 0.938<br>EC7 0.167 | RUX 8.333<br>DEX 6.25 | GAN 0.006<br>LY2 10 | AVA 1.562<br>SNS 1.429 | DEC 0.185<br>FK5 50 | ENT 8.333<br>PAN 0.2 |
| AA | FK5 1.852<br>DEC 1.667 | IBR 3.125<br>IDE 7.5 | CLA 0.2<br>VOR 100 | AG2 3.125<br>BOS 6 | SGC 1.562<br>VEN 7.5 | BAR 0.056<br>BOS 2 | RUX 2.778<br>ENT 0.04 | OLA 10<br>LEN 25 | YM1 0.005<br>PON 0.5 | NSC 0.469<br>DEC 1.667 | SGC 1.562<br>RO5 1.25 | LY2 0.625<br>DEC 5 |
| AB | ENT 0.926<br>BAR 0.5 | VOL 0.009<br>TIP 3.571 | OLA 0.4<br>PON 0.125 | PON 0.125<br>TRA 0.01 | SNS 1.429<br>MOC 5 | LEN 6.25<br>IRA 2.5 | NSC 7.5<br>RUX 75 | IDE 1.875<br>PON 2 | NSC 0.469<br>AMS 0.833 | GDC 100<br>EVE 10 | BOS 2<br>WP1 5 | FK5 1.852<br>MET 0.002 |
| AC | CCT 0.4<br>RO5 20 | EC7 0.005<br>FK5 5.556 | YM1 0.001<br>DAU 0.075 | APO 2.5<br>AZA 25 | ENT 0.926<br>NSC 30 | DMSO | RO5 5<br>MOC 5 | MI4 2.5<br>MET 0.15 | CHL 0.625<br>APO 0.625 | IBR 0.781<br>EC7 0.167 | IBR 0.781<br>WM8 100 | WP1 0.556<br>ENT 5 |
| AD | ABE 0.16<br>CCT 0.4 | | GIL 0.078<br>BAR 0.019 | LY2 2.5<br>SNS 10 | GAN 0.025<br>DMSO | NSC 60 | ENT 0.2<br>LEN 25 | BOR 0<br>AG2 0.781 | GAL 3.704<br>RUX 25 | IRA 10<br>CYT 5 | GAN 0.1<br>TIP 25 | ABE 0.16<br>EC7 0.028 |
| AE | SGC 1.562<br>ENT 8.333 | YM1 0.001<br>LAP 50 | DJ3 0.156<br>WP1 0.185 | CLA 0.04<br>SAP 50 | CHI 0.312<br>IDE 7.5 | BOS 0.222<br>LY2 2.5 | AZA 0.694<br>PAN 0.04 | AG2 0.781<br>MOC 0.078 | AMS 0.023<br>WP1 0.556 | SB2 12.5<br>OLA 50 | DEX 1.562<br>DEC 0.556 | BOR 0<br>CCT 0.016 |
| AF | | | | | | | | | | | | |

| | 25 | 26 | 27 | 28 | 29 | 30 | 31 | 32 | 33 | 34 | 35 | 36 |
|---|---|---|---|---|---|---|---|---|---|---|---|---|
| A | | | | | | | | | | | | |
| B | WP1 1.667<br>DEC 1.667 | MOC 0.078<br>DEX 100 | SNS 1.429<br>EVE 1 | PON 0.125<br>MOC 1.25 | TOR 0.012<br>CLA 0.2 | SB2 3.125<br>DEC 0.556 | GDC 6.25<br>GSK 7.5 | DJ3 0.156<br>EVE 10 | VOL 0.009<br>BOR 0.03 | ABE 20<br>DMSO | IBR 3.125<br>BOS 2 | VOL 2<br>BOS 6 |
| C | RUX 2.778<br>DEX 25 | MET 0.002<br>GAN 0.1 | CHL 2.5<br>ATR 6.25 | TOR 0.012<br>AZA 4.167 | ENT 0.926<br>GDC 100 | ATR 25<br>VOR 100 | SAP 0.4<br>APO 0.625 | ENT 0.2<br>BOS 0.667 | GSK 0.469<br>LY2 2.5 | CHI 0.312<br>RUX 8.333 | DAU 0.005<br>YM1 0.005 | AVA 1.562<br>TIP 3.571 |
| D | CHL 0.625<br>IRA 0.156 | MET 0.002<br>GIL 5 | GAL 3.704<br>PON 0.031 | GIL 0.312<br>WP1 5 | EC7 0.005<br>DJ3 2.5 | PON 0.031<br>SB2 12.5 | SGC 6.25<br>VEN 7.5 | TIP 0.073<br>NSC 30 | RUX 2.778<br>LEN 6.25 | RO5 5<br>TRA 0.05 | GSK 0.469<br>RUX 25 | GSK 1.875<br>GAL 100 |
| E | TIP 0.51<br>AZA 4.167 | PAN 0.2<br>TOR 0.3 | TOR 0.002<br>TIP 3.571 | GIL 1.25<br>VEN 30 | EVE 0.01<br>GAN 0.025 | PON 0.031<br>SNS 10 | GSK 0.75<br>TIP 25 | OLA 10<br>VOR 100 | AZA 0.694<br>LEN 100 | CLA 0.04<br>CCT 0.08 | ENT 0.04<br>FK5 16.667 | RUX 8.333<br>ATR 6.25 |
| F | IDE 1.875<br>RO5 20 | SGC 0.391<br>GIL 0.078 | PAN 0.008<br>GSK 1.875 | LEN 6.25<br>CCT 0.4 | GIL 0.078<br>DEX 6.25 | SAP 2<br>CHI 5 | CLA 1<br>QUI 5 | WP1 1.667<br>OLA 10 | NSC 0.469<br>CLA 0.2 | CYT 0.2<br>RO5 5 | VEM 0.047<br>TRA 0.002 | DEX 1.562<br>PON 0.125 |
| G | ABE 4<br>ATR 25 | ABE 16<br>SNS 0.029 | GAN 0.1<br>YM1 0.02 | APO 0.156<br>VEM 3 | CHL 40<br>MI4 10 | MI4 0.625<br>EC7 0.167 | ATR 6.25<br>CCT 0.08 | VOL 2<br>CLA 5 | | SB2 0.781<br>AZA 0.116 | IRA 0.156<br>SNS 0.204 | CHL 0.625<br>GSK 1.875 |
| H | ABE 0.16<br>DJ3 10 | SAP 100 | WM8 12.5<br>DMSO | IDE 7.5<br>LY2 10 | IDE 1.875<br>DEX 100 | BzCl | ENT 0.2<br>AG2 12.5 | LY2 0.156<br>SNS 0.204 | GSK 1.875<br>PAN 0.04 | ATR 1.562<br>EVE 0.01 | DMSO | BOS 0.667<br>CHI 20 |
| I | AMS 0.023<br>QUI 5 | AZA 0.116<br>SNS 0.204 | RO5 0.312<br>GAL 33.333 | BzCl | FK5 5.556<br>QUI 1.25 | DAU 0.075<br>GAL 100 | VOL 0.333<br>ENT 8.333 | ENT 0.926<br>CCT 0.08 | GSK 7.5<br>BAR 0.5 | MET 0.009<br>LAP 50 | GSK 0.188<br>GAL 100 | DEX 1.562<br>MI4 0.625 |
| J | OLA 50<br>DMSO | GAL 3.704<br>DJ3 10 | ATR 100<br>GSK 3 | LEN 1.562<br>ABE 0.8 | GSK 0.469<br>AZA 0.116 | CLA 0.2<br>AVA 100 | TIP 3.571<br>ABE 20 | BAR 0.019<br>SAP 2 | WM8 25<br>CLA 1 | WP1 0.185<br>VEN 1.875 | MET 0.038<br>YM1 0.02 | ENT 0.926<br>TOR 0.06 |
| K | CCT 0.08<br>AZA 4.167 | IBR 0.781<br>ATR 1.562 | WP1 0.556<br>GIL 1.25 | BzCl | IRA 0.156<br>AZA 0.694 | WM8 12.5<br>GIL 0.078 | SB2 0.781<br>CYT 0.2 | TIP 3.571<br>VEN 30 | BAR 0.167<br>DJ3 10 | LAP 0.781<br>GIL 1.25 | VEM 0.047<br>PAN 0.2 | VEM 0.188<br>CCT 0.08 |
| L | VEN 1.875<br>LAP 12.5 | VEM 0.75<br>IRA 10 | TIP 0.073<br>PAN 0.2 | EC7 0.005<br>BOS 6 | ABE 0.8<br>SB2 12.5 | BOS 0.222<br>CYT 0.04 | APO 0.156<br>EVE 10 | EVE 0.01<br>APO 2.5 | GDC 1.562<br>IBR 12.5 | LEN 25<br>YM1 0.02 | BOS 0.222<br>GAN 0.006 | GAL 3.704<br>TRA 0.01 |
| M | SAP 10<br>BOR 0.03 | EC7 0.028<br>MK2 0.938 | NSC 30<br>VOL 2 | TIP 0.51<br>SNS 1.429 | SAP 2<br>CCT 2 | SNS 0.029<br>SB2 3.125 | RUX 2.778<br>SGC 6.25 | DMSO | SB2 12.5<br>DAU 0.3 | MET 0.038<br>AG2 50 | YM1 0.001<br>BOR 0.03 | CHL 0.625<br>OLA 50 |
| N | CHL 0.625<br>AMS 0.833 | MI4 0.039<br>DMSO | SAP 50<br>ENT 20 | WM8 50<br>PON 2 | LY2 0.625<br>PON 0.125 | BOR 0.002<br>AMS 0.833 | SAP 0.4<br>VEM 0.75 | MK2 0.234<br>ABE 4 | ATR 1.562<br>QUI 0.312 | BOS 0.222<br>VEN 30 | LAP 3.125<br>APO 10 | IBR 3.125<br>MET 0.038 |
| O | GSK 1.875<br>ENT 5 | GDC 1.562<br>RO5 1.25 | ENT 8.333<br>VOR 100 | GIL 0.078<br>DEC 5 | GAN 0.002<br>MET 0.15 | BOR 0.002<br>IDE 7.5 | LY2 0.625<br>MK2 15 | SAP 2<br>SB2 3.125 | TRA 0.1 | GAN 0.006<br>IBR 3.125 | FK5 5.556<br>VOL 2 | CYT 0.2<br>FK5 16.667 |
| P | AG2 3.125<br>MET 0.038 | VEM 0.047<br>LAP 12.5 | IRA 0.625<br>EVE 10 | DEC 0.185<br>TIP 25 | DMSO | TRA 0.002<br>EC7 0.167 | YM1 0.001<br>SB2 50 | IBR 12.5<br>GAN 0.1 | CHI 0.312<br>WP1 0.556 | CLA 0.2<br>APO 10 | VEM 0.047<br>EC7 0.167 | BOS 0.667<br>LEN 25 |
| Q | WM8 12.5<br>RUX 75 | WM8 100<br>GAL 100 | APO 10<br>DEC 5 | AG2 3.125<br>AZA 0.694 | IRA 0.625<br>MET 0.038 | NSC 0.469<br>APO 10 | RUX 2.778<br>TRA 0.002 | QUI 0.312<br>NSC 7.5 | RO5 0.312<br>DJ3 2.5 | GSK 0.469<br>SNS 1.429 | LEN 6.25<br>SNS 1.429 | PON 0.5<br>MET 0.038 |
| R | LAP 0.781<br>SAP 50 | SNS 1.429<br>VOL 2 | GSK 1.875<br>RUX 25 | GAN 0.006<br>DMSO | CHI 5<br>GSK 7.5 | EVE 0.1<br>AMS 5 | EVE 0.1<br>MET 0.15 | IRA 2.5<br>VEN 30 | DEC 1.667<br>MET 0.038 | NSC 1.875<br>ATR 100 | BOR 0.002<br>AMS 0.139 | BOS 0.222<br>GSK 1.875 |
| S | VEM 3<br>CHL 40 | TOR 0.012<br>LAP 12.5 | QUI 0.078<br>BOR 0.008 | LEN 1.562<br>ENT 5 | MI4 2.5<br>AG2 50 | WM8 12.5<br>FK5 50 | TOR 0.002<br>AZA 0.694 | AG2 12.5<br>DMSO | ENT 0.2<br>MET 0.15 | DAU 0.005<br>GSK 0.188 | | LEN 1.562<br>SNS 0.204 |
| T | MET 0.002<br>EVE 1 | TOR 0.002<br>MI4 0.625 | MI4 0.625<br>EVE 0.1 | CHL 10<br>LAP 12.5 | RUX 2.778<br>LAP 50 | AZA 0.116<br>SB2 50 | PON 0.125<br>ABE 20 | GIL 0.078<br>ATR 1.562 | LAP 12.5<br>RO5 20 | AG2 3.125<br>ATR 6.25 | WP1 0.556<br>BOS 2 | SNS 0.029<br>CCT 0.08 |
| U | GAN 0.002<br>GIL 5 | FK5 16.667<br>WM8 100 | EC7 0.005<br>ATR 6.25 | ENT 0.2<br>ENT 8.333 | WM8 12.5<br>SNS 0.029 | WM8 25<br>NSC 7.5 | TRA 0.01<br>FK5 50 | ABE 0.8<br>APO 10 | RUX 8.333<br>QUI 0.312 | PAN 0.008<br>EVE 10 | GIL 0.078<br>PAN 0.2 | LAP 12.5<br>RO5 5 |
| V | ENT 0.04<br>RO5 20 | BAR 0.019<br>PAN 0.008 | MK2 3.75<br>DMSO | WM8 12.5<br>MI4 0.156 | AG2 3.125<br>VEN 7.5 | RUX 2.778<br>CCT 2 | DEC 0.185<br>GAL 33.333 | BOS 0.222<br>OLA 10 | WM8 50<br>TRA 0.05 | GSK 0.047<br>CCT 0.4 | DEX 1.562<br>FK5 5.556 | MK2 3.75<br>CHI 5 |
| W | VEN 0.469<br>LEN 25 | YM1 0<br>NSC 7.5 | AZA 4.167<br>GAN 0.1 | EC7 0.028<br>ATR 6.25 | AG2 3.125<br>AVA 100 | SAP 10<br>IRA 10 | AZA 0.694<br>TOR 0.3 | CLA 0.04<br>GDC 100 | EC7 0.028<br>RUX 25 | MI4 0.625<br>SGC 1.562 | PAN 0.002<br>GDC 1.562 | QUI 0.312<br>AZA 25 |
| X | VEM 0.047<br>GSK 3 | MK2 3.75<br>ATR 25 | APO 10<br>ENT 20 | ENT 20<br>DMSO | WM8 12.5<br>AVA 100 | GSK 0.75<br>VOR 100 | OLA 0.4<br>DAU 0.019 | LEN 1.562<br>DAU 0.005 | TOR 0.012<br>MI4 2.5 | DEC 1.667<br>GDC 100 | GIL 0.078<br>IRA 10 | MET 0.038<br>LAP 50 |
| Y | BOS 0.222<br>FK5 5.556 | BOS 0.667<br>RO5 20 | LEN 1.562<br>LY2 0.625 | MET 0.002<br>BOR 0.008 | ENT 2.778<br>MI4 0.156 | GAL 11.111<br>LY2 0.625 | VOR 6.25<br>YM1 0.005 | PON 0.031<br>YM1 0.02 | MK2 0.234<br>DAU 0.075 | VOR 1.562<br>RO5 5 | VOR 1.562<br>LY2 10 | DEC 0.556<br>APO 2.5 |
| Z | DAU 0.019<br>DMSO | VOL 0.009<br>BAR 0.056 | LY2 0.156<br>IBR 3.125 | PAN 0.04<br>GDC 25 | MK2 0.938<br>RUX 25 | FK5 1.852<br>MOC 0.312 | MI4 0.625<br>GSK 0.188 | GAL 3.704<br>VEN 1.875 | WM8 25<br>IBR 50 | TOR 0.06<br>PAN 0.2 | GSK 0.047<br>GIL 5 | GIL 0.312<br>VEN 1.875 |
| AA | DEC 0.556<br>BAR 0.5 | AVA 1.562<br>PON 0.125 | YM1 0.005<br>LEN 25 | ENT 0.2<br>DJ3 2.5 | CYT 0.04<br>EVE 0.1 | APO 0.625<br>LAP 50 | BzCl | LAP 0.781<br>GAN 0.006 | VOR 6.25<br>AVA 25 | CLA 1<br>DMSO | CLA 0.04<br>TIP 25 | DEC 0.185<br>AVA 25 |
| AB | BAR 0.019<br>ENT 8.333 | GAN 0.002<br>SAP 2 | RO5 1.25<br>SAP 50 | DEX 25<br>OLA 10 | DJ3 2.5<br>ATR 100 | PON 0.125<br>WP1 5 | VOR 6.25<br>SGC 25 | WM8 50<br>MK2 15 | ENT 20<br>AMS 5 | LEN 1.562<br>GDC 25 | RUX 2.778<br>RO5 5 | AZA 0.116<br>ENT 2.778 |
| AC | GDC 25<br>QUI 5 | LY2 0.156<br>VEN 7.5 | EC7 0.005<br>MET 0.15 | RUX 2.778<br>BOS 6 | WM8 100<br>ENT 5 | ATR 6.25<br>QUI 0.312 | DAU 0.005<br>QUI 1.25 | ENT 0.926<br>WM8 25 | GAL 200 | VOL 0.056<br>BOS 6 | LAP 50<br>MOC 5 | GAL 11.111<br>MET 0.15 |
| AD | CYT 0.2<br>TIP 3.571 | SB2 3.125<br>RO5 5 | VOR 1.562<br>GIL 1.25 | VOR 1.562<br>VEN 7.5 | GIL 1.25<br>EVE 10 | QUI 0.312<br>TIP 25 | PAN 0.04<br>LY2 10 | GDC 6.25<br>WP1 1.667 | DEC 0.556<br>LY2 10 | SGC 1.562<br>MK2 15 | PON 0.5<br>SNS 10 | TOR 0.012<br>BOS 0.667 |
| AE | WP1 0.185<br>DAU 0.3 | LAP 12.5<br>IDE 30 | CHL 2.5<br>VOR 100 | OLA 2<br>DMSO | VOR 25<br>WM8 100 | NSC 0.469<br>ABE 0.8 | DJ3 0.156<br>DAU 0.075 | CCT 0.016<br>DJ3 2.5 | SB2 0.195<br>DMSO | IBR 100 | LAP 3.125<br>CCT 2 | CYT 1<br>PON 2 |
| AF | | | | | | | | | | | | |

| | 37 | 38 | 39 | 40 | 41 | 42 | 43 | 44 | 45 | 46 | 47 | 48 |
|---|---|---|---|---|---|---|---|---|---|---|---|---|
| A | | | | | | | | | | | | |
| B | AVA 6.25<br>IRA 10 | NSC 0.469<br>ENT 20 | MI4 0.156<br>TRA 0.05 | LAP 100 | GAL 3.704<br>ENT 0.2 | NSC 1.875<br>BOS 0.667 | GSK 0.188<br>NSC 30 | CHI 5<br>BOS 6 | CLA 1<br>DMSO | ENT 0.926<br>CYT 0.04 | EC7 1<br>EVE 10 | |
| C | AZA 0.116<br>SB2 12.5 | SAP 50<br>DAU 0.3 | TRA 0<br>PAN 0.008 | GSK 1.875<br>IRA 2.5 | ABE 0.16<br>DMSO | DEC 1.562<br>OLA 2 | ABE 4<br>ATR 100 | DAU 0.075<br>ATR 100 | NSC 0.469<br>RO5 1.25 | TOR 0.002<br>DEX 25 | | |
| D | AMS 0.139<br>TRA 0.01 | FK5 5.556<br>GAN 0.1 | ENT 0.04<br>EC7 1 | MET 0.009<br>TIP 3.571 | TRA 0.002<br>CHI 20 | LAP 3.125<br>RO5 20 | TIP 3.571<br>WP1 5 | CCT 0.016<br>AG2 3.125 | PON 0.5<br>SB2 50 | LY2 0.156<br>DJ3 0.625 | GSK 0.188<br>DMSO | |
| E | BOR 0.008<br>EVE 1 | RO5 0.312<br>GDC 25 | TRA 0.002<br>CLA 5 | AZA 0.116<br>DJ3 0.625 | DJ3 2.5<br>EVE 1 | YM1 0<br>VEM 0.188 | DAU 0.005<br>FK5 50 | ENT 2.778<br>ENT 5 | LEN 6.25<br>WM8 100 | EC7 0.005<br>AMS 0.833 | EC7 0.028<br>OLA 50 | |
| F | DEX 25<br>LY2 10 | PON 2<br>DMSO | DAU 0.075<br>DEC 5 | GAL 100<br>DMSO | LAP 50<br>EC7 1 | DJ3 10<br>GDC 100 | EVE 10<br>VEN 30 | WP1 0.185<br>DEC 0.185 | LEN 6.25<br>NSC 7.5 | GIL 1.25<br>QUI 1.25 | GAN 0.002<br>NSC 7.5 | |
| G | GSK 12.5<br>TIP 25 | LY2 0.156<br>BAR 0.056 | MK2 3.75<br>CHI 20 | DEC 0.556<br>WM8 100 | CLA 0.2<br>ATR 25 | WP1 0.556<br>IDE 30 | FK5 16.667<br>MK2 15 | PON 2<br>ENT 5 | VOR 6.25<br>EVE 10 | IDE 1.875<br>GIL 2 | IDE 0.469<br>WM8 100 | |
| H | TOR 0.3<br>CYT 5 | SGC 1.562<br>FK5 5.556 | AZA 4.167<br>CLA 1 | CHL 10<br>SB2 12.5 | BAR 1 | VOR 6.25<br>CHL 5 | DMSO | EVE 1<br>CLA 5 | VOL 0.056<br>MOC 1.25 | ENT 2.778<br>CLA 0.2 | CHL 0.312<br>IDE 30 | |
| I | LAP 0.781<br>DEX 25 | OLA 2<br>RO5 1.25 | WM8 25<br>DEX 6.25 | RUX 2.778<br>GSK 0.188 | GSK 0.047<br>TOR 0.012 | AVA 1.562<br>GIL 0.078 | EC7 0.005<br>TIP 0.51 | DEC 0.185<br>DMSO | ENT 5<br>EVE 10 | VOR 100<br>DMSO | IBR 0.781<br>VEM 0.188 | |
| J | WM8 25<br>AMS 0.833 | WP1 0.185<br>CHI 20 | RUX 2.778<br>NSC 30 | SAP 0.4<br>BOS 0.667 | MOC 0.078<br>VOR 100 | WP1 5<br>SNS 10 | SB2 0.781<br>GAN 0.025 | IRA 2.5<br>SAP 50 | OLA 2<br>MET 0.15 | GAN 0.006<br>WM8 25 | SNS 0.204<br>TIP 3.571 | |
| K | AZA 0.116<br>DAU 0.019 | TRA 0.01<br>TIP 3.571 | EC7 0.028<br>MET 0.15 | WM8 25<br>DJ3 10 | RO5 5<br>QUI 1.25 | GAL 3.704<br>CYT 0.04 | ATR 1.562<br>QUI 1.25 | ABE 20<br>DEC 5 | RUX 2.778<br>AMS 0.833 | TRA 0<br>DJ3 2.5 | DJ3 0.625<br>IBR 50 | |
| L | ABE 0.8<br>GAL 11.111 | VEM 0.047<br>GAN 0.1 | IDE 1.875<br>CHI 5 | RUX 2.778<br>ENT 0.926 | EVE 0.01<br>VEN 1.875 | BOR 0.008<br>WM8 100 | AMS 0.023<br>BOR 0.008 | WM8 25<br>ENT 2.778 | LEN 1.562<br>BOR 0.008 | WP1 1.667<br>IBR 12.5 | GIL 0.078<br>AG2 50 | |
| M | RUX 2.778<br>ENT 20 | ENT 2.778<br>SNS 0.204 | GAL 11.111<br>PAN 0.04 | ABE 0.8<br>CHI 5 | SGC 1.562<br>GSK 1.875 | BAR 0.019<br>RUX 8.333 | SB2 0.781<br>NSC 1.875 | TOR 0.012<br>IDE 30 | CHI 1.25<br>SGC 6.25 | GAN 0.006<br>ATR 6.25 | CYT 0.2<br>BOR 0.03 | |
| N | GSK 0.469<br>ENT 2.778 | DEX 6.25<br>BAR 0.5 | OLA 0.4<br>APO 0.625 | GSK 3<br>DMSO | PAN 0.2<br>DMSO | BOR 0.008<br>AG2 12.5 | EC7 2 | AMS 0.023<br>APO 10 | TOR 0.002<br>BOS 0.222 | PON 0.031<br>SNS 1.429 | ABE 0.16<br>RO5 5 | |
| O | CLA 1<br>GAN 0.1 | MET 0.038<br>GIL 5 | VEM 0.75<br>ENT 5 | GDC 1.562<br>VEM 3 | TRA 0<br>VEN 30 | CYT 0.04<br>TIP 0.51 | VOR 6.25<br>QUI 0.312 | EC7 0.005<br>CCT 0.016 | LAP 0.781<br>ENT 1 | MOC 1.25<br>BOS 2 | ENT 0.04<br>IRA 2.5 | |
| P | IRA 2.5<br>VEN 7.5 | RO5 1.25<br>ENT 1 | CHL 10<br>BAR 0.167 | PON 0.125<br>AG2 50 | YM1 0<br>OLA 50 | SAP 0.4<br>BOR 0.002 | AG2 0.781<br>PON 2 | VOR 6.25<br>LY2 2.5 | QUI 1.25<br>VOL 2 | ENT 0.926<br>DEX 25 | YM1 0.001<br>CLA 0.2 | |
| Q | GAL 33.333<br>SNS 10 | EC7 0.028<br>CHL 10 | CHI 0.312<br>SNS 0.204 | ATR 6.25<br>SNS 1.429 | APO 0.625<br>RUX 75 | GAL 33.333<br>YM1 0.02 | APO 0.625<br>CHI 5 | EVE 1<br>GAN 0.1 | MI4 0.156<br>AVA 25 | CYT 0.2<br>CCT 2 | EVE 0.01<br>YM1 0.02 | |
| R | WM8 12.5<br>RO5 1.25 | GSK 0.469<br>AG2 50 | ATR 1.562<br>SGC 25 | YM1 0.001<br>AZA 0.694 | MOC 0.312<br>GAN 0.1 | DEX 25<br>NSC 30 | GSK 7.5<br>EVE 10 | CHI 5<br>VOR 100 | EC7 0.028<br>CLA 5 | SGC 0.391<br>DJ3 10 | AVA 1.562<br>TIP 0.073 | |
| S | ABE 4<br>IBR 50 | QUI 1.25<br>ATR 100 | RO5 1.25<br>AMS 0.139 | BOS 0.222<br>GIL 1.25 | YM1 0.02<br>GSK 3 | DEC 1.562<br>EC7 0.167 | CHL 0.625<br>TOR 0.3 | BzCl | PON 0.031<br>IBR 12.5 | LY2 10<br>AMS 5 | MK2 0.234<br>ATR 1.562 | |
| T | TOR 0.012<br>IRA 2.5 | EVE 0.01<br>VEM 0.188 | CHI 0.312<br>LAP 3.125 | MET 0.009<br>EVE 10 | FK5 1.852<br>LEN 6.25 | CHI 0.312<br>GSK 7.5 | LAP 0.781<br>CYT 0.04 | OLA 50<br>EVE 10 | ENT 8.333<br>LEN 25 | VOR 1.562<br>TIP 0.51 | VEM 0.188<br>CLA 5 | |
| U | GDC 1.562<br>GSK 0.047 | NSC 7.5<br>LEN 25 | GIL 0.078<br>SAP 2 | PON 0.125<br>YM1 0.005 | TIP 0.073<br>DAU 0.075 | IDE 7.5<br>WM8 100 | DEX 6.25<br>AG2 12.5 | DEX 1.562<br>MOC 1.25 | IDE 1.875<br>IRA 10 | RO5 5<br>YM1 0.02 | LAP 12.5<br>ATR 100 | |
| V | MK2 3.75<br>SNS 1.429 | CCT 0.08<br>GAN 0.025 | APO 0.156<br>AVA 25 | WM8 50<br>GAL 100 | YM1 0.001<br>GIL 5 | AVA 25<br>ATR 25 | CYT 0.2<br>AMS 0.833 | RO5 1.25<br>EC7 0.167 | AG2 12.5<br>MK2 15 | ABE 0.16<br>LEN 100 | GIL 0.078<br>DAU 0.005 | |
| W | LY2 0.156<br>FK5 5.556 | PON 0.5<br>DMSO | MK2 0.938<br>AG2 12.5 | PAN 0.002<br>EC7 0.167 | WP1 0.556<br>DEC 1.667 | AVA 6.25<br>GIL 5 | OLA 0.4<br>EVE 0.01 | | LY2 0.156<br>AZA 0.694 | GIL 0.312<br>DEC 5 | MI4 0.156<br>TOR 0.012 | |
| X | RUX 25<br>CLA 5 | IDE 0.469<br>GSK 0.469 | SAP 10<br>IDE 7.5 | IBR 0.781<br>MK2 0.938 | WM8 25<br>CCT 2 | LY2 0.156<br>ABE 20 | YM1 0<br>DEC 0.556 | LY2 0.625<br>PAN 0.2 | IBR 0.781<br>WP1 5 | MET 0.038<br>GAN 0.1 | SGC 25<br>DEX 100 | |
| Y | YM1 0.001<br>ENT 8.333 | | RO5 0.312<br>LAP 3.125 | MI4 0.625<br>MK2 15 | GSK 1.875<br>GIL 5 | IRA 0.625<br>QUI 5 | VEM 0.047<br>MET 0.15 | BOS 2<br>CLA 1 | LY2 0.156<br>IBR 50 | LY2 2.5<br>MK2 15 | MET 0.001<br>DMSO | |
| Z | DMSO | GDC 6.25<br>IRA 2.5 | ENT 8.333<br>LAP 50 | MOC 0.078<br>VEM 0.75 | BAR 0.019<br>CLA 1 | DAU 0.3<br>CYT 5 | NSC 1.875<br>VOR 6.25 | WM8 100<br>ATR 100 | TRA 0.002<br>VEN 1.875 | CLA 0.2<br>VOL 2 | AZA 0.694<br>MET 0.15 | |
| AA | MI4 0.156<br>TOR 0.002 | RUX 2.778<br>APO 2.5 | MI4 2.5<br>IDE 7.5 | GDC 1.562<br>DEC 0.556 | MI4 0.156<br>GSK 3 | BOS 0.222<br>VOR 6.25 | SGC 6.25<br>SNS 10 | GAL 33.333<br>LEN 100 | PON 0.031<br>TRA 0.05 | DAU 0.005<br>CLA 0.2 | SNS 10<br>CLA 5 | |
| AB | VOL 0.333<br>GSK 30 | PAN 0.2<br>VOR 100 | TIP 3.571<br>ENT 8.333 | AMS 0.023<br>DEC 5 | SAP 50<br>SNS 10 | DEX 6.25<br>GAL 100 | CYT 0.04<br>VEM 3 | ABE 4<br>SB2 50 | DEC 0.185<br>MET 0.038 | GAN 0.006<br>AZA 25 | MK2 3.75<br>WM8 100 | |
| AC | BAR 0.019<br>MOC 5 | GSK 1.875<br>ATR 25 | AZA 0.116<br>GSK 7.5 | SNS 1.429<br>PAN 0.2 | WP1 1.667<br>MOC 1.25 | PON 0.031<br>RO5 20 | QUI 0.312<br>DMSO | PON 0.5<br>SB2 50 | GIL 0.078<br>GAN 0.1 | PON 0.031<br>RUX 75 | DAU 0.019<br>APO 2.5 | |
| AD | GDC 6.25<br>ENT 5 | MI4 2.5<br>ENT 20 | IBR 3.125<br>DEC 1.667 | PAN 0.002<br>BAR 0.5 | FK5 1.852<br>YM1 0.005 | TIP 0.073<br>AZA 4.167 | GSK 0.469<br>AVA 100 | VEN 0.469<br>DEX 25 | CYT 0.2<br>AMS 5 | BOR 0.002<br>PAN 0.04 | DEC 1.667<br>IDE 30 | |
| AE | LAP 0.781<br>CHL 2.5 | EVE 0.1<br>GAL 33.333 | ABE 0.8<br>PON 0.125 | GDC 25<br>RO5 20 | SAP 0.4<br>CHL 10 | CYT 0.04<br>QUI 5 | CHI 0.312<br>LEN 25 | BAR 0.056<br>IDE 7.5 | BOS 0.222<br>ENT 2.778 | IRA 0.156<br>NSC 30 | CCT 0.08<br>SAP 50 | |
| AF | | | | | | | | | | | | |

| | | |
|---|---|---|
| qcdata | list [1] (S3: controlData) | List of length 1 |
| MeWo | list [3] | List of length 3 |
| data | list [332 x 14] (S3: data.fram | A data.frame with 332 rows and 14 columns |
| variance | list [3] | List of length 3 |
| data | list [3] | List of length 3 |
| BzCl | list [149 x 15] (S3: data.fram | A data.frame with 149 rows and 15 columns |
| DMSO | list [151 x 15] (S3: data.fram | A data.frame with 151 rows and 15 columns |
| Untreated | list [32 x 15] (S3: data.frame | A data.frame with 32 rows and 15 columns |
| boxplot | list [3] | List of length 3 |
| boxplot-byplate | list [3] | List of length 3 |
| z-factor | list [2] | List of length 2 |
| data | list [23 x 4] (S3: data.frame) | A data.frame with 23 rows and 4 columns |
| original | list [9] (S3: gg, ggplot) | List of length 9 |
