## Supplementary Figures (5-23) for "metascreen: A modular tool for the design and analysis of drug combination screens"

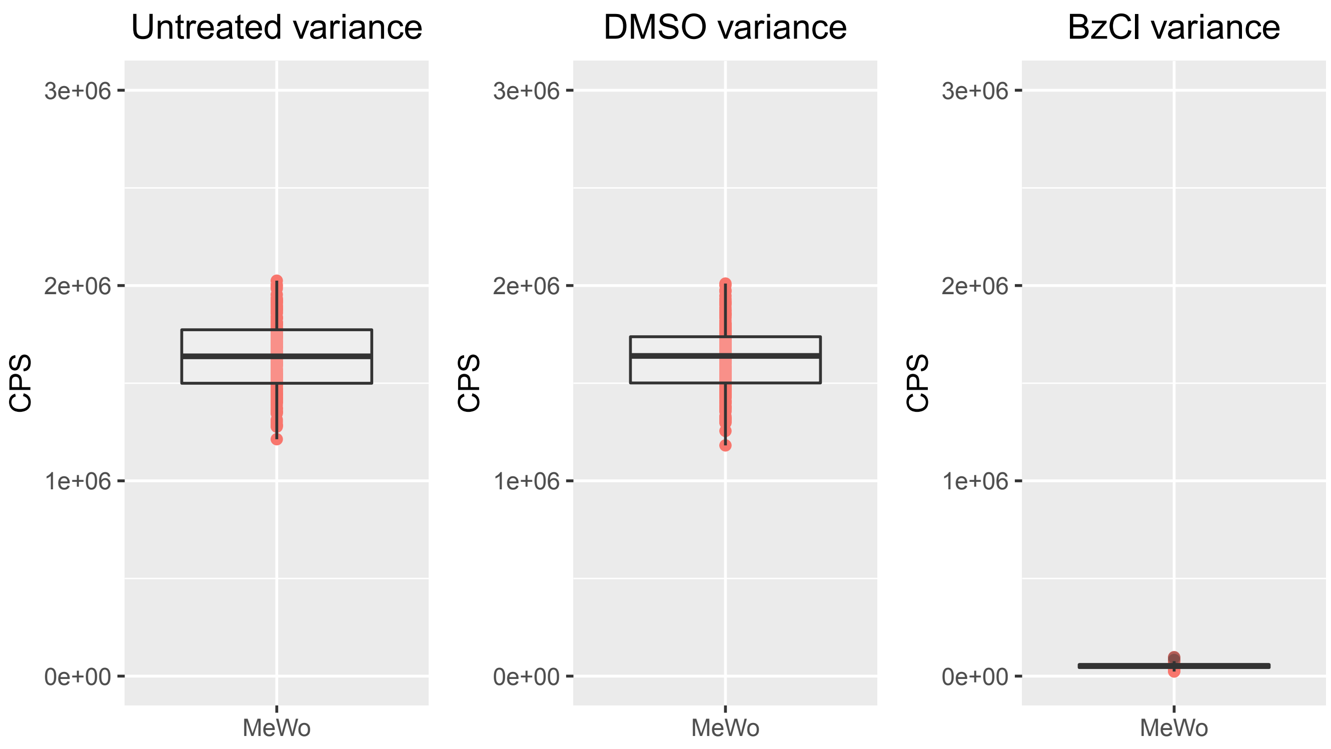
Untreated variance
DMSO variance
BzCl variance
CPS
3e+06
2e+06
1e+06
0e+00
MeWo

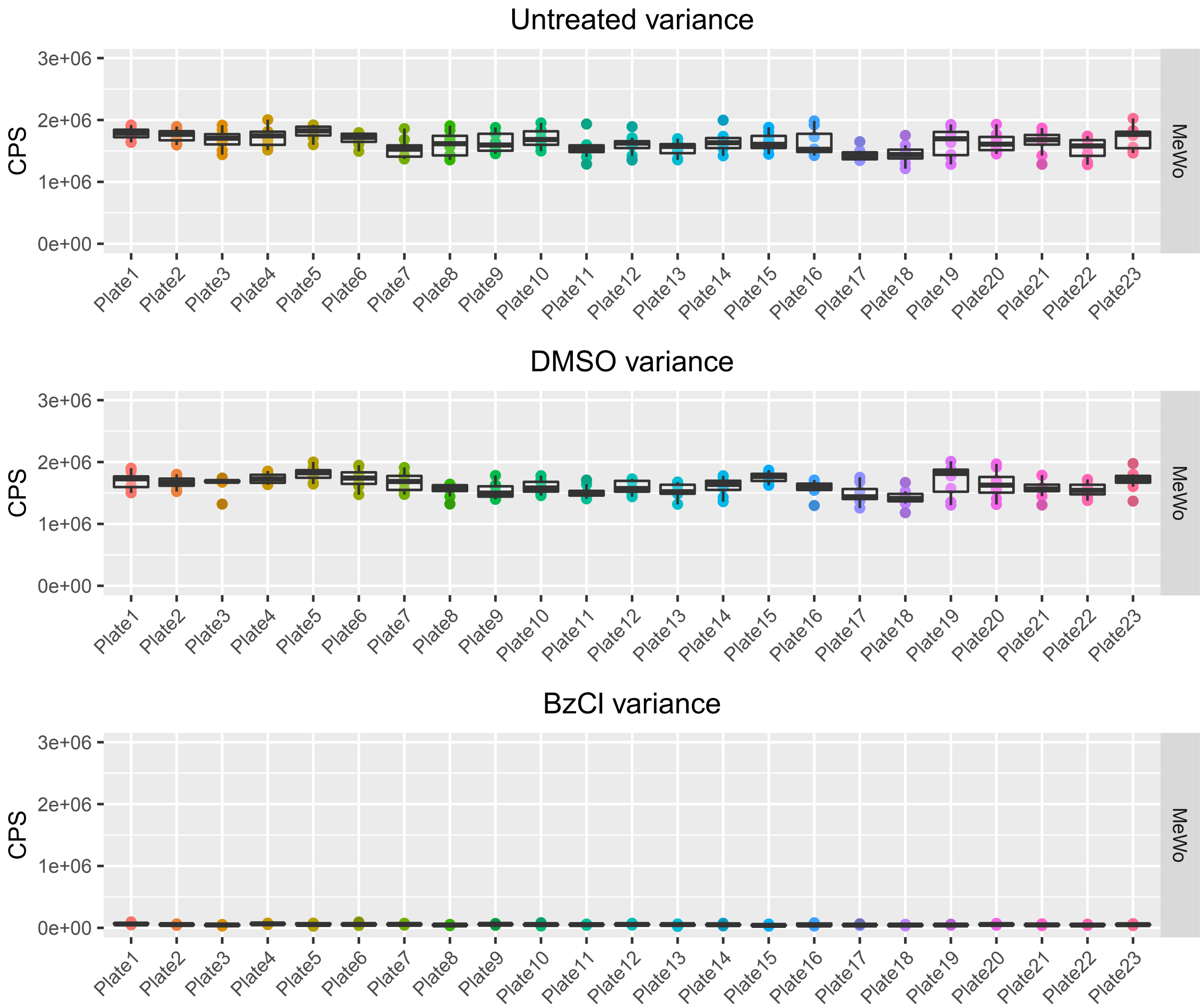
Untreated variance
CPS
3e+06
2e+06
1e+06
0e+00
MeWo
Plate1
Plate2
Plate3
Plate4
Plate5
Plate6
Plate7
Plate8
Plate9
Plate10
Plate11
Plate12
Plate13
Plate14
Plate15
Plate16
Plate17
Plate18
Plate19
Plate20
Plate21
Plate22
Plate23
DMSO variance
BzCl variance

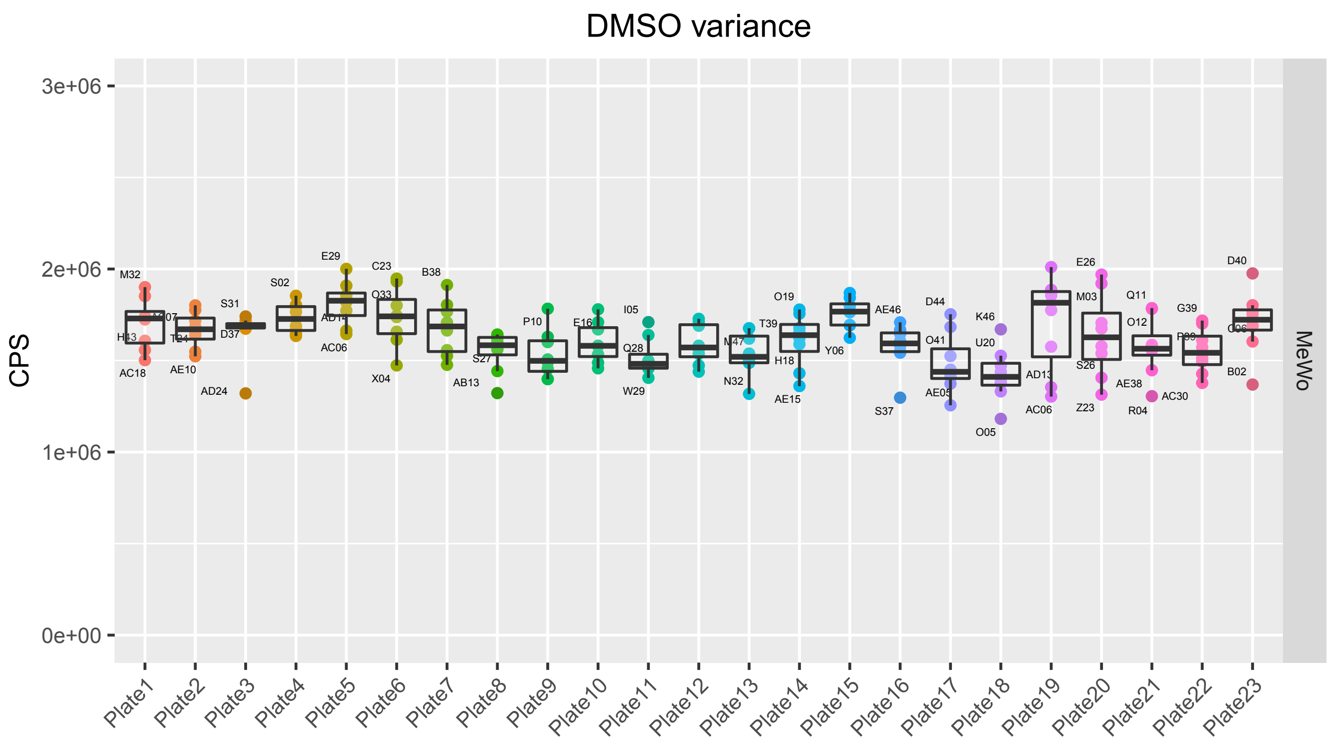

DMSO variance
CPS
0e+00
1e+06
2e+06
3e+06
Plate1
Plate2
Plate3
Plate4
Plate5
Plate6
Plate7
Plate8
Plate9
Plate10
Plate11
Plate12
Plate13
Plate14
Plate15
Plate16
Plate17
Plate18
Plate19
Plate20
Plate21
Plate22
Plate23
MeWo

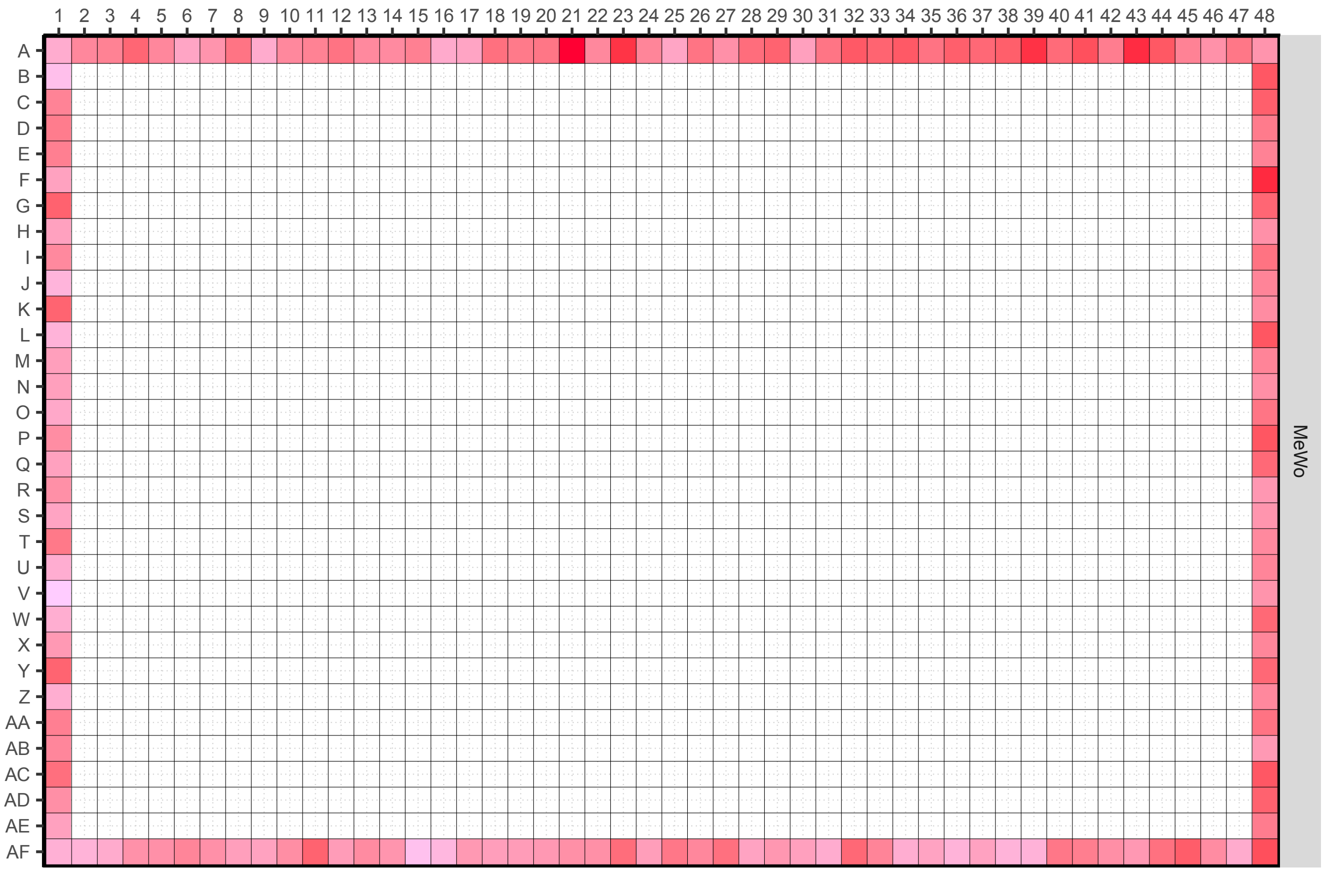

1 2 3 4 5 6 7 8 9 10 11 12 13 14 15 16 17 18 19 20 21 22 23 24 25 26 27 28 29 30 31 32 33 34 35 36 37 38 39 40 41 42 43 44 45 46 47 48
A B C D E F G H I J K L M N O P Q R S T U V W X Y Z AA AB AC AD AE AF
MeWo

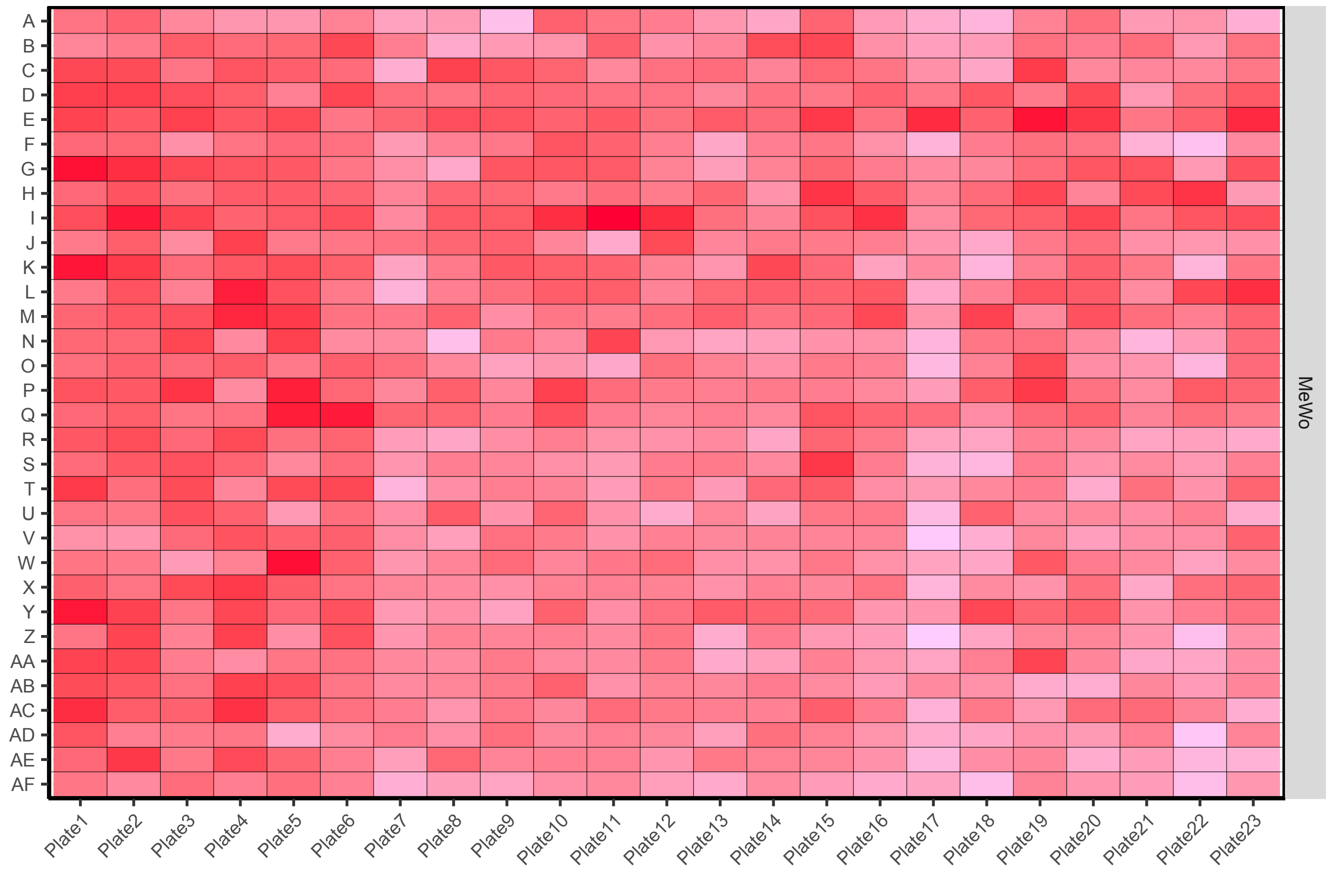
A
B
C
D
E
F
G
H
I
J
K
L
M
N
O
P
Q
R
S
T
U
V
W
X
Y
Z
AA
AB
AC
AD
AE
AF
Plate1
Plate2
Plate3
Plate4
Plate5
Plate6
Plate7
Plate8
Plate9
Plate10
Plate11
Plate12
Plate13
Plate14
Plate15
Plate16
Plate17
Plate18
Plate19
Plate20
Plate21
Plate22
Plate23
MeWo

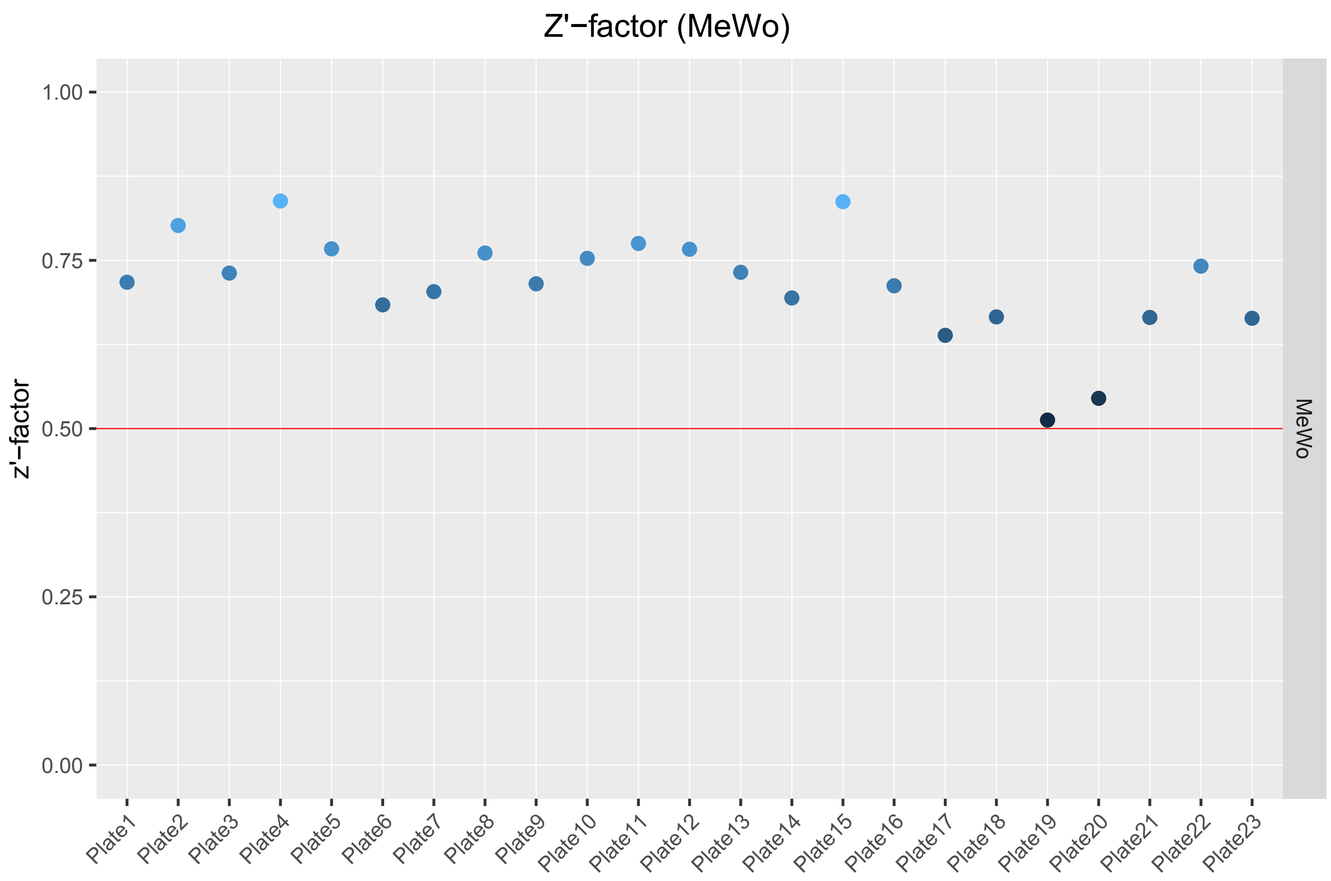

Z'−factor (MeWo)
z'−factor
1.00
0.75
0.50
0.25
0.00
MeWo
Plate1
Plate2
Plate3
Plate4
Plate5
Plate6
Plate7
Plate8
Plate9
Plate10
Plate11
Plate12
Plate13
Plate14
Plate15
Plate16
Plate17
Plate18
Plate19
Plate20
Plate21
Plate22
Plate23

### MeWo: Dose Response (Viability)

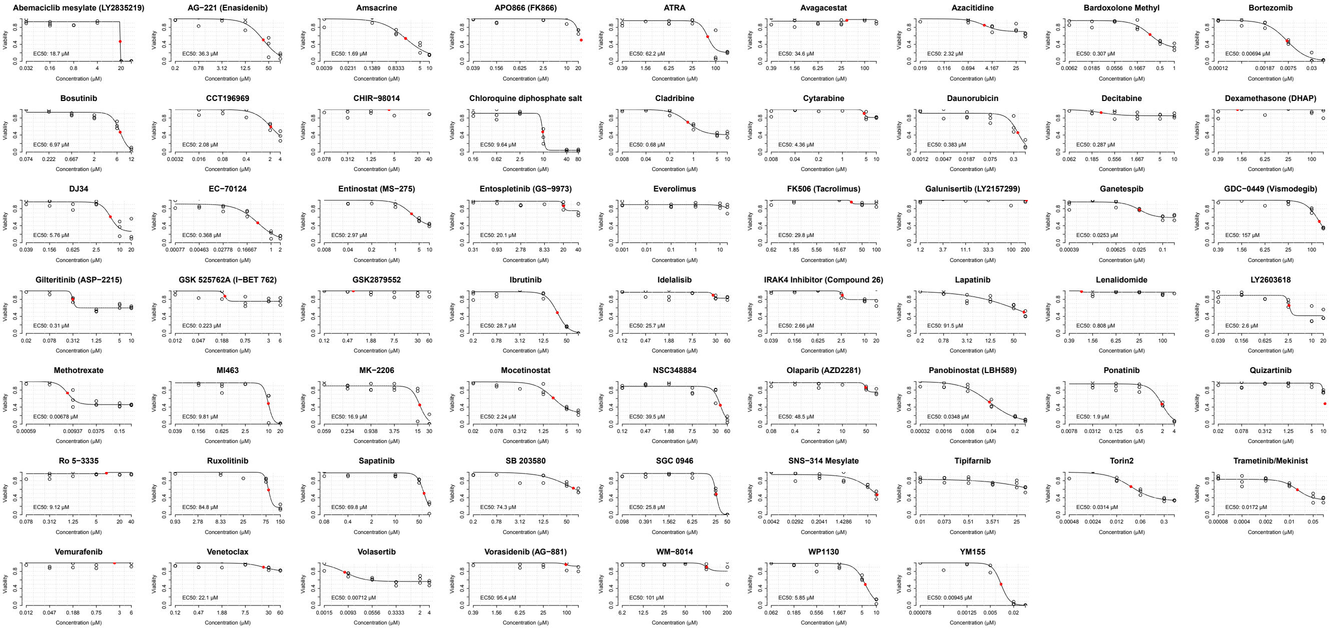

### MeWo: Dose Response (Inhibition)

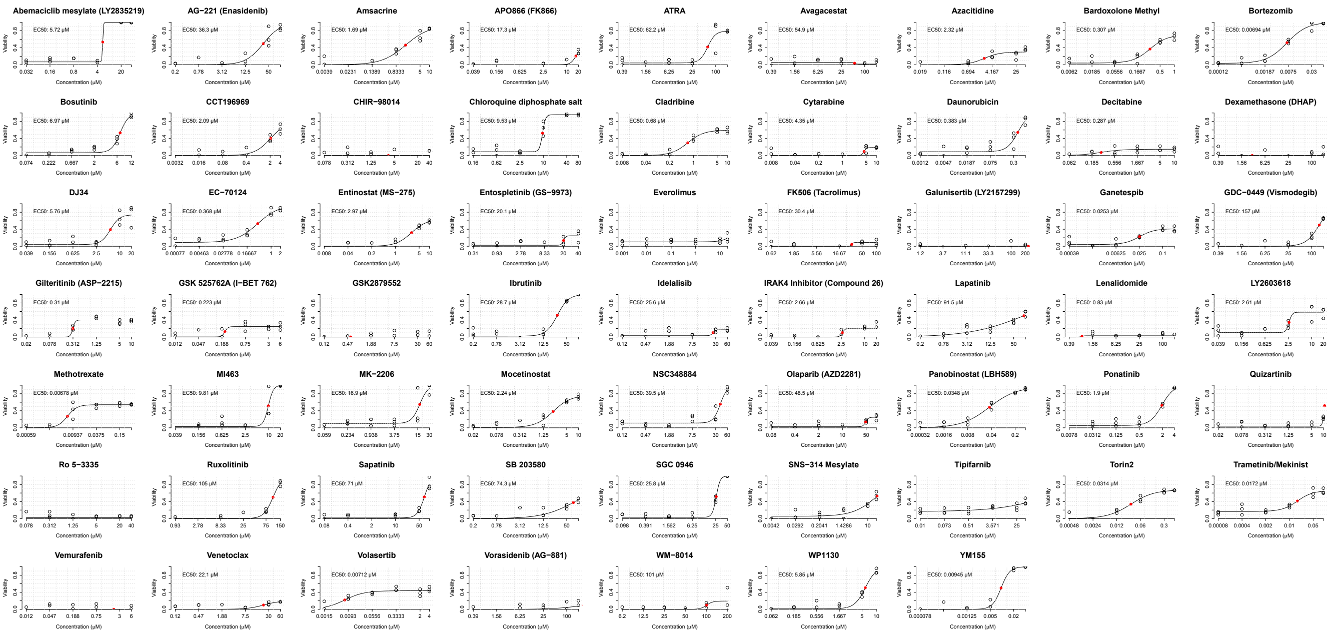

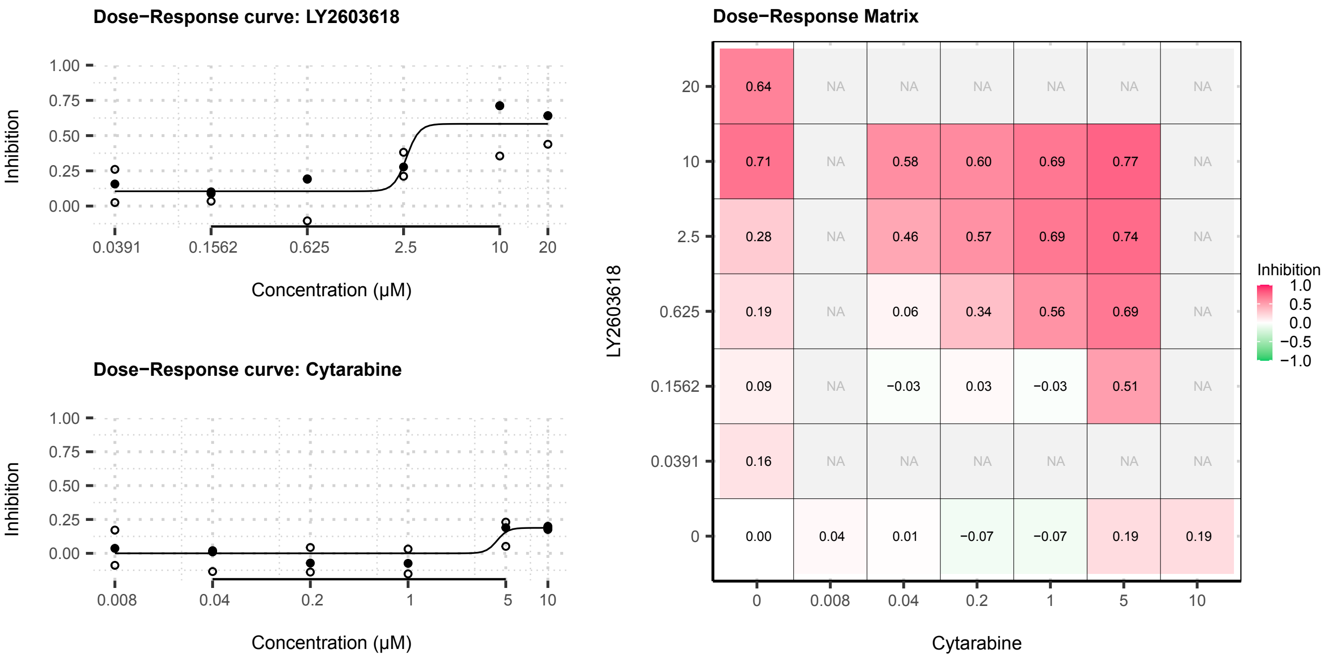
Dose−Response curve: LY2603618
Inhibition
1.00
0.75
0.50
0.25
0.00
0.0391
0.1562
0.625
2.5
10
20
Concentration (μM)
Dose−Response curve: Cytarabine
Inhibition
1.00
0.75
0.50
0.25
0.00
0.008
0.04
0.2
1
5
10
Concentration (μM)
Dose−Response Matrix
LY2603618
20
10
2.5
0.625
0.1562
0.0391
0
0.64 NA NA NA NA NA NA
0.71 NA 0.58 0.60 0.69 0.77 NA
0.28 NA 0.46 0.57 0.69 0.74 NA
0.19 NA 0.06 0.34 0.56 0.69 NA
0.09 NA −0.03 0.03 −0.03 0.51 NA
0.16 NA NA NA NA NA NA
0.00 0.04 0.01 −0.07 −0.07 0.19 0.19
0
0.008
0.04
0.2
1
5
10
Cytarabine
Inhibition
1.0
0.5
0.0
−0.5
−1.0

#### MeWo: Dose−Response Curves

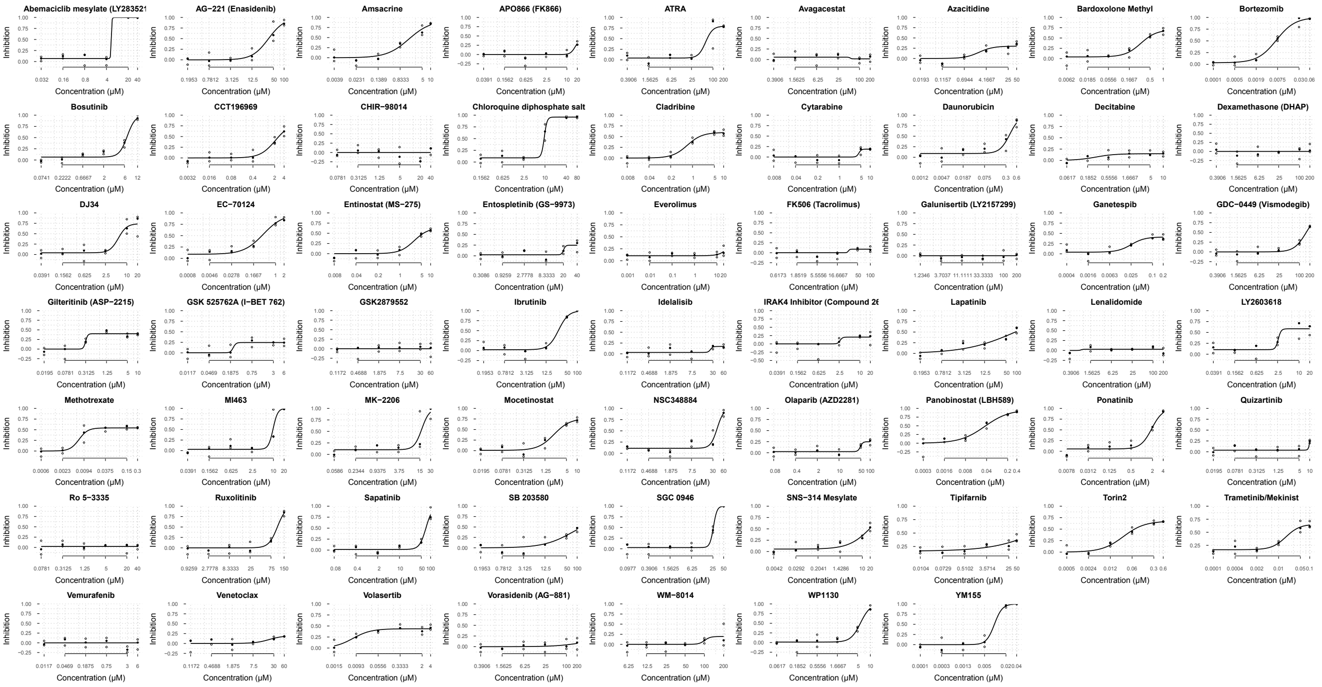

#### Cytarabine: Dose−Response Curves

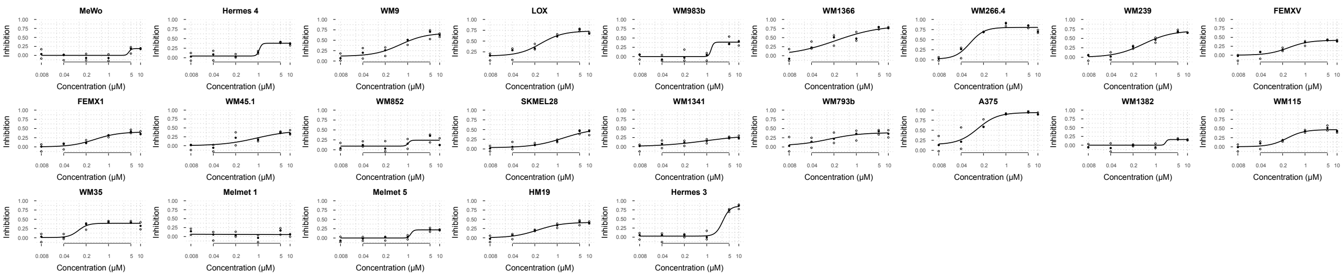

### MeWo: Dynamic Drug Activity Range

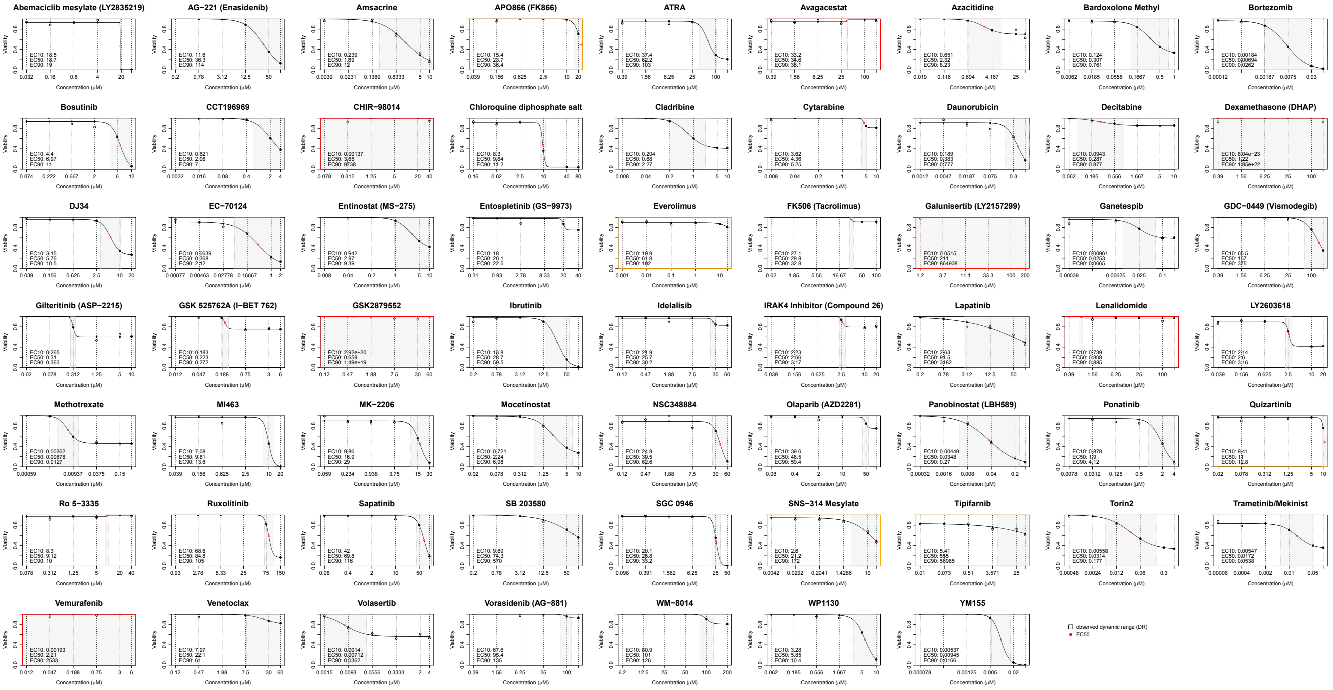

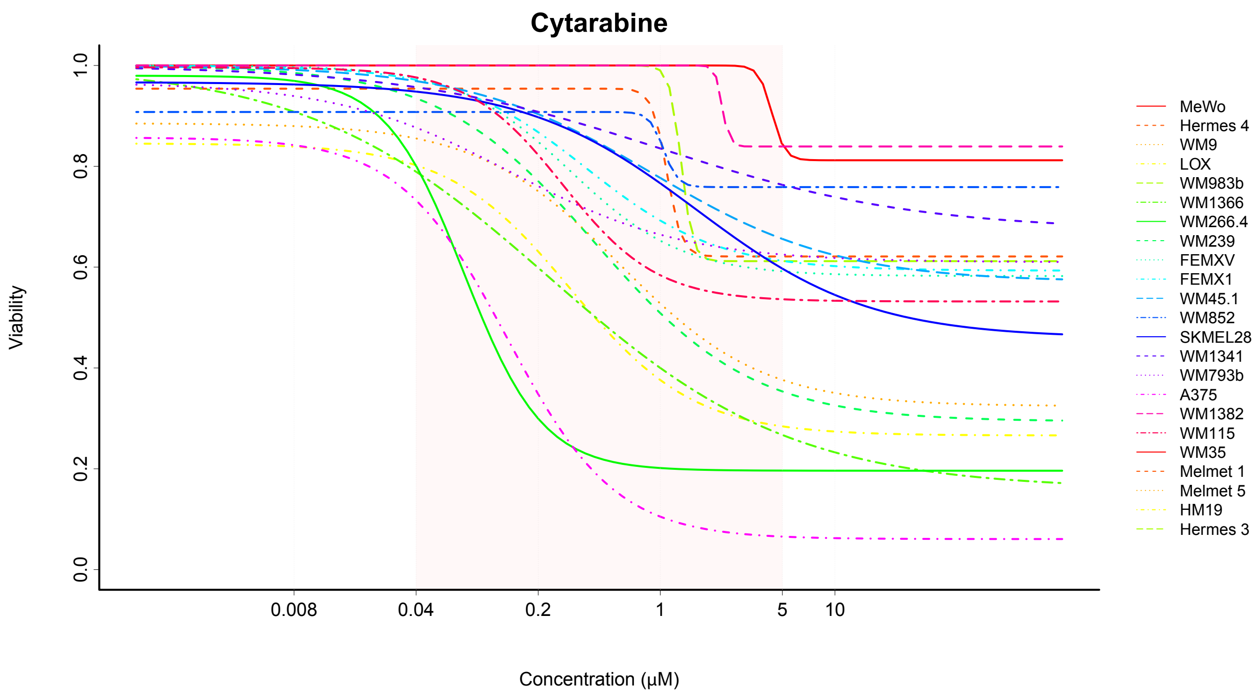
Cytarabine
Viability
Concentration (μM)
0.0
0.2
0.4
0.6
0.8
1.0
0.008
0.04
0.2
1
5
10
MeWo
Hermes 4
WM9
LOX
WM983b
WM1366
WM266.4
WM239
FEMXV
FEMX1
WM45.1
WM852
SKMEL28
WM1341
WM793b
A375
WM1382
WM115
WM35
Melmet 1
Melmet 5
HM19
Hermes 3

### Dynamic Drug Activity Range (all cell lines)

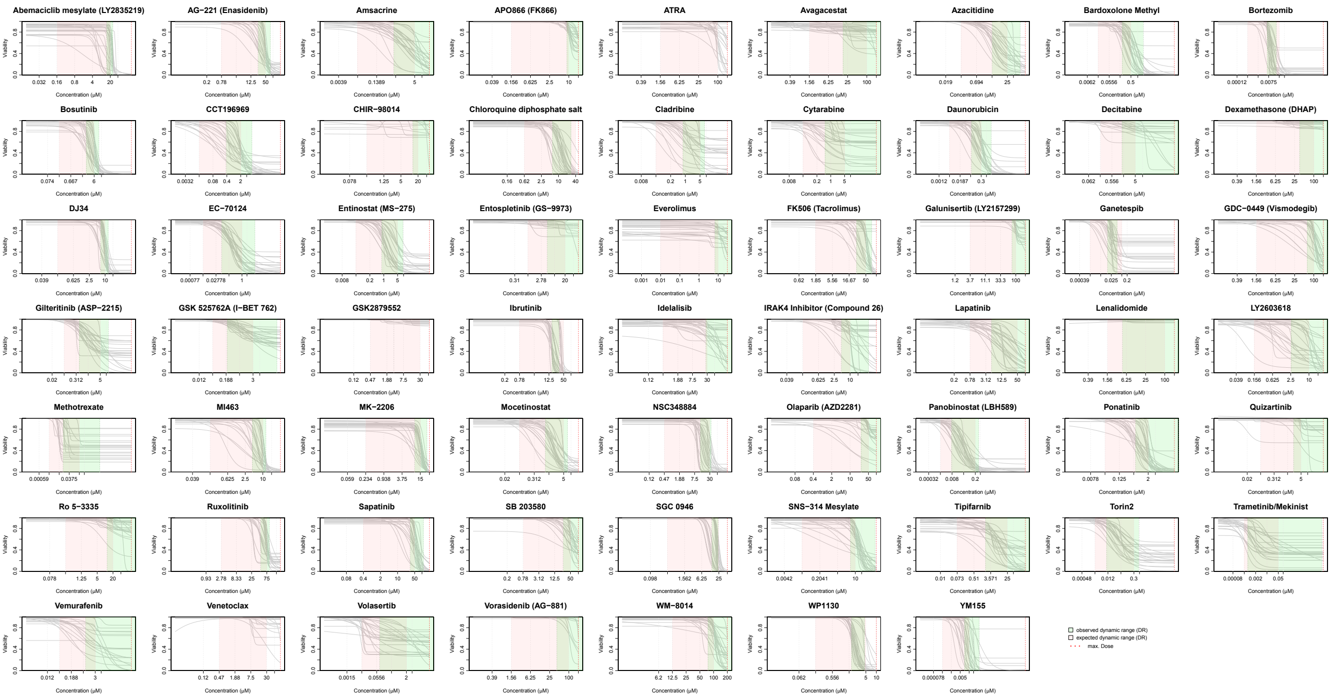

#### Dynamic Drug Activity Range (all cell lines)

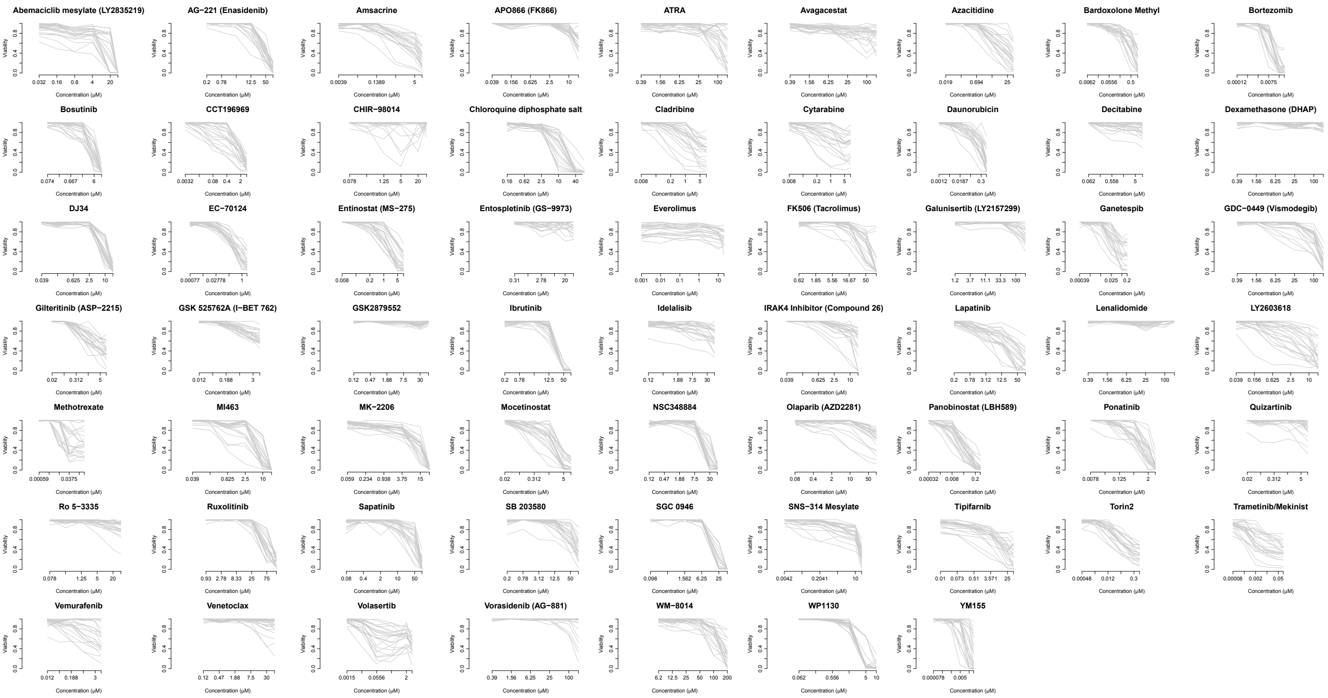

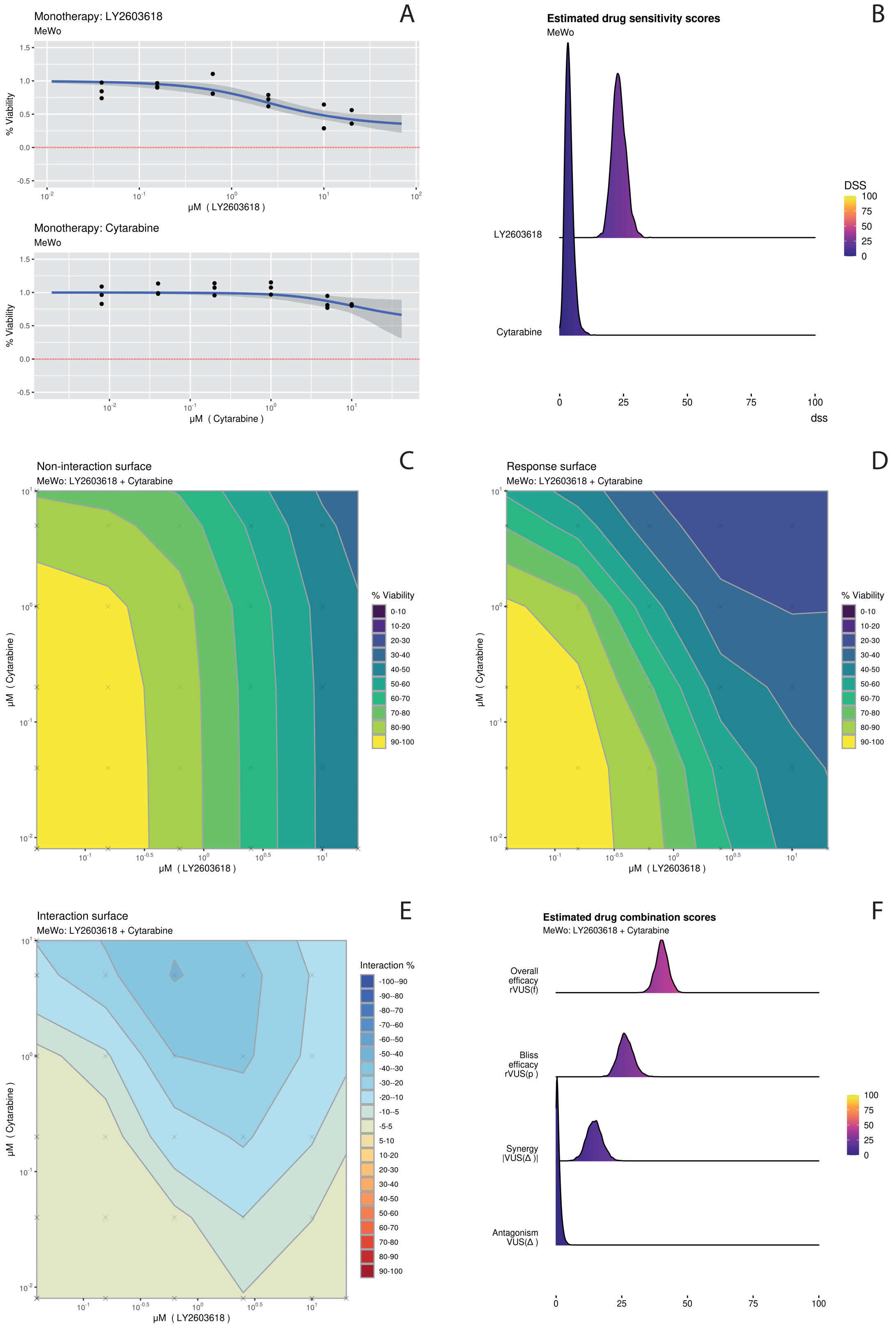
A
Monotherapy: LY2603618
MeWo
% Viability
μM ( LY2603618 )
Monotherapy: Cytarabine
MeWo
% Viability
μM ( Cytarabine )
B
Estimated drug sensitivity scores
MeWo
LY2603618
Cytarabine
DSS
dss
C
Non-interaction surface
MeWo: LY2603618 + Cytarabine
μM ( Cytarabine )
μM ( LY2603618 )
% Viability
D
Response surface
MeWo: LY2603618 + Cytarabine
μM ( Cytarabine )
μM ( LY2603618 )
% Viability
E
Interaction surface
MeWo: LY2603618 + Cytarabine
μM ( Cytarabine )
μM ( LY2603618 )
Interaction %
F
Estimated drug combination scores
MeWo: LY2603618 + Cytarabine
Overall efficacy rVUS(f)
Bliss efficacy rVUS(p )
Synergy |VUS(Δ )|
Antagonism VUS(Δ )

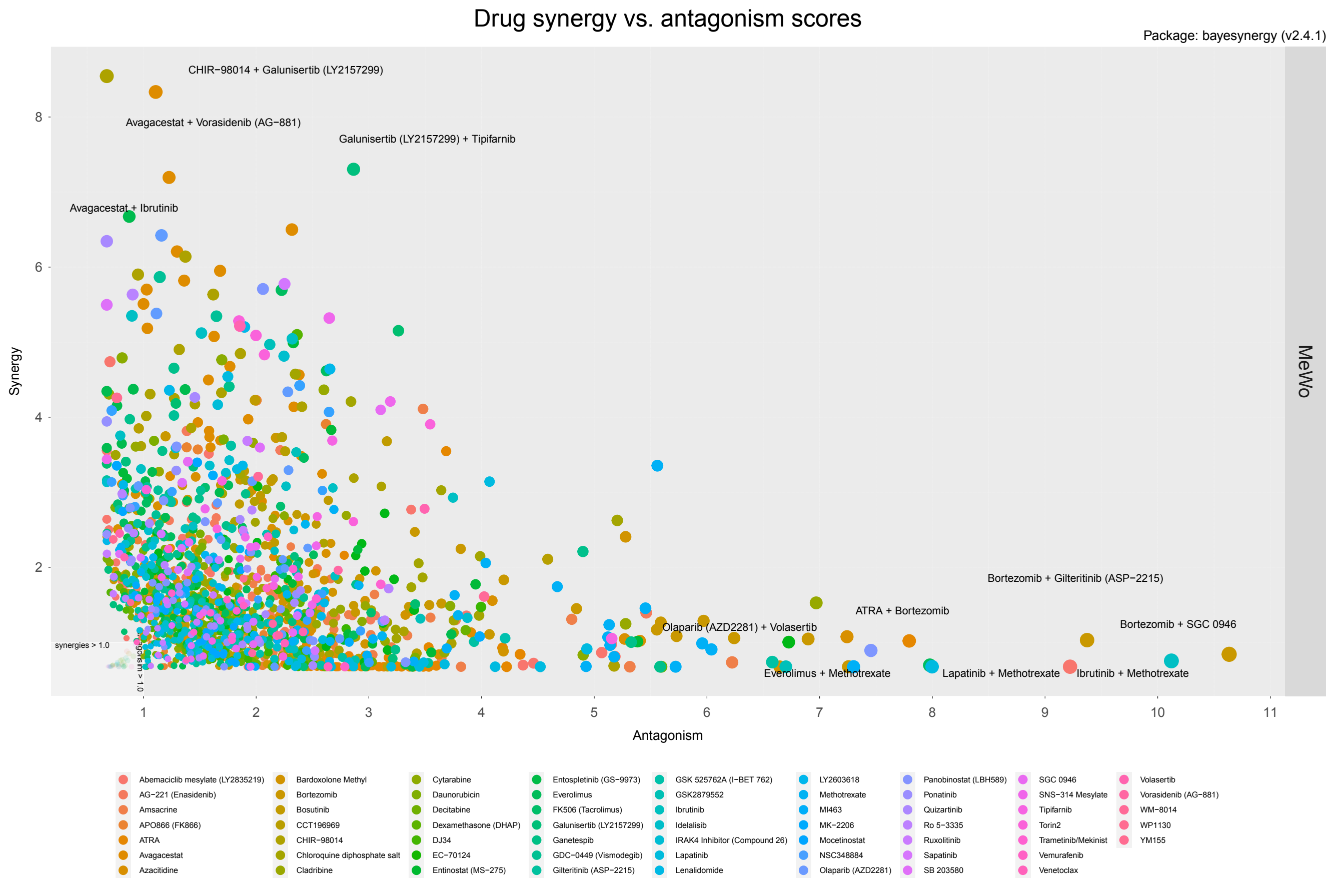

Drug synergy vs. antagonism scores
Package: bayesynergy (v2.4.1)
MeWo
Synergy
Antagonism
CHIR−98014 + Galunisertib (LY2157299)
Avagacestat + Vorasidenib (AG−881)
Galunisertib (LY2157299) + Tipifarnib
Avagacestat + Ibrutinib
Bortezomib + Gilteritinib (ASP−2215)
ATRA + Bortezomib
Bortezomib + SGC 0946
Olaparib (AZD2281) + Volasertib
Everolimus + Methotrexate
Lapatinib + Methotrexate
Ibrutinib + Methotrexate
synergies > 1.0
8
6
4
2
1
2
3
4
5
6
7
8
9
10
11
Abemaciclib mesylate (LY2835219)
AG−221 (Enasidenib)
Amsacrine
APO866 (FK866)
ATRA
Avagacestat
Azacitidine
Bardoxolone Methyl
Bortezomib
Bosutinib
CCT196969
CHIR−98014
Chloroquine diphosphate salt
Cladribine
Cytarabine
Daunorubicin
Decitabine
Dexamethasone (DHAP)
DJ34
EC−70124
Entinostat (MS−275)
Entospletinib (GS−9973)
Everolimus
FK506 (Tacrolimus)
Galunisertib (LY2157299)
Ganetespib
GDC−0449 (Vismodegib)
Gilteritinib (ASP−2215)
GSK 525762A (I−BET 762)
GSK2879552
Ibrutinib
Idelalisib
IRAK4 Inhibitor (Compound 26)
Lapatinib
Lenalidomide
LY2603618
Methotrexate
MI463
MK−2206
Mocetinostat
NSC348884
Olaparib (AZD2281)
Panobinostat (LBH589)
Ponatinib
Quizartinib
Ro 5−3335
Ruxolitinib
Sapatinib
SB 203580
SGC 0946
SNS−314 Mesylate
Tipifarnib
Torin2
Trametinib/Mekinist
Vemurafenib
Venetoclax
Volasertib
Vorasidenib (AG−881)
WM−8014
WP1130
YM155

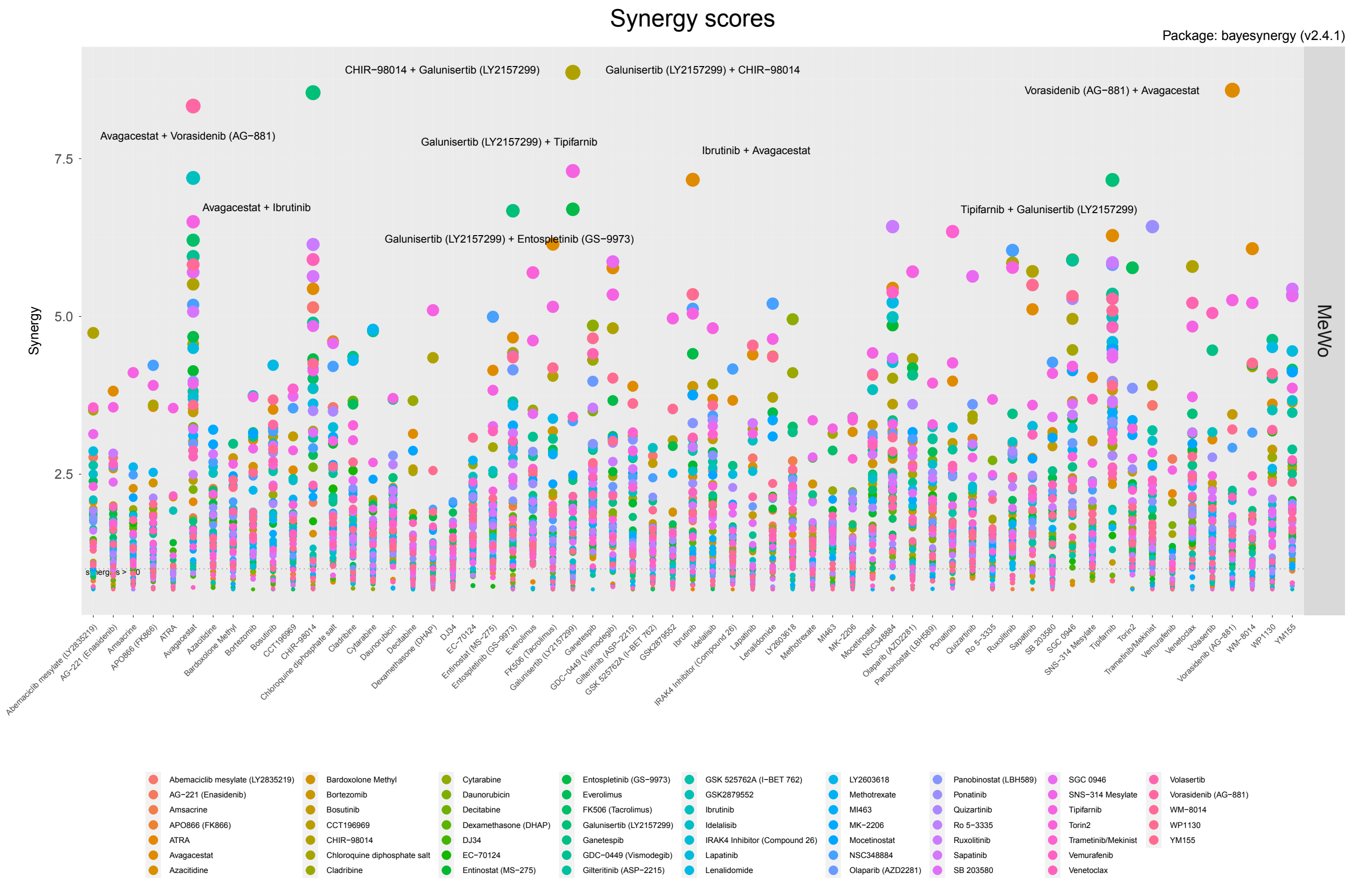
Synergy scores
Package: bayesynergy (v2.4.1)
CHIR−98014 + Galunisertib (LY2157299)
Galunisertib (LY2157299) + CHIR−98014
Vorasidenib (AG−881) + Avagacestat
Avagacestat + Vorasidenib (AG−881)
Galunisertib (LY2157299) + Tipifarnib
Ibrutinib + Avagacestat
Avagacestat + Ibrutinib
Tipifarnib + Galunisertib (LY2157299)
Galunisertib (LY2157299) + Entospletinib (GS−9973)
Synergy
7.5
5.0
2.5
MeWo
Abemaciclib mesylate (LY2835219)
AG−221 (Enasidenib)
Amsacrine
APO866 (FK866)
ATRA
Avagacestat
Azacitidine
Bardoxolone Methyl
Bortezomib
Bosutinib
CCT196969
CHIR−98014
Chloroquine diphosphate salt
Cladribine
Cytarabine
Daunorubicin
Decitabine
Dexamethasone (DHAP)
DJ34
EC−70124
Entinostat (MS−275)
Entospletinib (GS−9973)
Everolimus
FK506 (Tacrolimus)
Galunisertib (LY2157299)
Ganetespib
GDC−0449 (Vismodegib)
Gilteritinib (ASP−2215)
GSK 525762A (I−BET 762)
GSK2879552
Ibrutinib
Idelalisib
IRAK4 Inhibitor (Compound 26)
Lapatinib
Lenalidomide
LY2603618
Methotrexate
MI463
MK−2206
Mocetinostat
NSC348884
Olaparib (AZD2281)
Panobinostat (LBH589)
Ponatinib
Quizartinib
Ro 5−3335
Ruxolitinib
Sapatinib
SB 203580
SGC 0946
SNS−314 Mesylate
Tipifarnib
Torin2
Trametinib/Mekinist
Vemurafenib
Venetoclax
Volasertib
Vorasidenib (AG−881)
WM−8014
WP1130
YM155

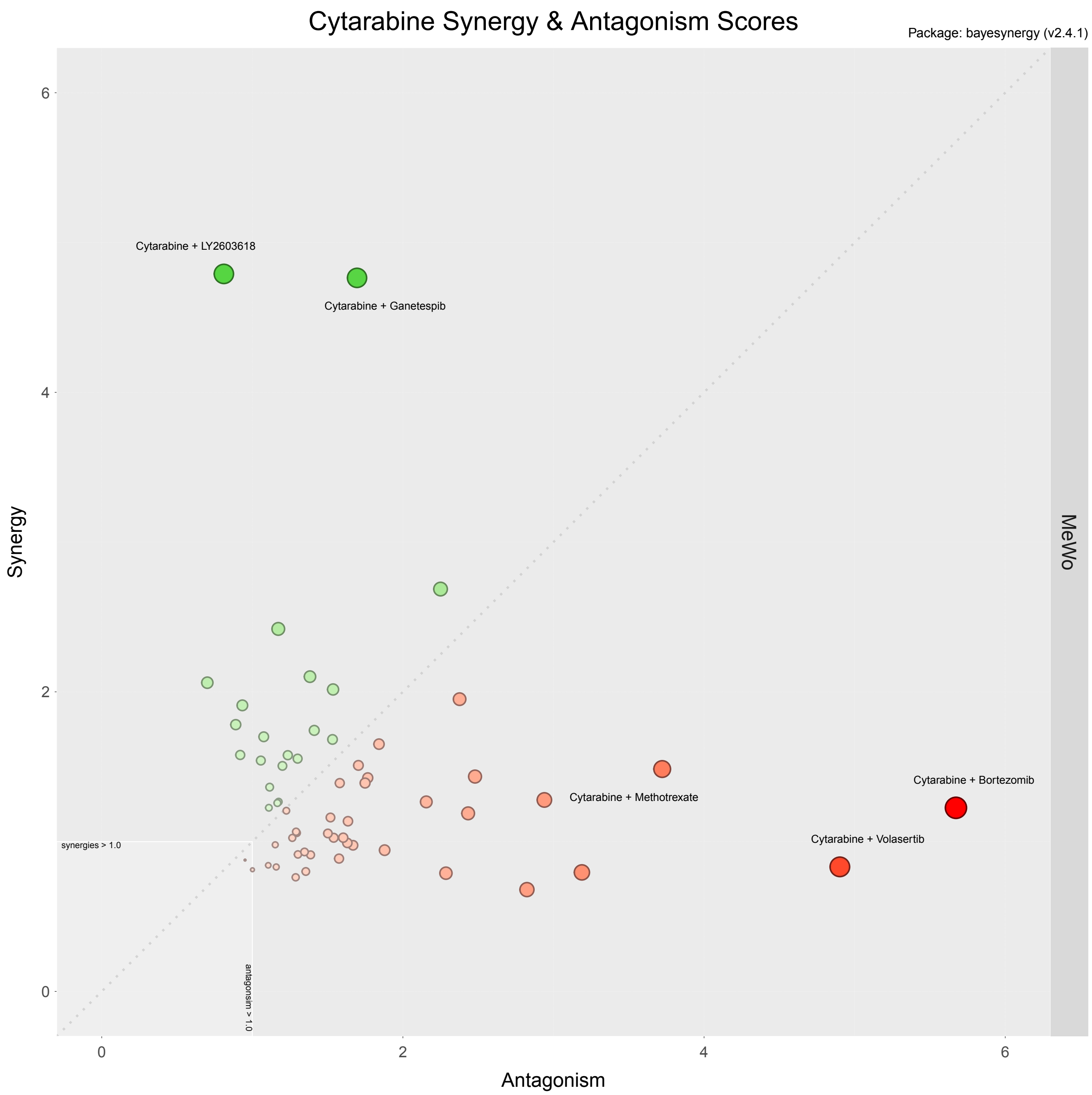
Cytarabine Synergy & Antagonism Scores
Package: bayesynergy (v2.4.1)
Synergy
Antagonism
MeWo
Cytarabine + LY2603618
Cytarabine + Ganetespib
Cytarabine + Methotrexate
Cytarabine + Bortezomib
Cytarabine + Volasertib
synergies > 1.0
antagonsim > 1.0
0
2
4
6
